# Automated AI labelling of optic nerve head enables new insights into cross-ancestry glaucoma risk and genetic discovery in over 280,000 images from the UK Biobank and Canadian Longitudinal Study on Aging

**DOI:** 10.1101/2020.11.03.367623

**Authors:** Xikun Han, Kaiah Steven, Ayub Qassim, Henry N Marshall, Cameron Bean, Michael Tremeer, Jiyuan An, Owen Siggs, Puya Gharahkhani, Jamie E Craig, Alex W Hewitt, Maciej Trzaskowski, Stuart MacGregor

**Affiliations:** QIMR Berghofer Medical Research Institute, Brisbane, Australia; School of Medicine, University of Queensland, St Lucia, Brisbane, Australia; Max Kelsen, Brisbane, Australia; Department of Ophthalmology, Flinders University, Flinders Medical Centre, Australia; Queensland University of Technology, Brisbane, Australia; Menzies Institute for Medical Research, University of Tasmania, Australia; Centre for Eye Research Australia, University of Melbourne, Australia

**Keywords:** artificial intelligence, image, optic nerve head, glaucoma, GWAS, UK Biobank, CLSA

## Abstract

Cupping of the optic nerve head, a highly heritable trait, is a hallmark of glaucomatous optic neuropathy. Two key parameters are vertical cup-to-disc ratio (VCDR) and vertical disc diameter (VDD). However, manual assessment often suffers from poor accuracy and is time-intensive. Here, we show convolutional neural network models can accurately estimate VCDR and VDD for 282,100 images from both UK Biobank and an independent study (Canadian Longitudinal Study on Aging), enabling cross-ancestry epidemiological studies and new genetic discovery for these optic nerve head parameters. Using the AI approach we perform a systematic comparison of the distribution of VCDR and VDD, and compare these with intraocular pressure and glaucoma diagnoses across various genetically determined ancestries, which provides an explanation for the high rates of normal tension glaucoma in East Asia. We then used the large number of AI gradings to conduct a more powerful genome-wide association study (GWAS) of optic nerve head parameters. Using the AI based gradings increased estimates of heritability by ~50% for VCDR and VDD. Our GWAS identified more than 200 loci for both VCDR and VDD (double the number of loci from previous studies), uncovers dozens of novel biological pathways, with many of the novel loci also conferring risk for glaucoma.

## Introduction

The optic nerve head is the exit point of retinal ganglion cell axons from the eye to the brain.^1^ It is commonly assessed during ophthalmic examinations using fundoscopy or optical imaging technology for multiple ocular diseases, such as glaucoma, which is the leading cause of irreversible blindness globally and is characterized by characteristic cupping of the optic disc as a result of retinal ganglion cell apoptosis.^2,3^ Enlarged vertical cup-to-disc ratio (VCDR) is considered a hallmark of glaucomatous optic neuropathy and is often used to define glaucoma in general population based prevalence surveys.^4^ However, VCDR alone is not adequate to assess glaucomatous damage in part because of the variation of optic disc size. For instance, a vertical cup:disc ratio of 0.5 in a small optic disc could be pathologic whereas a vertical cup:disc ratio of 0.8 in a large disc size may represent physiologic cupping. Adjusting for optic disc size is hence important to maximizing the clinical utility of VCDR in diagnosing glaucoma.

Family studies have shown that optic disc morphology traits are highly heritable with an estimated heritability of 0.48 and 0.57 for VCDR and optic disc diameter, respectively.^5^ Large-scale genome-wide association studies (GWAS) for optic disc morphology have begun to shed light on the development and pathogenesis of glaucoma and other optic nerve diseases.^6–8^ However, both large sample sizes and accurate phenotyping are critical in GWAS and further progress will be difficult under the existing manual phenotype paradigm. Manual assessment of optic disc photographs is time-intensive and often suffers from poor inter-observer concordance, even when performed by trained specialists and an alternative approach is required.^9,10^ Clinical estimates of VCDR are more difficult from monoscopic photographs compared with stereoscopic viewing of the optic nerve head which can be achieved during slit-lamp biomicroscopy or from stereoscopic photographs.

Recent advances in artificial intelligence (AI) algorithms have shown exciting promise in healthcare^11^, including the automated diagnosis of eye diseases.^12,13^ With the high performance of AI technology, the U.S. Food and Drug Administration approved the first medical device to use AI technology to detect diabetic retinopathy in 2018.^14,15^ The probabilistic nature and non-linear capabilities, as well as analytical capabilities to deal with single and multimodal, high-dimensional data, has seen application of AI experience lower resistance to adoption in the medical field when applied to computer vision applications. Two fundamental properties have facilitated AI application to medical diagnostics. Firstly, the problem space (medical imaging) is, relative to other medical domains, well studied and very well understood. Secondly, an observation of the output can be quickly validated by a clinical practitioner, who by having access to additional clinical or historical data about that patient, may suggest alternative diagnosis. A motivating factor driving utilisation of AI on data such as fundus images is the large volume of images available for algorithms to be trained on. Furthermore, standardised imaging techniques can drastically reduce the dataset heterogeneity. This is highlighted by the collection of images as part of the UKB and CLSA biobanks completed over a decade. Automated diagnosis from retinal fundus imaging has been approached through a number of different algorithms, ranging from multi-stage “classical” learning algorithms to end-to-end deep learning models.^16–19^

In this study, a convolutional neural network (CNN) model was utilised in a transfer learning approach, training on clinical assessments of the optic nerve head in ~70,000 photographs (Labelled Training Data) of UK Biobank (UKB) participants. Automatic labelling by the CNN model dramatically boosts the effective sample size (n=282,100 total images graded), presenting an opportunity to greatly expand the utility of the GWAS approach for VCDR and optic disc diameter. We also apply the AI labels systematically across the multiple different ancestries in UKB and CLSA and investigate how VCDR and other glaucoma risk factors, such as IOP, relate to glaucoma risk in different ancestries.

## Results

### Study Design And Overview

The overall study design is summarised in Figure 1. We use transfer learning to train three CNN models for image gradability, VCDR, and vertical disc diameter (VDD) values from ~70,000 UKB fundus images graded by clinicians. These models were then applied to all UKB fundus images (85,736 participants and 175,770 images in total) and another independent cohort - the Canadian Longitudinal Study on Aging (CLSA, 29,635 participants and 106,330 images in total). We performed the largest AI-based GWAS for VCDR and VDD, and replicated novel genetic discoveries in clinician-graded fundus images from International Glaucoma Genetics Consortium (IGGC) and in glaucoma case-control studies (UKB and the Australian and New Zealand Registry of Advanced Glaucoma; ANZRAG). The large scale biobank data for both VCDR and IOP also allow us to systematically compare the glaucoma risk and optic nerve head parameters across different ancestries.

**Figure 1.**
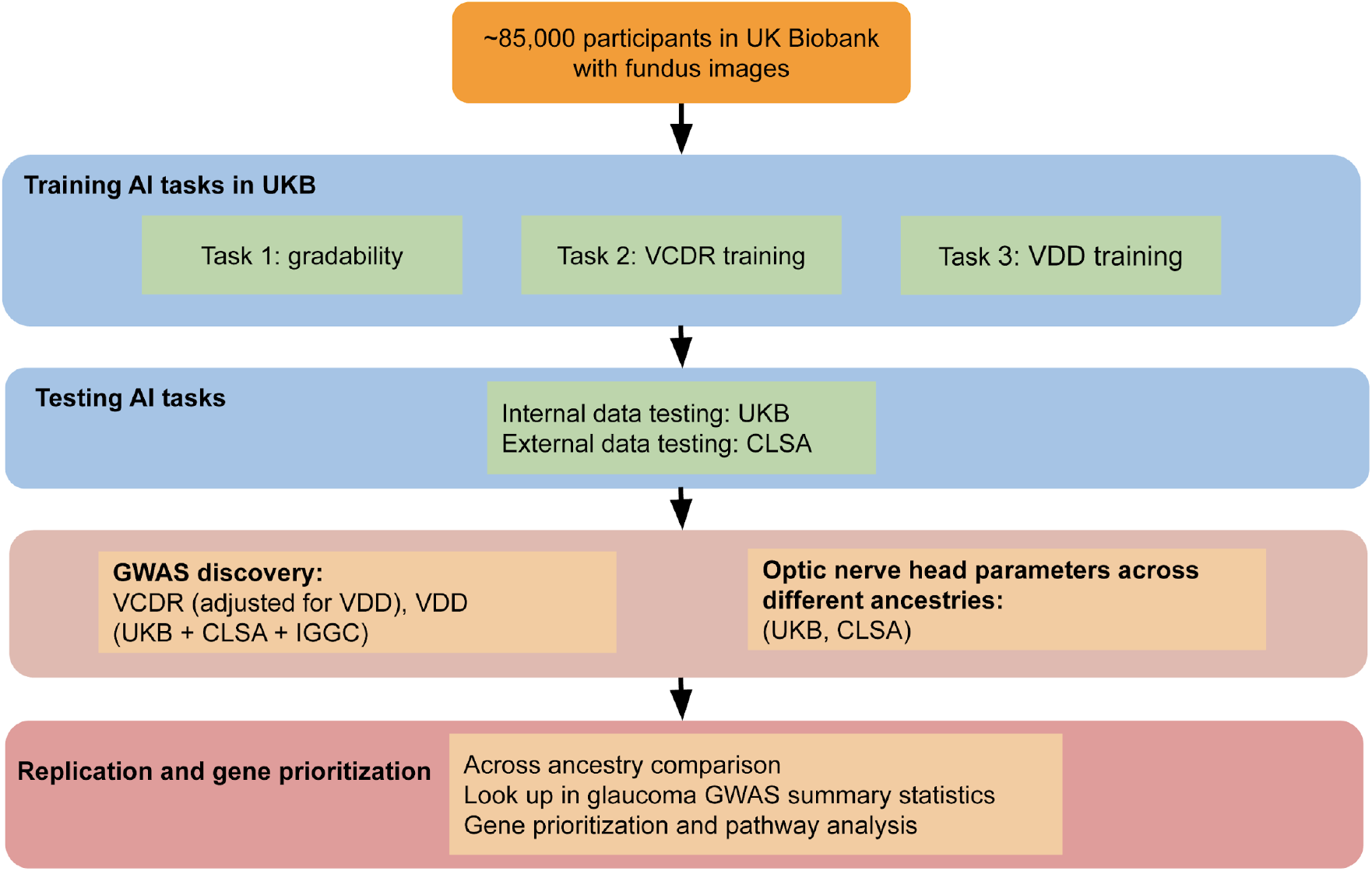
Flowchart of AI framework and datasets. In UK Biobank (UKB), the fundus retinal eye images were available for ~85,000 participants (~68,000 participants in the baseline visit and ~19,000 participants in the first repeat assessment visit). In our previous study, vertical cup-to-disc ratio (VCDR) and vertical disc diameter (VDD) were graded by two clinicians in ~70,000 photographs using a custom Java program. These clinical assessments were used as Training Data for three convolutional neural network (CNN) models for image gradability, VCDR, and VDD values. The learned models were then applied to all UKB fundus images (85,736 participants and 175,770 images in total) and another independent cohort - the Canadian Longitudinal Study on Aging (CLSA, 29,635 participants and 106,330 images in total). The AI labels were further used to systematically evaluate optic nerve head parameters across the multiple different ancestries in UKB and CLSA, and allowed us to perform the largest AI-based GWAS for VCDR and VDD.

### Study data and performance of the trained AI model

In the UKB, 85,736 participants had at least one fundus retinal image, with a total of 175,770 images available (Table 1). The mean age at baseline was 57.0 (SD: 8.1) years and 54% were women. In the CLSA cohort, 29,635 participants with 106,330 images were included in analysis, of whom 50% were women, and the mean age at recruitment was 62.6 (SD: 10.0) years.

**Table 1.** Characteristics of retinal fundus images from the UK Biobank and Canadian Longitudinal Study on Aging participants.

| Variable |  | UKB | CLSA |
| --- | --- | --- | --- |
| Number of images |  | 175,770 | 106,330 |
| Number of participants |  | 85,736 | 29,635 |
| % with at least one gradable image |  | 95% | 99% |
| Sex | Women (%) | 44,017 (54%) | 14,941 (51%) |
| Age at recruitment | Mean (SD), years | 57 ± 8 | 63 ± 10 |
| Vertical cup-disc ratio | Unit in 0-1 | 0.37 ± 0.14 | 0.35 ± 0.15 |
| Vertical disc diameter | Unit in pixel count | 129.0 ± 10.5 | 121.4 ± 10.6 |

CLSA, Canadian Longitudinal Study on Aging cohort; SD, standard deviation; UKB, UK Biobank.

We first trained a convoluted neural network to assess if each image was gradable in the UKB training sample. We found that most participants (> 95%) had gradable images in the UKB and the CLSA cohort (Supplementary Figure 1). We then predicted the measurements of both VCDR and VDD, and compared the AI-based measures with clinician gradings. The AI-based VCDR and VDD measurements exhibited a higher concordance to clinician gradings compared with previous gradings by two clinicians.^8,20–22^ For instance, the Pearson’s correlation coefficient of the VCDR measurements in the UKB samples was 0.81 (95% confidence interval [CI]: 0.80-0.81), and 0.84 (95% CI: 0.82-0.86) for an independent Canadian data set (CLSA) (Supplementary Figure 2). We therefore speculated that with the improved accuracy of VCDR and VDD measurements and the larger number of images graded, the optic nerve head assessment would increase the power for genetic discovery.

### Optic nerve head parameters and intraocular pressure across different ancestries

We compared AI model-derived VCDR and VDD measurements across different genetically-defined ancestry groups. VDD was similar across 3 ancestral groups (Europeans, East Asians and South Asians) and larger in Africans (Figure 2B, 2E). On average, after adjusting for age, sex, and VDD, VCDR was markedly higher in Asians and Africans than it was in Europeans (similar results in UKB Figure 2A and in CLSA Figure 2D). A different ancestry-based trend was also observed for intraocular pressure (IOP); relative to Europeans, South Asians had similar IOP, East Asians had lower IOP, and Africans had higher IOP (Figure 2C,F).

**Figure 2.**
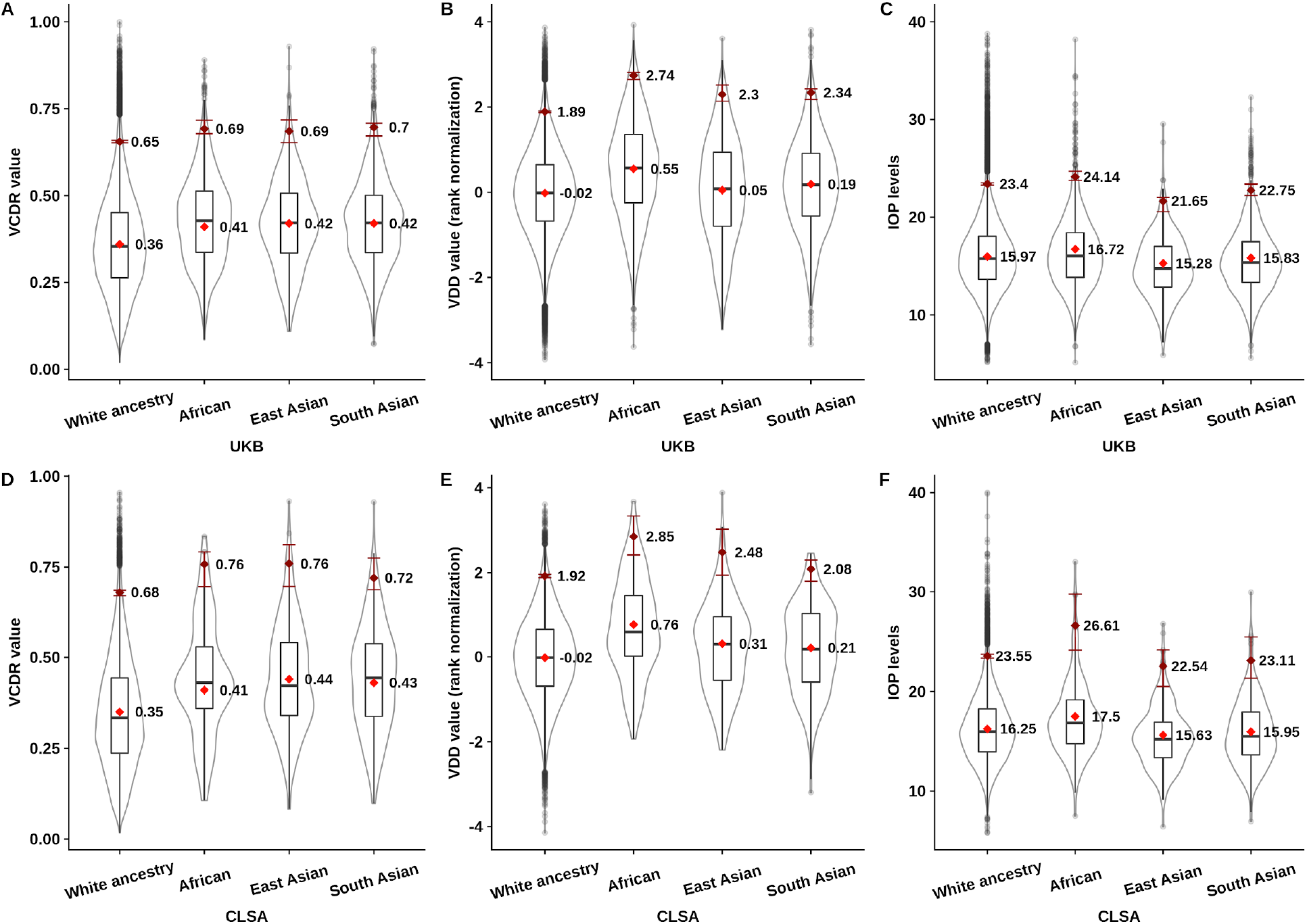
Optic nerve head measurements and intraocular pressure across different ancestry groups. Panel A shows the boxplot for VCDR values from different ancestry groups in UK Biobank. The box represents median value with first and third quartiles. The red diamond is the mean value of VCDR after accounting for age, sex, and VDD, where the mean value is annotated as text. The dark red diamond is the 97.5th percentile of VCDR value. The dark red error bar is the 95% confidence interval (2.5% to 97.5% quantiles) of the 97.5th percentile based on 1000 bootstrapped samples, which is essential for CLSA data, where the sample size for African, East Asian and South Asian was substantially smaller (N < 300). Panel B shows the boxplot for VDD values from different ancestry groups in UK Biobank. Due to the scale from fundus images, the VDD was rank normalized (mean = 0, SD = 1). The red diamond is the mean value of VDD after accounting for age and sex. Panel C shows the boxplot for IOP levels from different ancestry groups in the UK Biobank (truncated at 40 mm Hg, with 15 participants between 40 - 60 mm Hg). Panel D, E and F show the boxplots for VCDR, VDD and IOP in the CLSA cohort, respectively.

We then examined whether the systematically assessed VCDR, VDD and IOP can explain the observed prevalence of glaucoma seen across different ancestries in the UK and Canada. Figure 3 shows the glaucoma risk of Africans, East Asians and South Asians, with European ancestry (the most common ancestry in UKB and CLSA data sets) as the baseline. Consistent with previous epidemiological studies, Africans have the highest glaucoma risk (Figure 3 base model, correcting for only age and sex OR = 2.5 relative to the reference of Europeans). As seen in Figure 2, Africans have higher VCDR and higher IOP than Europeans and when these were corrected for, the glaucoma risk approached that of Europeans in both CLSA and UKB. East Asians had a similar base model risk to Europeans, although the contribution of IOP and VDR differs; on average their IOP is lower and their VCDR is larger (Figure 2), with the pattern of glaucoma risk changing as either IOP alone or VCDR alone were adjusted for in the regression model. Adjusting for both IOP and VCDR, the risk of glaucoma in East Asians was not significantly different to Europeans, suggesting that the higher VCDR and lower IOP in this group relative to Europeans counteract each other, explaining the similar glaucoma incidences between these ancestries. Interestingly, in South Asians, IOP is similar to Europeans, but VCDR is higher (Figure 2). This means that South Asian base model risk does not change when IOP is included in the model, but when VCDR is included the glaucoma risk decreases to become indistinguishable from the incidence in Europeans. In summary, by examining individuals of varying ancestry living in the UK and Canada, we show that relative to European ancestry, African ancestry glaucoma incidence is driven by both elevated VCDR and IOP, East Asian ancestry glaucoma is driven by elevated VCDR but ameliorated by lower IOP and finally that South Asian glaucoma is driven by elevated VCDR, but not by changes in IOP (relative to that in Europeans).

**Figure 3.**
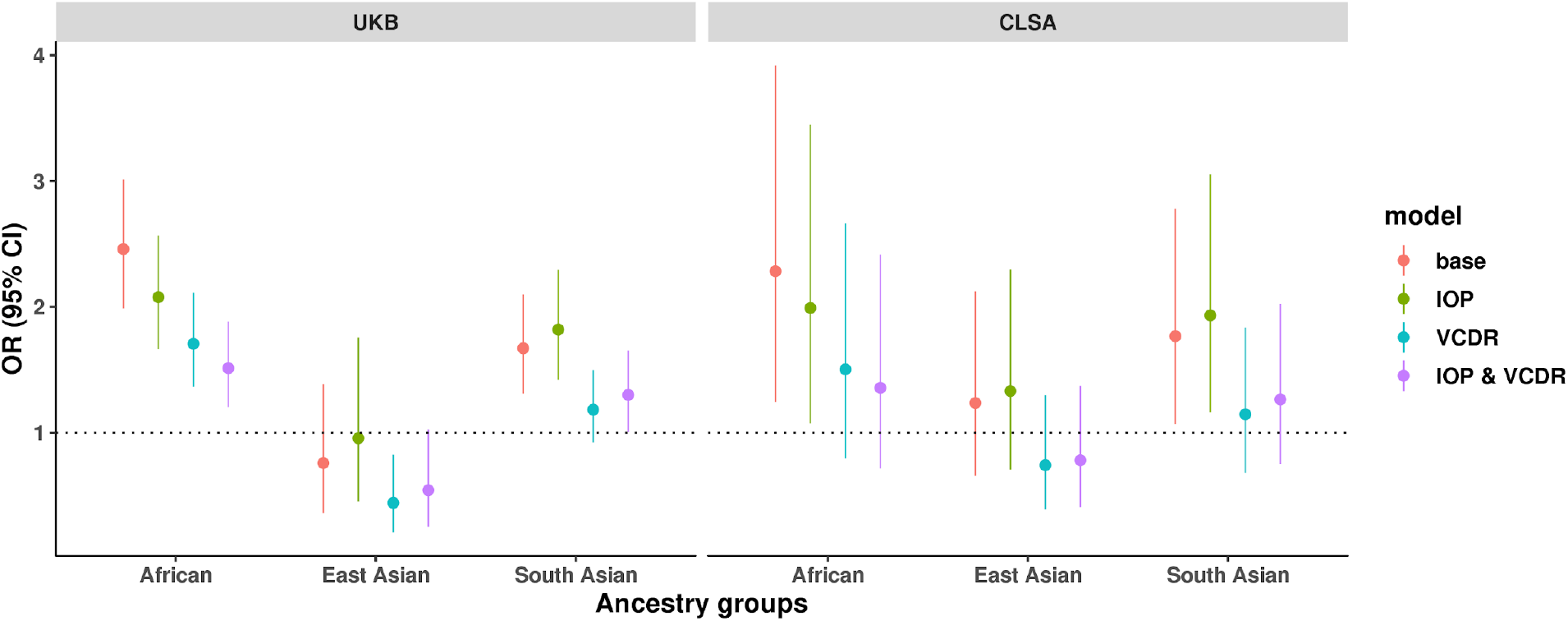
Glaucoma risk across different ancestry groups. The figure shows the risk of glaucoma in different ancestry groups. The horizontal line at OR = 1 is the reference for European ancestry. The Y-axis is the odds ratio (OR) and 95% confidence interval (CI) for three ethnic groups (African, South Asian, and East Asian). In each different model, different covariates were adjusted to evaluate the association of ethnic groups and glaucoma risk. In the base model, only sex and age were adjusted for; the other models also include either IOP, VCDR, or both (IOP & VCDR).

### AI-based phenotypes greatly increase SNP-based heritability and identify more loci

In the GWAS of VDD-adjusted VCDR, 145 and 19 statistically independent genome-wide significant SNPs were respectively identified in the UKB alone and CLSA alone (Supplementary Figure 3). The analogous numbers of SNPs for VDD were 142 and 17 for UKB and CLSA, respectively. We found weak evidence of genomic inflation from linkage disequilibrium score regression (Supplementary Table 1). From UKB, the AI-based GWAS of VDD-adjusted VCDR and VDD identified substantially more loci than our previous GWAS based on clinician gradings (76 for VDD-adjusted VCDR and 91 for VDD)^8,20^. Strikingly, the SNP-based heritability increased by ~50% for VCDR and VDD (Supplementary Figure 4). For instance, the SNP-based heritability for VCDR was 0.22 from clinician gradings (only single measure), whereas the heritability increased to 0.35 from AI-based GWAS (average of multiple measures). The increased heritability indicated that AI-based phenotyping was substantially cleaner than clinician gradings, which may be a result of two aspects: 1) higher accuracy of AI-based gradings; 2) improved accuracy from multiple measures per individual. We further tested the hypothesis in UKB and CLSA using only one measure per individual from AI-based gradings. The SNP-based heritability from a single measure (left or right eyes in the baseline or first follow-up visit) was ~0.3, which is roughly in the middle of heritability estimation from clinician gradings and AI-based multiple measures (Supplementary Figure 4). These results indicate the higher accuracy of AI-based single measure per individual contributes to the increase of heritability estimation, and averaging of multiple measures per individual can further increase the heritability. Consistent with our previous study, correcting for VDD in VCDR GWAS also improved the relevance to glaucoma, with a higher genetic correlation with glaucoma in VDD-adjusted VCDR compared with unadjusted VCDR GWAS (genetic correlation rg = 0.502 vs 0.457 in UKB, and 0.543 vs 0.481 in CLSA).

### Validation AI-based GWAS

We then compared AI-based and clinician grading-based GWAS using independent samples from the IGGC. The concordance of SNP effect sizes of top SNPs between the AI-based and clinician gradings was essentially one (Panel A and D in Figure 4), and nearly all previously published loci using clinician ratings were replicated. The estimated effect sizes at the top SNPs from AI-based GWAS were also highly concordant between UKB and CLSA (Panel B and E in Figure 4). When combining UKB and CLSA AI-based GWAS we identified 193 and 188 loci for VDD-adjusted VCDR and VDD, respectively, again exhibiting very high concordance with IGGC (Panel C and F in Figure 4). The high concordance and more loci support the better-powered GWAS from AI-based measurements.

**Figure 4.**
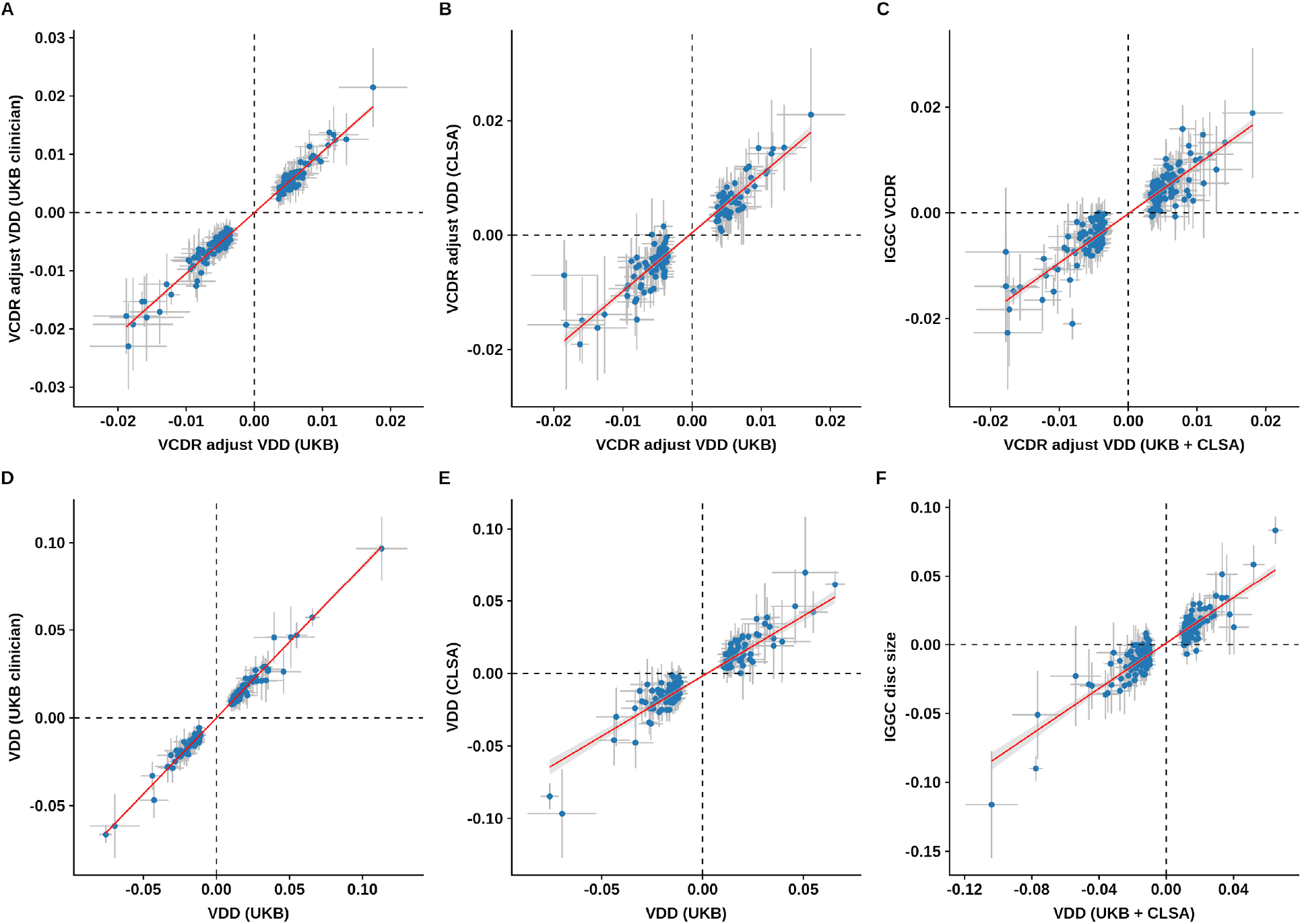
Validation AI-based GWAS. The figure shows the effect sizes for VDD-adjusted VCDR and VDD from different data sets. The vertical and horizontal error bars are the 95% confidence interval for SNP effect sizes. The red line is the best fit line with 95% confidence interval region in grey.

### New genetic discovery of optic nerve head measures, cross-ancestry comparison, and implications for glaucoma

To maximize power for locus discovery, we combined UKB, CLSA and IGGC GWAS (European ancestry), and identified 230 and 231 independent genome-wide significant SNPs for VDD-adjusted VCDR and VDD, respectively (Figure 5). Of them, we found 111 and 107 novel loci for VDD-adjusted VCDR and VDD, respectively (Supplementary Table 2 and 3). We then compared the effect sizes of top VDD-adjusted VCDR and VDD loci across different ancestries (Asian and African GWAS), due to the much smaller available sample sizes, their confidence intervals of effect estimations were very large, however the clear linear trend indicated the loci identified from European ancestry also had an effect on Asian populations (Figure 6A, B, for VCDR and VDD the Pearson’s correlation coefficient is 0.65 [P value 3.6 × 10^−27^] and 0.62 [P value 9.3 × 10^−23^], respectively). The sample size of African ancestry was much smaller than Asian ancestry (N = 2,245 versus 8,373 for VCDR) and showed a lower concordance (Supplementary Figure 5). The genetic correlations across the genome were essentially one based on the Popcorn approach for VCDR and VDD (Supplementary Table 4). We also compared the effect sizes of VDD-adjusted VCDR top loci with their effect sizes on glaucoma (Figure 6C), and found a relatively high concordance (Pearson’s correlation coefficient 0.61, P = 8.2 × 10^−25^). Of the 230 VCDR (adjusted for VDD) loci (227 available in glaucoma GWAS), 187 (82%) were in the same direction, 84 were associated with glaucoma at a nominal significance level (P<0.05) and 24 were associated with glaucoma after Bonferroni correction (P< 0.05/227= 2.2 × 10^−4^, the nearest gene names are highlighted in Figure 6C, *e.g. LMX1B, ABCA1, CAV1*, and *GAS7*).

**Figure 5.**
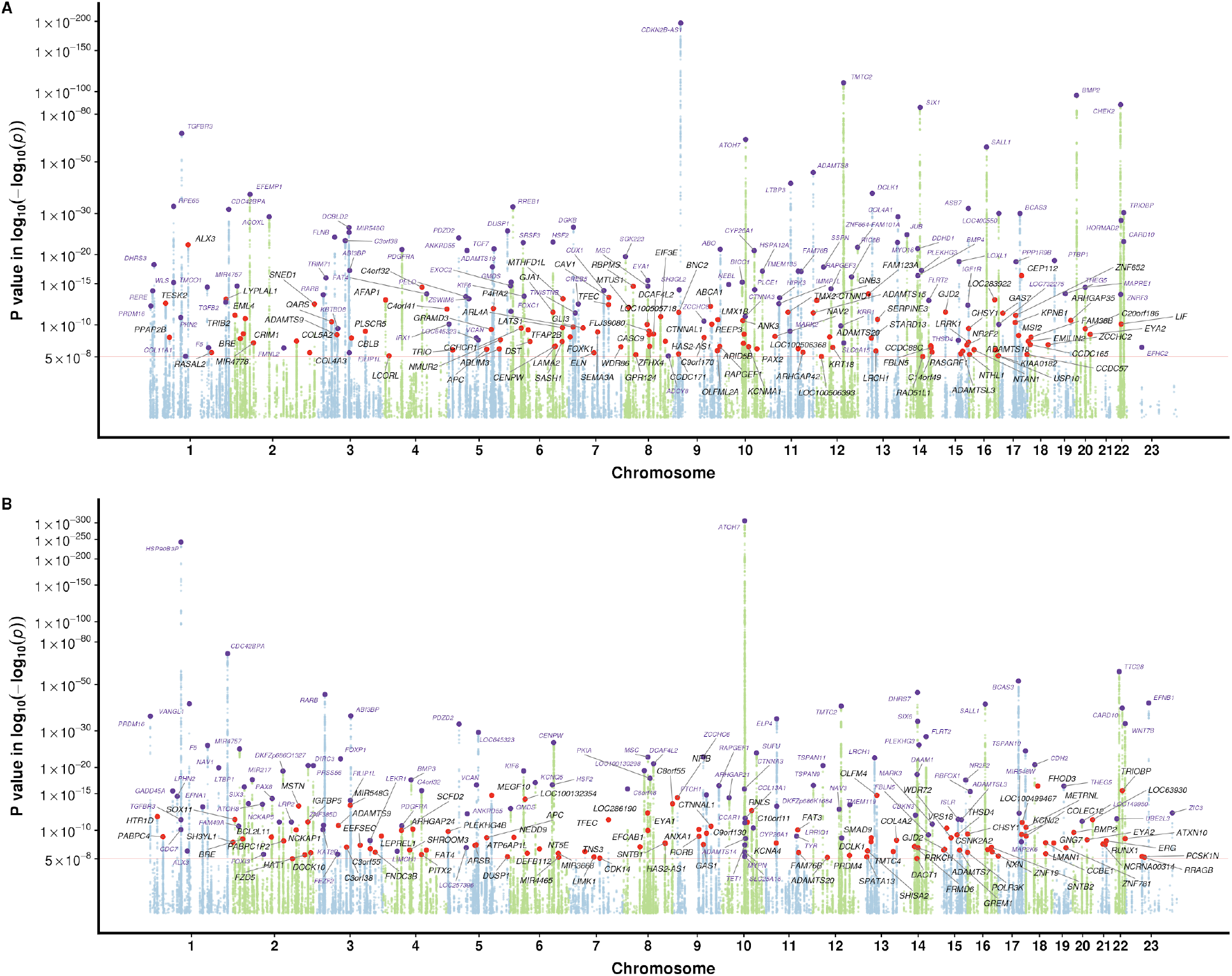
AI enables new genetic discovery for optic nerve head measures. Manhattan plot panel A shows P values for VDD-adjusted VCDR from the meta-analysis of UKB, CLSA, and IGGC (European ancestry). Panel B shows P values for VDD from the meta-analysis of UKB, CLSA, and IGGC (European ancestry). The Y-axis is in log-log scale. The red horizontal line is the genome-wide significance level at P = 5 × 10^−8^. SNPs with P value less than 1 × 10^−4^ are not shown in Manhattan plot. Previously unknown loci are highlighted with red dots, with the nearest gene names in black text. Known SNPs are highlighted with purple dots, with the nearest gene names in purple text.

**Figure 6.**
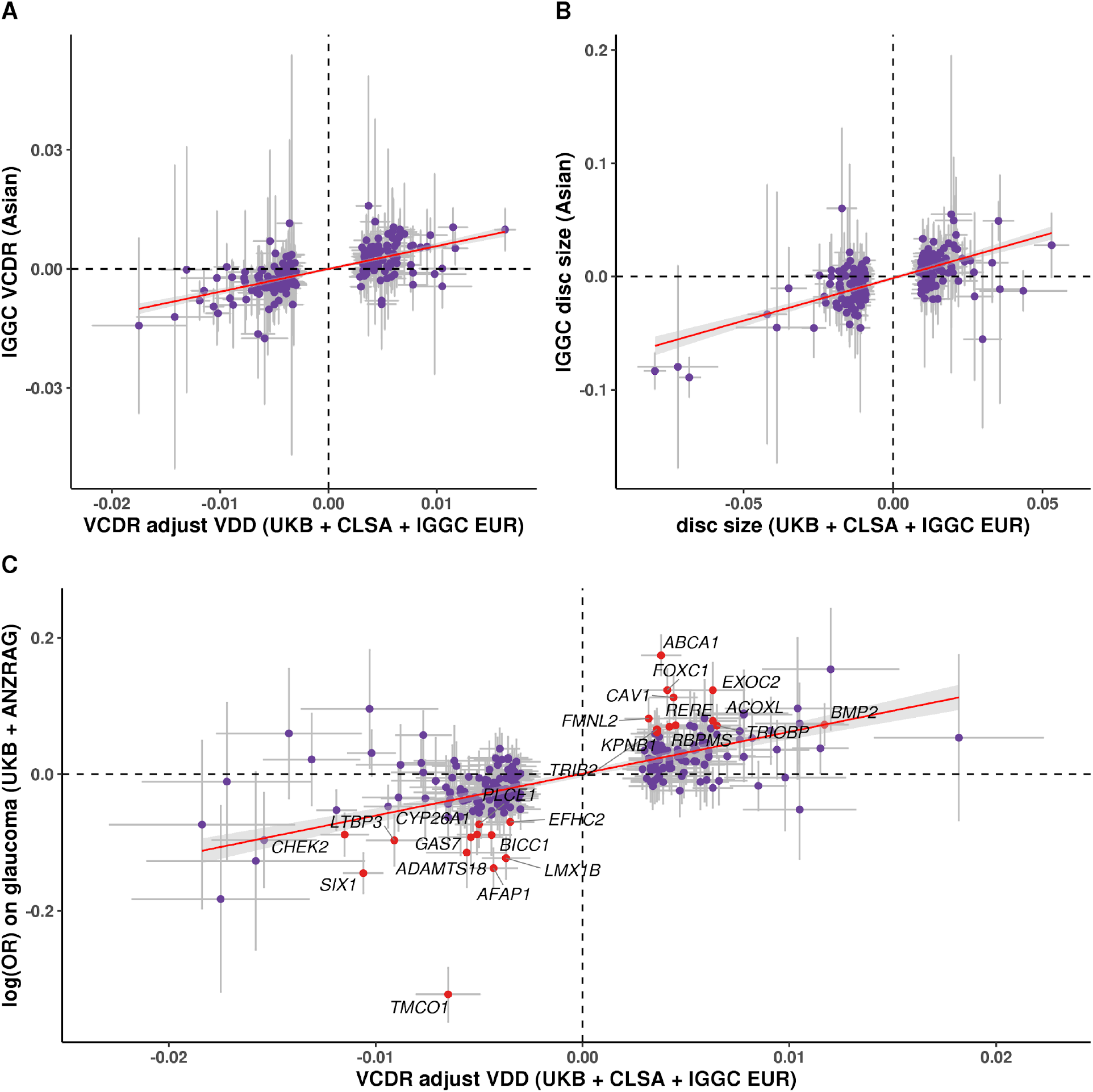
Comparison of the effect sizes for VCDR (adjusted for VDD) and VDD lead SNPs versus those observed in the Asian ancestry group and in independent glaucoma cohorts. Panel A and B show the effect sizes for lead VCDR (adjusted for VDD) and VDD loci (European versus Asian population). Panel C shows the effect sizes for VCDR (adjusted for VDD) lead SNPs versus log odds ratio in meta-analysis of UKB and ANZRAG glaucoma GWAS. The 24 SNPs associated with glaucoma after Bonferroni correction (P<0.05/227 = 2.2 × 10^−4^) are highlighted with red dots, with the nearest gene names in black text.

### Gene prioritization and pathway analysis

We performed TWAS analysis in FUSION based on the VDD-adjusted VCDR and VDD GWAS summary statistics and retinal gene expression data. For VDD-adjusted VCDR we identified 101 genes that were significant after Bonferroni correction for multiple testing, nine of which were not genome-wide significant in the per-SNP analysis (Supplementary Figure 6A and 6B). For VDD we identified 64 genes that were significant after Bonferroni correction for multiple testing, 13 of which were not genome-wide significant in the per-SNP analysis. From SMR analysis, we identified 29 and 24 genes for VDD-adjusted VCDR and VDD, respectively, that were significant after multiple testing. We also compared the genes identified from both FUSION and SMR, 11 and 8 genes overlap from the two methods for VDD-adjusted VCDR and VDD, respectively (Supplementary Figure 6C and 6D). For instance, of the 11 genes that were associated with VDD-adjusted VCDR for the two approaches, 6 genes also passed the HEIDI tests (*P4HTM, SNX32, RASGRF, HAUS4, LRP11, AC012613.2*), suggesting the effects on VCDR may be mediated via these gene expression in retina tissue. The large increase in power resulting from the use of AI grading to improve accuracy and enable substantially larger datasets with multiple images per participant meant we were able to discover many new biological pathways influencing optic nerve head development and aging. Our pathway enrichment analysis uncovered 65 pathways for VCDR and 82 pathways for VDD after Bonferroni correction for multiple testing (Supplementary Table 5 and 6). As well as extracellular matrix pathways uncovered by our previous work, these new pathway analysis uncovered associations with telencephalon (forebrain) regionalization, embryo development, and anatomical tube development. There were several unexpected but statistically robust associations with kidney development (e.g. GO mesonephros development, Praw = 3.45 × 10^−8^, P=0.00053 after correction for multiple comparisons). The genes driving the kidney development pathway enrichment included *BMP2, BMP4, EYA1, FAT4, FOXC1, GLI3, PAX2, RARB, SIX1*, and *SALL1*. Several kidney pathways were also significant in the pathway enrichment analysis applied to our VDD GWAS.

## Discussion

Our results show the promising application of AI algorithms in genetics studies. Large scale biobanks such as UKB and CLSA have accumulated hundreds of thousands of optic nerve images containing important information for glaucomatous optic neuropathy. However, the time-intensive and moderate agreement of manual assessment have impeded the usage of retinal fundus images. We trained a deep learning model using clinically estimated VCDR and VDD, and found the trained model has a high accuracy. The large scale biobank data for both VCDR and IOP allow us to systematically compare the glaucoma risk and optic nerve head parameters across different ancestries. Combining genetic and image data, we doubled the number of loci for both VCDR and VDD, with increased heritability.

The scope of available deep learning models for computer vision tasks is extensive and continuously developing. Various approaches to grade fundus images often utilise intricate data preprocessing methods^23–25^ as well as computationally heavy models and training methods^18,26^. In the instance of statistically powered, large scale population study, fast inference and quick iterations are key, making heavy computational and design costs harder to justify. Here we demonstrate that a relatively lightweight, pretained CNN model is capable of producing highly accurate estimations of VCDR and VDD as evinced by high correlation with clinical grading, improved genetic discovery and further validations in independent samples.

Our AI approach has dramatically accelerated the pace of genetic discoveries. In our previous study, we laboriously manually assessed a subset of UKB images. With the deep learning model trained on clinical measurements, we were able to predict on a new subject within a fraction of a second, making time and effort of image labelling trivial, even when applied to large scale datasets (~1 hour for ~0.3 million images). Sample size is one of the most important limiting factors for genetic discovery. Leveraging the AI-based algorithm and large scale data, we were able to conduct the most powerful GWAS of optic nerve head parameters to date. We doubled the number of genome-wide significant loci for both VCDR and VDD. Interestingly, the estimated SNP-based heritability also increased by ~50% for VCDR and VDD (Supplementary Figure 4); the estimate for VCDR is not substantially lower than the heritability estimate from twin studies (~50%), although given more accurate (AI based) phenotypes, the twin study based heritability estimate may increase. The increased heritability is a result of more accurate measurements, which arises in part due to the higher accuracy of AI-based predictions and in part to the AI approach allowing time-efficient grading of multiple measures per individual.

Many of the newly identified VCDR genes are associated with other eye traits (*e.g*. glaucoma, IOP, exfoliation syndrome, myopia). For some loci associated with IOP, it is likely that they have an effect on VCDR as a secondary effect of the locus first acting on IOP. Loci including genes such as *ABCA1, CAV1, AFAP1* and *LMX1B* were associated with VCDR for the first time; a likely explanation for this association is that the associated variant alters IOP and subsequently VCDR. Over 20 of the VCDR loci are also associated with refractive error, with multiple aspects of eye physiology likely involved (axial length, corneal thickness, retinal ganglion cell function). We also found a significant genome-wide genetic correlation between VCDR (adjusted for VDD) and myopia (rg = 0.3, P = 1×10^−14^), as well as with well studied traits which are associated with myopia such as years of education.^27^

In addition, several of the new VCDR genes provide possible links to retinal ganglion cell biology and they may constitute possible drug repositioning candidates. There are too many to discuss individually but one SNP of interest is rs17855988; this missense variant in the elastin gene (*ELN)* has been associated with facial ageing. Elastin in the sclera is most dense around the optic nerve head^28^ and *ELN* expression has been shown to be high in exfoliation glaucoma lens^29^. A subset of the VDD loci have been found to be associated at genome-wide significance levels in previous glaucoma GWAS. However, in the majority of cases, the association with glaucoma appears to be driven by the lead SNP having a primary effect on VCDR (where the variance explained in VCDR for the peak SNP is larger than that for VDD: *e.g*. SNPs in or near *GMDS, CAV1, MYOF, SIX6, CHEK2, TMTC2)*. Hence, the primary link between the disc parameters and glaucoma is via VCDR rather than via VDD. This is also shown in the lower genetic correlation between glaucoma and VDD (rg = 0.01) compared with glaucoma and VCDR (rg = 0.5).^8,20^ With the high genetic correlation between VCDR and glaucoma, a multitrait analysis has recently shown that including VCDR can improve the power to identify glaucoma genes and to enable the development of polygenic risk score.^20^ Future studies of glaucoma would benefit from incorporating these accurate AI derived VCDR estimates.

Previous studies have looked at the differences between VDD across different ancestries.^30,31^ Our results were consistent with this, with Africans having the largest disc size, followed by those of Asian ancestry. For VCDR, an early study (100 black and 100 white) found that blacks had larger VCDR (mean values: blacks 0.35, white 0.24).^32^ A subsequent larger study (1534 black and 1853 white) reported larger VCDR in blacks (mean values: blacks 0.56, whites 0.49).^33^ A subsequent study in three different Asian ancestries, showed that VCDR values were similar between the studied ancestries (mean VCDR 0.40, 0.42 and 0.40, in Malay, Chinese, Indian, respectively).^34^ It is striking that despite VCDR theoretically being a simple parameter to assess, the mean VCDR varies widely across studies, possibly due to differences in measurement protocol, sex, age and eye disease status. A further study^4^ looked at the 97.5th percentile of VCDR instead of the mean and reported broadly similar values in the Netherlands (0.73), Bangladesh (0.7), Mongolia (0.70), Singapore (0.7), Tanzania (0.7). A major advantage of our study is that we use our AI derived gradings in two population-based cohort studies to systematically assess VCDR differences across ancestries in a consistent study design. By leveraging large sample sizes, we are able to clearly show both Asian and African ancestry individuals have larger VCDR values than Europeans. Our primary results in Figure 2 correct VCDR for VDD, given previous studies showing that correcting for VDD enhances the relevance to glaucoma.^35^

The raised VCDR in Asian and African ancestry individuals living in the UK and Canada is in keeping with elevated glaucoma rates in these ancestries.^36^ When combined with data on IOP, a combination of VCDR and IOP explains the vast majority of the variation between glaucoma rates in Europeans relative to Africans, South Asians and East Asians. Although crucially, our data show (Figure 3) that the relative contributions of VCDR and IOP are clearly different between all 4 major populations groups that we consider. For individuals of European, South Asian or African ancestry, the vast majority of broadly defined glaucoma cases are open angle glaucoma (OAG). In East Asia, angle closure glaucoma (ACG) is common and a limitation of our analysis is that we cannot distinguish between ACG and OAG in all cases - where available we have removed known cases of ACG in the broad glaucoma definition, but some ACG cases will remain.

A strength of our study is that a large number of clinically assessed images were used to train the deep learning model for VCDR and VDD; this allowed us to generate accurate predictions. Our study has shown that the AI-based measurements have a high accuracy. The AI-based optic nerve head assessment has also boosted the available sample size and dramatically expanded gene discovery for these key ocular phenotypes. We show that this deep learning model can also be used to assess future fundus images automatically and rapidly, especially in population-based studies with a large number of images. Moreover, the implementation of transfer-learning techniques shows that AI-aided labelling, with adequate sample size, has a potential to deliver equally successful genetic discoveries in other image based biological phenotypes. Our study has several limitations. Firstly, although our AI approach was able to grade a large proportion of images (particularly in the CLSA study), a small proportion remained ungradable due to poor picture quality. Future studies could explore adversarial architectures to improve clinical ratings of VCDR and VDD. However, a set of high quality truth labels would still be necessary for initial pre-training. Finally, although we were able to use genetic data to clearly identify the major ancestries within UKB and CLSA (European, African, South Asian, East Asian), there remained a group of uncategorized individuals with mixed ancestries that we did not include in our epidemiological or genetic analyses.

To conclude, we showed that AI-based optic nerve head assessment has a high accuracy and this greatly improves our power to discover new genes. These findings provide new insights into the pathogenesis of glaucomatous optic neuropathy. We also use the systematic assessment of VCDR across different ancestries to help explain how the pattern of IOP and VCDR measures underpin observed glaucoma risk; such findings in mixed ancestry groups living in the UK and Canada help explain the differing characteristics of glaucoma across ancestries. For example, relative to Europeans, individuals with East Asian ancestry are more likely to have lower IOP and increased VCDR. Given these East Asians are genetically similar to East Asians in countries such as China and Japan, this provides support for the assertion that normal tension forms of glaucoma predominate in East Asia due to genetic predisposition for high VCDR, despite low IOP.

## Methods

### Study populations

#### UK Biobank

The UK Biobank (UKB) is a population-based cohort study with deep genetic and phenotypic data from ~500,000 participants aged between 40 to 69 years at the time of recruitment (2006-2010), living in the United Kingdom.^37^ Retinal fundus images were available for both left and right eyes from two assessment visits, covering ~85,000 participants (~68,000 participants in the baseline visit and ~19,000 participants in the first repeat assessment visit [2012-2013]). In our previous study, vertical cup-to-disc ratio (VCDR) and vertical disc diameter (VDD) were graded by two clinicians using a custom Java program.^20^ Detailed image processing and quality control methods were described previously.^20^ Briefly, given the time-consuming nature of manual grading, we only graded the left eye images (if the left eye images were ungradable, the right eye images were used instead) and one visit (if the second visit measurements were unavailable, the first visit measurements were used instead) of white British ancestry participants. A total of 67,040 participants with both VCDR and VDD measurements were included in our previous GWAS. In this study, we used a CNN model to grade left and right eye images from two visits for all participants, irrespective of ancestry, with a total of 175,770 images.

In the UKBB, ~488,000 participants were genotyped for 805,426 variants on Axiom arrays (Affymetrix Santa Clara, USA). The genetic data, quality control procedures and imputation methods have been described previously.^37^ Briefly, ~96 million variants were imputed using Haplotype Reference Consortium (HRC) and UK10K haplotype resources^38–40^, and 487,409 individuals passed genotyping quality control. Of them, 438,870 individuals were genetically similar to those of white-British ancestry.^37,41^ For the GWAS in UKB, we retained SNPs with MAF > 0.01 and imputation quality score > 0.8. To verify self-reported diverse ancestry information (data field 21000 in UKB), we used a K-means clustering method based on genetic principal components (PCs). The genetic clusters were compared with self-reported ancestry. Participants within the same self-reported ancestry groups were largely in the same genetic clusters (*e.g*. African [N=9791], South Asian [N=2594], and East Asian [N=9941], detailed in Supplementary Figure 7), and on average ~20% of them have fundus retinal images.

#### The Canadian Longitudinal Study on Aging

The Canadian Longitudinal Study on Aging (CLSA) is a national, longitudinal cohort study of 51,338 participants from 10 Canadian provinces, aged 45 to 85 years at enrollment.^42,43^ Recruitment and baseline data collection were completed in 2015, with participants followed-up every 3 years, and an initial follow-up visit completed in 2018. In this study the nerve head photographs are available for a subset cohort “Comprehensive cohort” of 30,097 participants (for both left and right eyes, and the baseline and first follow-up visit). Retinal fundus imaging was performed using a Topcon (TRC-NW8) non-mydriatic retinal camera, with images saved in jpg format. A random sample of 1000 images was graded by a clinician for both VCDR and VDD using a custom Java program. The latest genome-wide genotype data (August 2019 release) are available for 19,669 participants of the Comprehensive cohort, comprising 794,409 genetic variants genotyped on the Affymetrix Axiom array, and ~40 million genetic variants imputed using the Haplotype Reference Consortium.^39^ Variant- and sample-based quality control procedures were consistent with standards of the UK Biobank^37^ with detailed steps presented in the CLSA support document (available at https://www.clsa-elcv.ca/researchers/data-support-documentation). For the GWAS analysis, we included 18,304 participants of European ancestry based on the K-means cluster method on genetic principal components, and the largest cluster also contains the majority of individuals that self-report European ancestry. SNPs with MAF > 0.01 and imputation quality score > 0.8 were retained in association analysis. From the K-means clustering method, the sample size for African South Asian, and East Asian is 135, 219, and 217, respectively (PC plot was shown in Supplementary Figure 8).

#### The International Glaucoma Genetic Consortium

The International Glaucoma Genetic Consortium (IGGC) is one of the largest international consortia established to identify glaucoma genetic risk variants through large-scale meta-analysis. The phenotype and genotype data of VCDR and optic disc area for IGGC have been previously described elsewhere.^7,44^ It should be noted the optic disc area is not in the same scale as VDD from the AI gradings. When comparing and meta-analyzing the VDD and disc area data, we applied a rank-based inverse normal transformation to AI gradings and rendered them back to disc area scale, as detailed in our previous study.^8^ Publicly available summary statistics were downloaded for individuals of European descent (N_VCDR_= 25,180, N_disc_= 24,509, from the latest HRC imputation), as well as Asian descent (N_VCDR_= 8,373, N_disc_= 7,307).^7,44^

#### Glaucoma GWAS dataset

The glaucoma datasets were described in our previous study^20^, including 7,947 glaucoma cases and 119,318 controls from UK Biobank and 3,071 POAG cases and 6,750 historic controls from the Australian & New Zealand Registry of Advanced Glaucoma (ANZRAG) study.^45,46^ The detailed information of phenotype definition and genetic association analyses were presented in detail previously.^20^ The two datasets were meta-analysed and the GWAS summary statistics were used to look up each of the VCDR loci (adjusted for VDD). We also computed the correlation between the effect size on genome-wide significant VCDR loci and the effect size on glaucoma. The pairwise genetic correlation between VCDR and glaucoma was examined using a genome-wide approach as implemented in LD-Score regression.^47^

#### AI algorithm on retinal images

Three separate CNN models were used to make inferences about image gradability, VCDR, and VDD values of retinal fundus images in UKB. The image gradability (gradable or ungradable) was defined as a binary classification, while the latter two tasks were modelled as regression problems. Images with a higher likelihood of gradability (i.e. designated softmax probability more than 0.5) were assigned as gradable. While a variety of CNN model architectures were tested, the final architecture used for all CNN models was ResNet-34.^48^ Pre-trained weights, initially trained on ImageNet^49^ classification tasks, were utilised for each model as a form of transfer learning. Untrained layers specific to each model were additionally added, forming a custom regression (Relu) and classification (softmax) heads for each respective task. All fundus images were cropped and scaled to a pixel ratio of (1080, 800) before training or validation. We used the highest native resolution for the UKBB training images as we found that using lower resolution negatively impacted inference metrics. The total dataset sizes used for the VCDR, VDD and gradability tasks were 71,950, 50,984, and 75,718, respectively. Each dataset was randomly split into 80% training and 20% validation. The model performance was validated by sample hold out, with final testing performed on images from the CLSA dataset. Model requirements for regression tasks were defined achieving a validation loss equal or lower than human inter-rater loss. The gradebillity task criteria was defined as accuracy above 95%. Both regression tasks utilised mean square error loss function, while the classification model optimised over the binary cross entropy loss function. Training of all models was completed using the FastAI framework^50^, while utilising the in-built data augmentations functionality to improve accuracy and generalisability. The specifics of which augmentations were used can be found in Supplementary Table 7. It should be noted that the regression task for VDD was dependent on image scale, as such, augmentations which introduced scaling were omitted. Training was carried out in two stages: the first involved freezing the pretrained weights and only training the task head; the second, the ‘fine tuning’ stage, all model weights were unfrozen. Each stage was trained with cyclical training rate as described elsewhere^51^, and performed until the validation loss reached a plateau.

#### Optic nerve head parameters, intraocular pressure and glaucoma risk across different ancestries

Previous studies have reported differences in VCDR and VDD values across different ancestry groups.^30,31^ Taking advantage of the diverse ancestries available in UKB and CLSA, we compared our AI derived VCDR and VDD values, as well as intraocular pressure (IOP, corneal-compensated^41^) values across different ancestry groups. We used the K-means clustering method to define ancestry groups based on genetic data (detailed above). Boxplots were used to show the differences of optic nerve head measurements across different ancestry groups (*e.g*. median value, upper and lower quartiles). The mean values of VCDR across different ancestries were estimated after adjusting for age, sex, and VDD. The 97.5th percentile of optic nerve head measurements and its 95% confidence interval (2.5% to 97.5% quantiles) were also calculated based on 1,000 bootstrapped samples, on account of the substantially smaller sample size for individuals of African, East Asian and South Asian ancestry. We then investigated how VCDR and IOP relate to glaucoma risk in different ancestries. The definition of glaucoma cases and controls was detailed in our previous study.^20^ Briefly, in UKB glaucoma cases were ascertained from International Classification of Diseases diagnosis, record-linkage data from general practitioners, and self-reported previous diagnosis. In the CLSA, participants were interviewed in-person with the question “Has a doctor ever told you that you have glaucoma?”. Logistic regression models were used to evaluate the association between genetically-defined ancestry groups and glaucoma risk. In each different model, different covariates were adjusted to evaluate the association of ethnic groups and glaucoma risk. In the base model, only sex and age were adjusted for; the other models also include either IOP, VCDR, or both (IOP & VCDR).

#### Genome-wide association analysis and meta-analysis

For both UKB and CLSA, the VCDR and VDD GWAS association tests were carried out using a linear mixed model (using BOLT-LMM version 2.3)^52^ to account for cryptic relatedness and population stratification, adjusting for sex and age. The first ten principal components were also included in the model to speed up the convergence of computations.^53^ The average values of measurements from left and right eyes and multiple visits (if available) were used, and were first transformed using a rank-based inverse-normal method before association tests.^54^ To account for optic disc size covariation, VCDR grading was adjusted for VDD in GWAS analyses.^8,55^ The VDD-adjusted VCDR and VDD GWAS results from UKB and CLSA were then meta-analysed with those from the IGGC based on the inverse variance-weighted method (METAL software 2011-03-25 release).^56^ We also conducted association tests for VCDR and VDD in African and South Asian populations in UKB. Due to the relatively small size of each of these populations (Supplementary Table 8, less than the recommended sample size of 5000 in BOLT-LMM), PLINK was used instead, after removing related individuals.^57^

SNP-based heritability was calculated by LD score regression (LDSC) from GWAS summary statistics.^47,58^ Bivariate LD score regression was used to estimate the genetic correlation between pairs of traits in European ancestry.^47^ We selected independent SNPs based on the PLINK clumping method with P value < 5×10^−8^, r^2^ < 0.01, and a window of 1Mb from the index variant.^57^ To define novel loci from the AI-based GWAS, we checked previous UKB VCDR and VDD GWAS based on clinician gradings^8,20^, we also looked up the proxy SNPs (r^2^ > 0.8) of top loci and their nearest genes in GWAS Catalog.^59^

#### Cross population genetic effects on optic nerve head parameters

We evaluated the effects of genetics variants on VCDR and VDD cross different populations based on the following methods: 1) we first compared and replicated the AI-based top loci from European ancestry with the GWAS from African and South Asian samples. The effect sizes and standard errors of top loci were shown in a scatter plot for different ancestries; 2) we calculated the trans-ethnic genetic effect correlation for VCDR and VDD using the “Popcorn” package.^60^ Specifically, the GWAS summary statistics for VCDR and VDD from European ancestry were compared with that in Asian and African ancestry.

#### Transcriptome-wide association study and pathway analysis

To prioritize potential causal genes, transcriptome-wide association study analysis (TWAS) was performed in FUSION using GWAS summary statistics and retina gene expression data.^61^ In FUSION, a reference data with both gene expression and genetic variants (SNPs) were used to train predictive models, which were used to impute the expression-trait association directly from large-scale GWAS summary statistics.^61^ The weights of retina gene expression were obtained from 406 individuals from Eye Genotype Expression database (EyeGEx).^61,62^ We also used the EyeGEx to perform a summary data-based Mendelian randomization (SMR) to investigate the association of gene expression levels (exposure) and phenotype (outcome).^63^ The heterogeneity in dependent instruments (HEIDI) tests were used to evaluate the null hypothesis that a single causal variant affecting both gene expression and outcome, and the significance threshold was set at 0.05 (P_HEIDI_ ≥ 0.05 not reject the null hypothesis).^63^ Pathway analysis were conducted in MAGMA as implemented in FUMA (version 1.3.6).^64,65^ All other analyses were performed with R software.^66^

## Data availability

UK Biobank data are available through the UK Biobank Access Management System https://www.ukbiobank.ac.uk/. We will return the derived data fields following the UK biobank policy and in due course they will be available through the UK Biobank Access Management System.

Data are available from the Canadian Longitudinal Study on Aging (www.clsa-elcv.ca) for researchers who meet the criteria for access to de-identified CLSA data.

## Acknowledgements

This work was conducted using the UK Biobank Resource (application number 25331) and publicly available data from the International Glaucoma Genetics Consortium. The UK Biobank was established by the Wellcome Trust medical charity, Medical Research Council (UK), Department of Health (UK), Scottish Government, and Northwest Regional Development Agency. It also had funding from the Welsh Assembly Government, British Heart Foundation, and Diabetes UK. The eye and vision dataset has been developed with additional funding from The NIHR Biomedical Research Centre at Moorfields Eye Hospital and the UCL Institute of Ophthalmology, Fight for Sight charity (UK), Moorfields Eye Charity (UK), The Macula Society (UK), The International Glaucoma Association (UK) and Alcon Research Institute (USA). This work was also supported by grants from the National Health and Medical Research Council (NHMRC) of Australia (#1107098; 1116360, 1116495, 1023911), the Ophthalmic Research Institute of Australia, the BrightFocus Foundation, UK and Eire Glaucoma Society and Charitable Funds from Royal Liverpool University Hospital. SM, JEC, and AWH are supported by NHMRC Fellowships. XH is supported by the University of Queensland Research Training Scholarship and QIMR Berghofer PhD Top Up Scholarship. We thank Scott Wood, John Pearson and Scott Gordon from QIMR Berghofer for support. The NEIGHBORHOOD consortium is supported by NIH grants P30 EY014104, R01 EY015473 and R01 EY022305. AI engineering was performed at and funded by Max Kelsen.

This research was made possible using the data/biospecimens collected by the Canadian Longitudinal Study on Aging (CLSA). Funding for the Canadian Longitudinal Study on Aging (CLSA) is provided by the Government of Canada through the Canadian Institutes of Health Research (CIHR) under grant reference: LSA 94473 and the Canada Foundation for Innovation. This research has been conducted using the CLSA dataset [Baseline Comprehensive Dataset version 4.0, Follow-up 1 Comprehensive Dataset version 1.0], under Application Number 190225. The CLSA is led by Drs. Parminder Raina, Christina Wolfson and Susan Kirkland.

## Disclaimer

The opinions expressed in this manuscript are the author’s own and do not reflect the views of the Canadian Longitudinal Study on Aging.

## Author contributions

S.M., M.T., X.H. and K.S. designed the research. S.M., M.T., A.W.H., J.E.C. and P.G. obtained the funding. X.H., K.S., M.T. and S.M. executed the research and analysed the data. X.H., K.S., M.T. and S.M. wrote the first draft of the manuscript. A.Q., H.N.M., C.B., M.T., O.S., P.G., J.E.C. and A.W.H. interpreted the results. All authors contributed to manuscript revision and approved the submitted version.

