## Supplementary information for "Automated AI labelling of optic nerve head enables new insights into cross-ancestry glaucoma risk and genetic discovery in over 280,000 images from the UK Biobank and Canadian Longitudinal Study on Aging"

**Affiliations:**

300 Herston Road, Brisbane, Queensland 4006, Australia.

### Table of contents

**Supplementary Figures:** 3

Figure S1. Image gradable possibility in the UK Biobank and the Canadian Longitudinal Study on Aging (CLSA). 3

Figure S2. Correlation between AI and clinician gradings for vertical cup-disc-ratio and vertical disc diameter measurements in the UK Biobank and the Canadian Longitudinal Study on Aging (CLSA). 4

Figure S3. Manhattan plot displaying AI-based GWAS for optic nerve head measures in the UK Biobank and Canadian Longitudinal Study on Aging participants. 6

Figure S4. AI-based gradings increase SNP-based heritability for optic nerve head parameters. 7

Figure S5. Comparison of the effect sizes for VCDR (adjusted for VDD) and VDD lead SNPs versus that in African and South Asian ancestry groups. 8

Figure S6. Transcriptome-wide association study analysis (TWAS) for VCDR (adjusted for VDD) and VDD. 9

Figure S7. K-means clustering of the first 20 genetic principal components in the UK Biobank. 10

Figure S8. K-means clustering of genetic principal components in the Canadian Longitudinal Study on Aging (CLSA). 12

**Supplementary Tables:** 13

Table S1. GWAS of AI-based optic nerve head measurements in the UK Biobank and Canadian Longitudinal Study on Aging participants - linkage disequilibrium score regression. 13

Table S2. 230 lead genome-wide significant independent SNPs for VCDR (adjusted for disc diameter) from the meta-analysis of UKB, CLSA, and IGGC (European ancestry). 14

Table S3. 231 lead genome-wide significant independent SNPs for vertical disc diameter from the meta-analysis of UKB, CLSA, and IGGC (European ancestry). 27

Table S4. Cross population genetic effect correlation for optic nerve head parameters. 39

Table S5. Pathway analysis of VDD-adjusted VCDR (list of 65 significant pathways after Bonferroni correction) 40

Table S6. Pathway analysis of VDD (list of 82 significant pathways after Bonferroni correction). 42

Table S7. Training specifics of Convolutional Neural Network algorithms. 46

Table S8. Summary of study datasets. 47

**Supplementary References** 48

#### Supplementary Figures:

##### Figure S1. Image gradable possibility in the UK Biobank and the Canadian Longitudinal Study on Aging (CLSA).

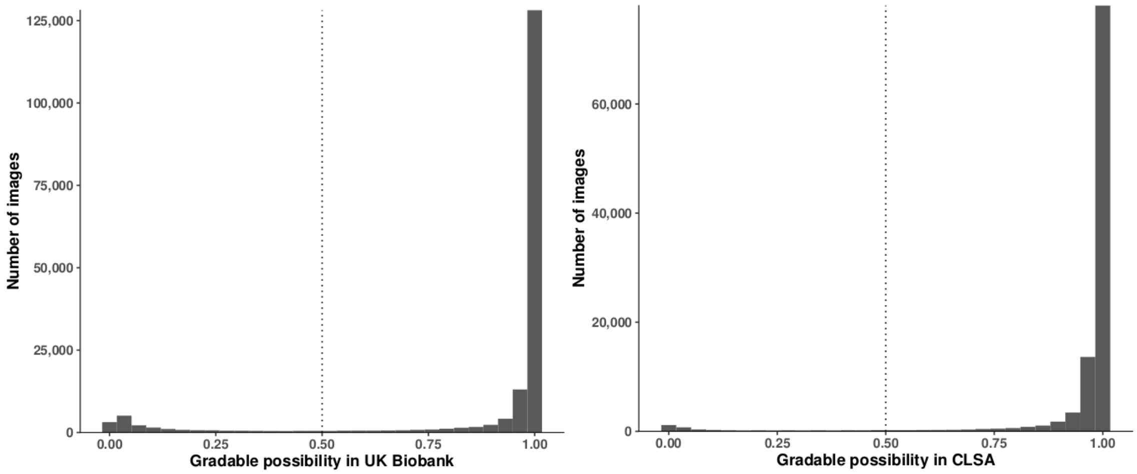


Histogram of image gradable possibility in the UK Biobank (UKB) and the Canadian Longitudinal Study on Aging (CLSA) data sets, where the x-axis represents the grable possibility, and y-axis represents the number of images, and the vertical dashed lines are the possibility threshold of 0.5 to label image gradable. In the UKB, there are 175,770 images in total covering 85,736 participants, and 81,492 (95%) individuals have at least one gradable fundus image. In the CLSA cohort, 29,635 participants have 106,330 images in total (left and right eyes and two assessment visits, **×4**), and 29,341 (99%) individuals have at least one gradable fundus image.

**Figure S2. Correlation between AI and clinician gradings for vertical cup-disc-ratio and vertical disc diameter measurements in the UK Biobank and the Canadian Longitudinal Study on Aging (CLSA).**

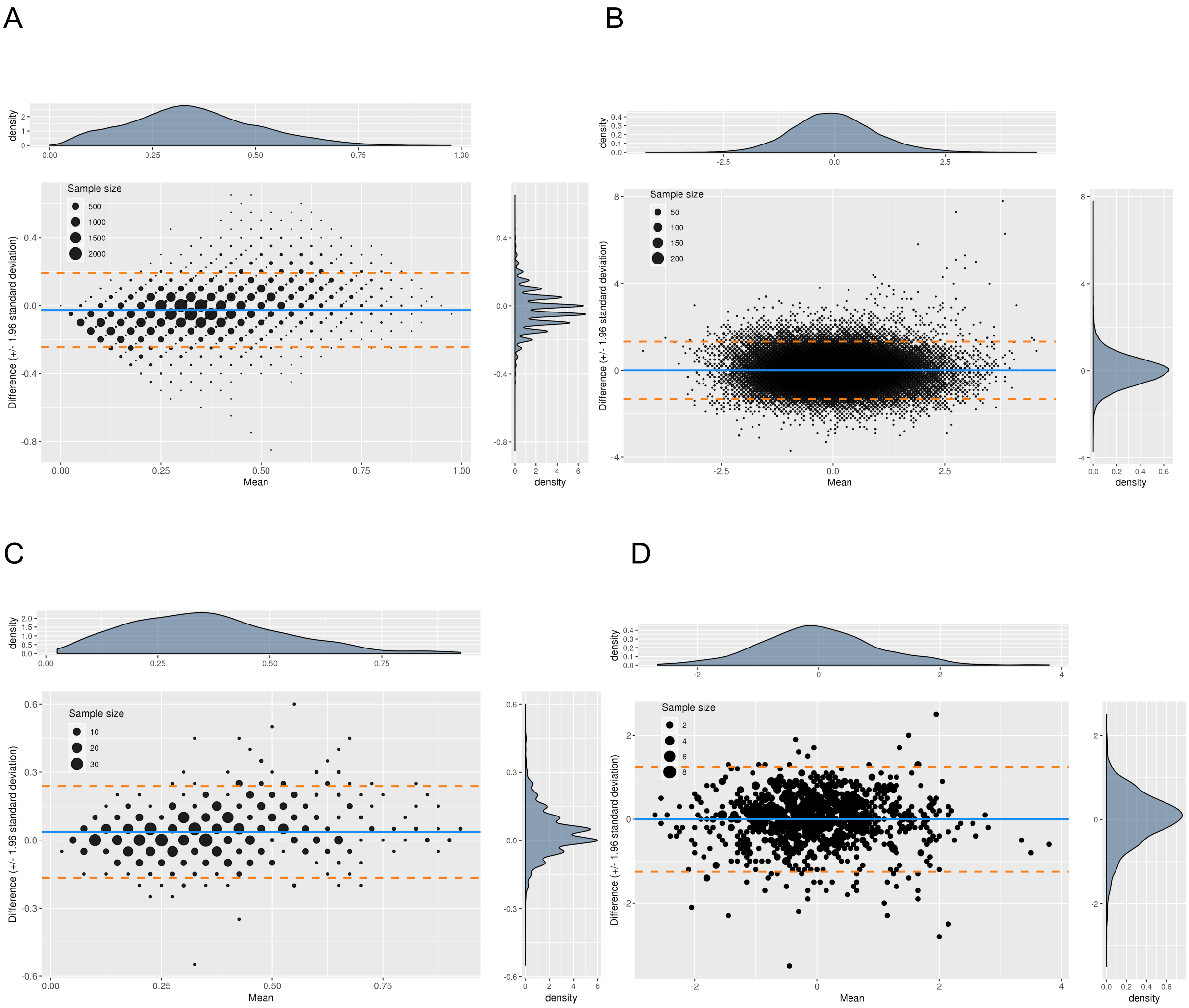


Bland-Altman plots for vertical cup-disc-ratio (VCDR) and vertical disc diameter (VDD. Panel A and B show VCDR and VDD in the UK Biobank, respectively. Panel C and D show VCDR and VDD in the CLSA, respectively. The x-axis represents the mean value of two measurements (AI and clinician gradings), the y-axis represents the difference between two measurements, the blue line is the mean value of difference, and the dashed orange lines are the 95% limits of agreement (95% confidence interval for the mean value of difference). The black dots are scaled by the number of samples. The right and top sub-panels are the density plots for the difference of the measurements and the mean value of measurements, respectively.

In the UKB (panel A and B), ~57,000 AI-based and clinician gradings were used to make Bland-Altman plots for VCDR and VDD. The Pearson's correlation coefficient of the VCDR and VDD measurements was 0.81 (95% confidence interval [CI]: 0.80-0.81) and 0.77 (95% CI: 0.77-0.78), respectively.

In the CLSA (panel C and D), one thousand images were randomly selected for testing, and the Pearson's correlation coefficient of the VCDR and VDD measurements was 0.84 (95% CI: 0.82-0.86) and 0.80 (95% CI: 0.77-0.82), respectively.

The AI-based VCDR and VDD measurements exhibited a higher correlation with clinician gradings than that between two clinicians, for instance, our previous studies showed the Pearson’s correlation coefficient of VCDR and VDD measurements between two clinicians was 0.75 (95% CI:0.72-0.77) and 0.64 (95% CI: 0.61–0.67), respectively.

**Figure S3. Manhattan plot displaying AI-based GWAS for optic nerve head measures in the UK Biobank and Canadian Longitudinal Study on Aging participants.**

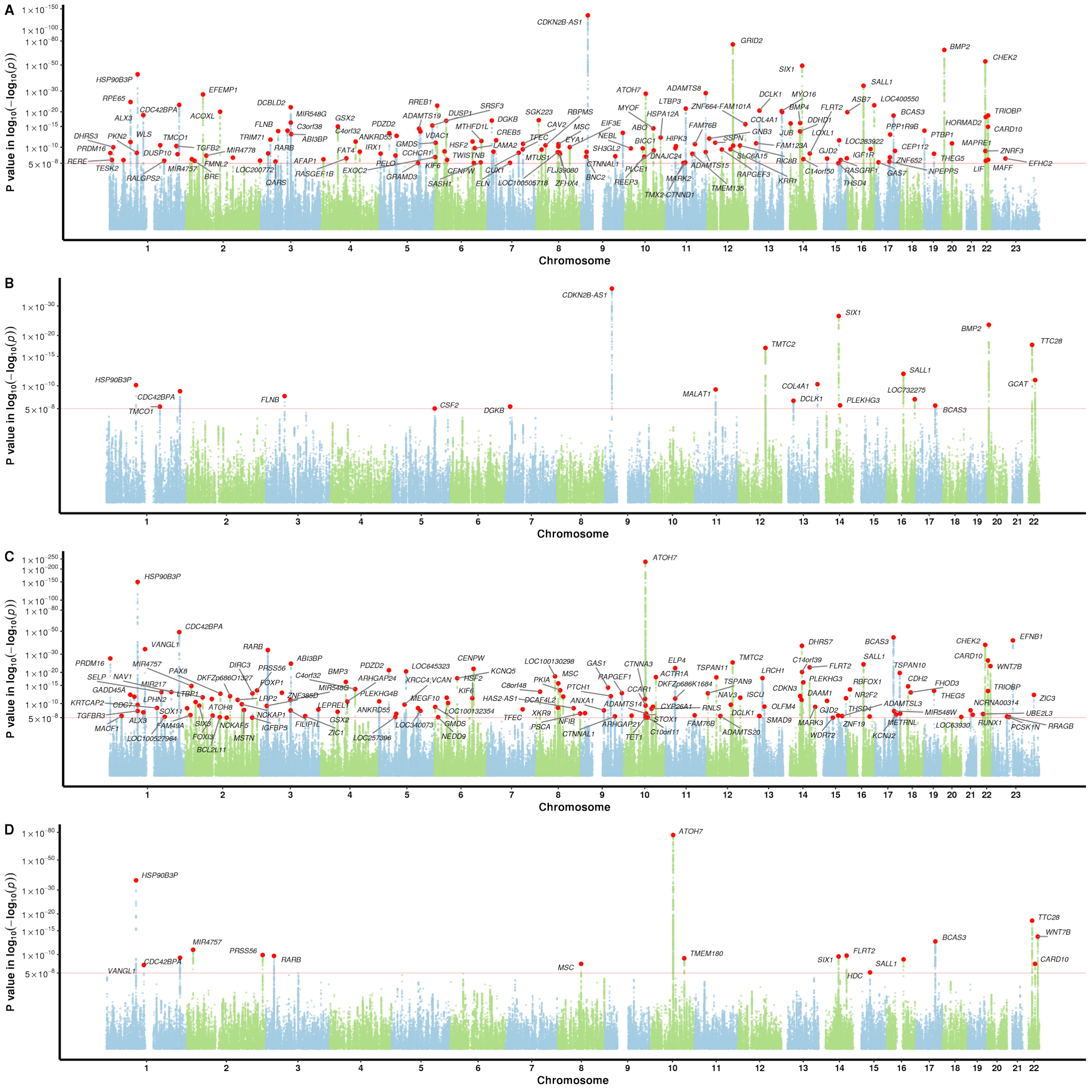


Panel A and B show P values for VCDR (adjusted for VDD) in the UKB and CLSA cohort, respectively. Panel C and D show P values for VDD in the UKB and CLSA cohort, respectively. The red line is the genome-wide significance level at $5 \times 10^{-8}$. SNPs with P values less than 0.01 are not shown in the Manhattan plot. SNPs are highlighted with red dots, with the nearest gene names in black text.

**Figure S4. AI-based gradings increase SNP-based heritability for optic nerve head parameters.**

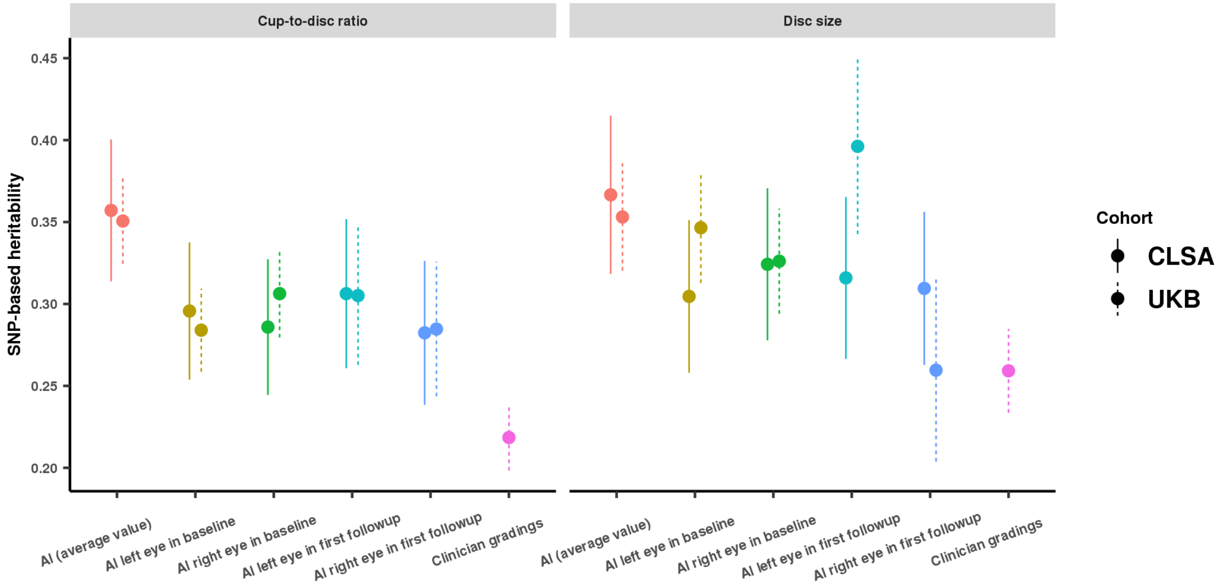


The figure shows SNP-based heritability for VCDR (panel A) and disc size (panel B) from AI-based gradings (single measures in left or right eyes, baseline or first follow-up visit, and average value of multiple measures) and clinician gradings. The X-axis is different types of gradings (with different colors), the Y-axis is SNP-based heritability (point estimation with one standard error from LD score regression approach). The estimations from UKB and CLSA are shown in different linetypes (UKB in solid lines, and CLSA in dotted lines). These results indicate the higher accuracy of AI-based single measure per individual contributes to the increase of heritability estimation, and averaging of multiple measures per individual can further increase the heritability. For instance, the SNP-based heritability for VCDR was 0.22 from clinician gradings (only single measure), whereas the heritability increased to 0.35 from AI-based GWAS (average of multiple measures). The SNP-based heritability from a single AI measure (left or right eyes in the baseline or first follow-up visit) was ~0.3, which is roughly in the middle of clinician gradings and AI-based multiple measures. The sample sizes for single measure in UKB follow-up visits and CLSA are relatively small (~ 15,000), therefore their SNP-based heritabilities have a larger standard error.

**Figure S5. Comparison of the effect sizes for VCDR (adjusted for VDD) and VDD lead SNPs versus that in African and South Asian ancestry groups.**

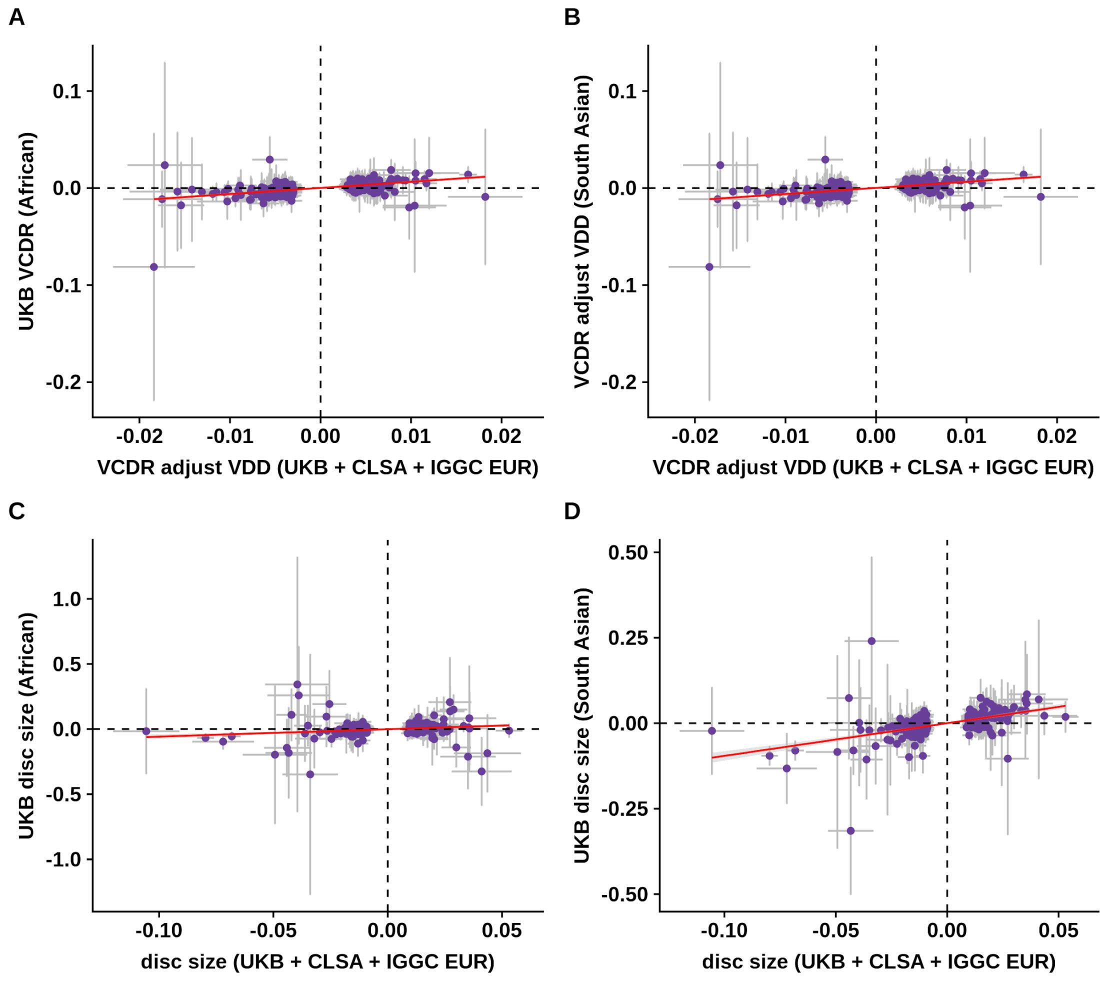


This figure shows the effect sizes in European versus that in African and South Asian populations for lead VCDR (adjusted for VDD) and VDD loci. The vertical and horizontal error bars are the 95% confidence interval for SNP effect sizes. Panel A shows a scatter plot of effect sizes for VDD-adjusted VCDR in European and African population (Pearson's correlation coefficient is 0.26, P = 8.7 × $10^{-5}$). Panel B shows the effect sizes for VDD-adjusted VCDR in European and South Asian population (Pearson's correlation coefficient is 0.43, P = 1.1 × $10^{-11}$). Panel C shows no obvious correlation of effect sizes for disc size in European and African populations (Pearson's correlation coefficient is 0.10, P = 0.14). Panel D shows the effect sizes for disc size in European and South Asian population (Pearson's correlation coefficient is 0.50, P = 9.1 × $10^{-16}$). The GWAS sample size in African and South Asian ancestries is very small (N = 2,245, and 21,00 respectively), therefore, the vertical error bars are very large.

**Figure S6. Transcriptome-wide association study analysis (TWAS) for VCDR (adjusted for VDD) and VDD.**

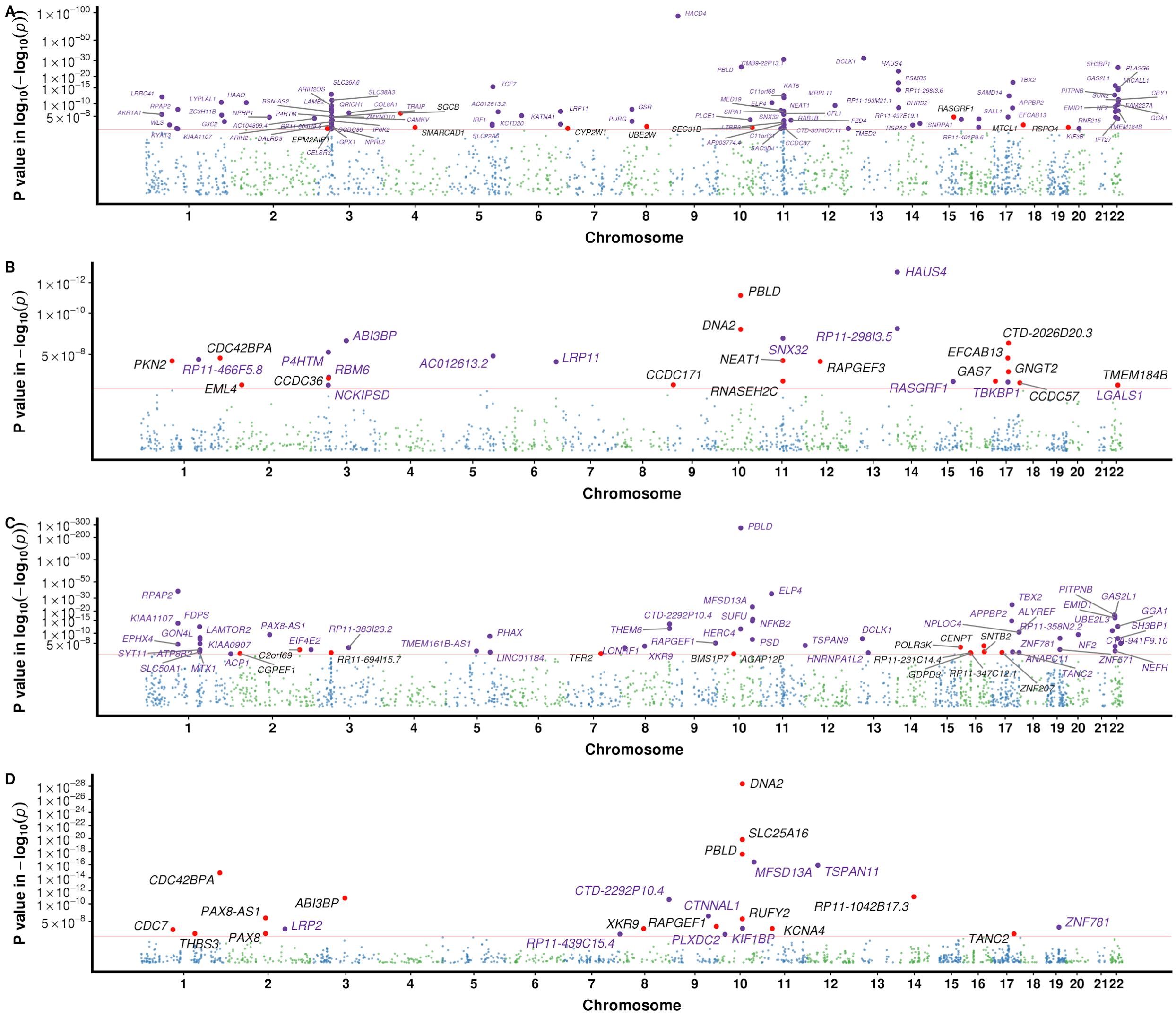


Transcriptome-wide association study analysis from FUSION and SMR: panel A and panel B shows FUSION and SMR P values, respectively, for VDD-adjusted VCDR from the meta-analysis of UKB, CLSA, and IGGC (European ancestry). Panel C and panel D shows FUSION and SMR P values, respectively, for VDD from the meta-analysis of UKB, CLSA, and IGGC (European ancestry). The Y-axis is P value in log-log or log scale. The red line is the P value at Bonferroni correction of the number of genes tested. Genes with P value less than 0.1 were not shown. In FUSION results (panel A and C), genes that are significant after Bonferroni correction for multiple testing are highlighted with purple dots, with the nearest gene names in purple text, of which not genome-wide significant in the per-SNP analysis (best GWAS SNP P value < 5× $10^{-8}$ ) are highlighted with red dots, with the nearest gene names in black text. In SMR results (panel B and D), genes that are significant after Bonferroni correction for multiple testing are highlighted with red dots, with the nearest gene names in black text, of which not pass the HEIDI tests ($P_{HEIDI}$ < 0.05) are highlighted with purple dots, with the nearest gene names in purple text.

**Figure S7. K-means clustering of the first 20 genetic principal components in the UK Biobank.**

**UK Biobank K−means clustering of 20 PCs**

| K−means clusters | African | Any other Asian | Any other Black | Any other white | Asian (British) | Bangladeshi | Black (British) | British | Caribbean | Chinese | Indian | Irish | Other/Unknown | Pakistani |
|---|---|---|---|---|---|---|---|---|---|---|---|---|---|---|
| 1 | 2967 | 1 | 82 | 0 | 0 | 0 | 22 | 2 | 3842 | 0 | 0 | 0 | 977 | 0 |
| 2 | 0 | 1 | 0 | 480 | 0 | 0 | 0 | 27126 | 0 | 1 | 1 | 2137 | 235 | 0 |
| 3 | 0 | 0 | 0 | 290 | 0 | 0 | 0 | 32816 | 0 | 0 | 1 | 441 | 168 | 0 |
| 4 | 0 | 0 | 0 | 89 | 0 | 0 | 0 | 8847 | 0 | 0 | 0 | 23 | 49 | 0 |
| 5 | 1 | 234 | 0 | 1476 | 4 | 0 | 0 | 2005 | 0 | 0 | 16 | 10 | 1394 | 1 |
| 6 | 0 | 0 | 0 | 436 | 0 | 0 | 0 | 47349 | 0 | 0 | 0 | 102 | 161 | 0 |
| 7 | 0 | 0 | 0 | 525 | 0 | 0 | 0 | 59033 | 0 | 0 | 2 | 74 | 310 | 0 |
| 8 | 233 | 6 | 30 | 12 | 0 | 0 | 2 | 24 | 421 | 0 | 1 | 0 | 1351 | 3 |
| 9 | 0 | 0 | 0 | 866 | 0 | 0 | 1 | 60072 | 0 | 0 | 0 | 40 | 254 | 0 |
| 10 | 2 | 1107 | 5 | 13 | 34 | 220 | 0 | 85 | 35 | 1 | 5611 | 3 | 1275 | 1729 |
| 11 | 0 | 366 | 0 | 1 | 2 | 1 | 0 | 5 | 0 | 1483 | 1 | 0 | 745 | 0 |
| 12 | 0 | 1 | 0 | 3318 | 0 | 0 | 0 | 51423 | 0 | 0 | 2 | 46 | 381 | 0 |
| 13 | 0 | 18 | 0 | 150 | 1 | 0 | 0 | 130 | 1 | 17 | 1 | 2 | 739 | 2 |
| 14 | 1 | 0 | 0 | 36 | 0 | 0 | 0 | 24258 | 0 | 0 | 0 | 24 | 82 | 0 |
| 15 | 0 | 0 | 0 | 133 | 0 | 0 | 0 | 12907 | 0 | 0 | 0 | 30 | 74 | 0 |
| 16 | 0 | 0 | 0 | 113 | 0 | 0 | 0 | 10360 | 0 | 0 | 0 | 9138 | 227 | 0 |
| 17 | 0 | 1 | 1 | 722 | 0 | 0 | 0 | 46856 | 0 | 0 | 3 | 553 | 261 | 0 |
| 18 | 0 | 11 | 0 | 3517 | 1 | 0 | 0 | 5249 | 0 | 0 | 21 | 49 | 1072 | 12 |
| 19 | 0 | 0 | 0 | 247 | 0 | 0 | 1 | 39278 | 0 | 1 | 0 | 25 | 136 | 0 |
| 20 | 0 | 1 | 0 | 3336 | 0 | 0 | 0 | 2476 | 0 | 0 | 0 | 21 | 172 | 0 |

Self−reported ethnic background

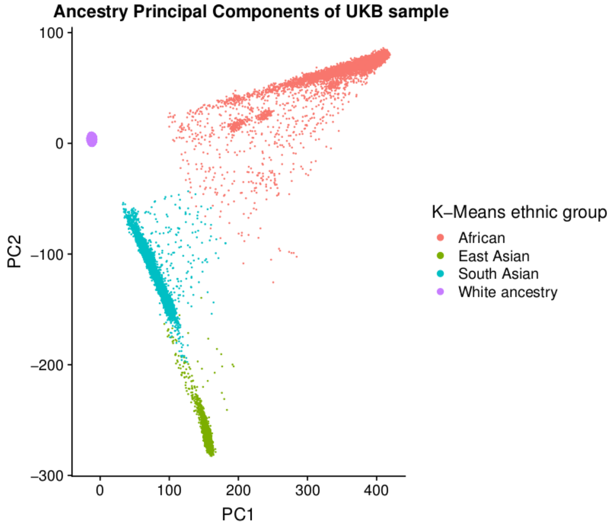


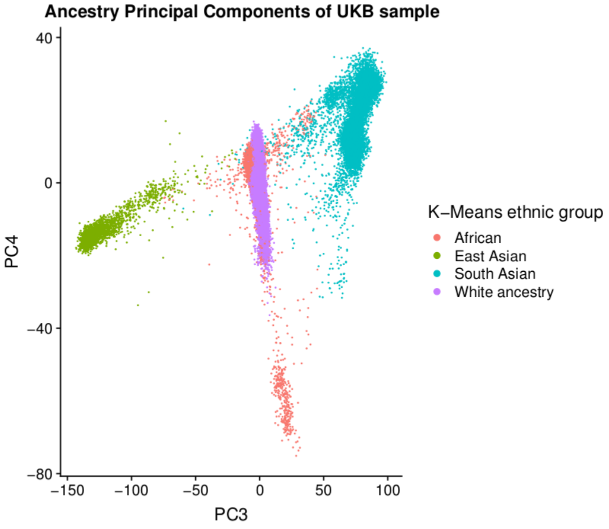


The upper panel shows the self-reported ethnic background (data field 21000 in UKB) versus K-means clustering on the first 20 principal components (PCs). Self-reported African are mainly in cluster 1 and 8, which also include samples reported as “Caribbean”, “Any other Black background”, and “Other/Unknown”. Self-reported Chinese (East Asian) are mainly in cluster 11, which also includes samples reported as “Any other Asian background”, and “Other/Unknown”. Self-reported Indian (South Asian) are mainly in cluster 10, which also includes samples reported as “Pakistani”, “Bangladeshi”, “Any other Asian background”, and “Other/Unknown”. The lower panels show the four PCs (PC1-4) for different ancestry backgrounds. Participants within the same self-reported ancestry groups were largely in the same genetic clusters, and the sample size is  in line with a recent study (*e.g.* African (N=9791), South Asian (N=2594), and East Asian (N=9941))[1].

**Figure S8. K-means clustering of genetic principal components in the Canadian Longitudinal Study on Aging (CLSA).**

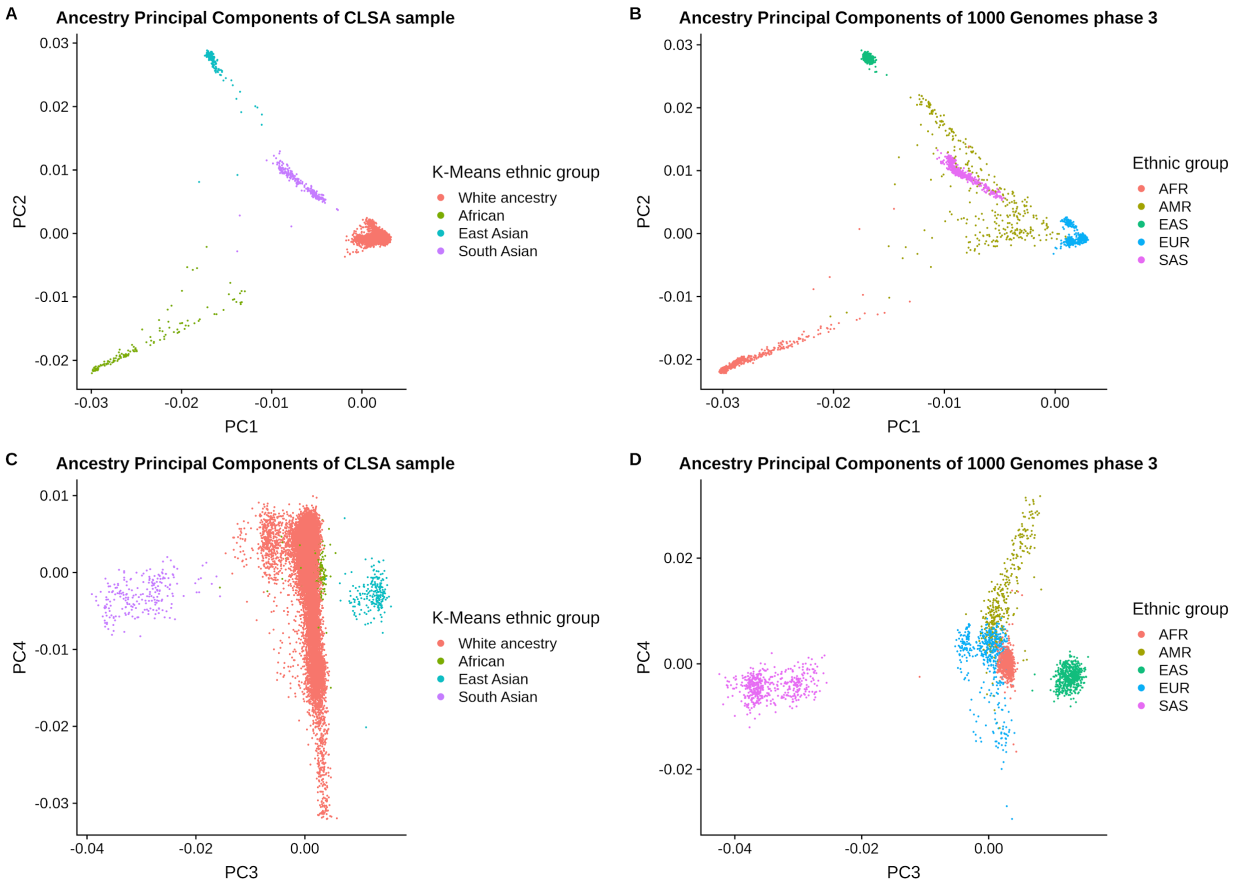


This figure shows the ethnic groups based on K-means clustering in the CLSA sample. The CLSA samples are merged with 1000 genomes phase 3 data (N = 2504) to calculate PCs. Panel A shows the PC1 vs PC2 in the CLSA sample, and panel B shows that in 1000 genomes data. Panel C shows the PC3 vs PC4 in the CLSA sample, and panel D shows that in 1000 genomes data.

#### Supplementary Tables:

##### Table S1. GWAS of AI-based optic nerve head measurements in the UK Biobank and Canadian Longitudinal Study on Aging participants - linkage disequilibrium score regression.

| | UKB AI | CLSA AI | UKB clinician |
|---|---|---|---|
| VCDR (adjusted for VDD) | 145 loci<br>LDSC $h^2$= 0.3506 (0.0261)<br>Intercept: 1.0588 (0.0114)<br>Ratio: 0.1096 (0.0214) | 19 loci<br>LDSC $h^2$= 0.3571 (0.0433)<br>Intercept: 1.0155 (0.0076)<br>Ratio: 0.1057 (0.052) | 76 loci<br>LDSC $h^2$=0.2185 (0.0202)<br>Intercept: 1.041 (0.0095)<br>Ratio: 0.1223 (0.0282) |
| VDD | 142 loci<br>LDSC $h^2$= 0.3531 (0.0328)<br>Intercept: 1.053 (0.01)<br>Ratio: 0.0973 (0.0184) | 17 loci<br>LDSC $h^2$= 0.3666 (0.0483)<br>Intercept: 1.0172 (0.0075)<br>Ratio: 0.1126 (0.0487) | 91 loci<br>LDSC $h^2$ = 0.2592 (0.0256)<br>Intercept: 1.0491 (0.0095)<br>Ratio: 0.1225 (0.0236) |

$h^2$, SNP-based heritability; LDSC, linkage disequilibrium score regression; the attenuation ratio was calculated as (Intercept-1) / ($\underline{\chi 2}$-1). The intercept from LDSC has been used to evaluate population stratification, but it can be larger than 1 with increased sample size. The attenuation ratio can calibrate the intercept against increased chi-squared statistics (i.e. polygenicity). A previous study in over 20 traits showed on average the attenuation ratio is 8.2%, and few traits > 15%.[2]

#### Table S2. 230 lead genome-wide significant independent SNPs for VCDR (adjusted for disc diameter) from the meta-analysis of UKB, CLSA, and IGGC (European ancestry).

Abbreviations: SNP, single nucleotide polymorphism. CHR, Chromosome; BP, position, A1, effect allele; A2, non-effect allele; SE, Standard error; P, p value; novel, “1” novel SNPs for VCDR.

| SNP | CHR | BP | A1 | A2 | BETA | SE | P | Nearest gene name | Genes within 200kb | novel |
|---|---|---|---|---|---|---|---|---|---|---|
| rs12725529 | 1 | 3051749 | T | C | -0.0076 | 0.0011 | 8.7E-13 | PRDM16 | ACTRT2(+112.3kb)\|LINC00982(+67.46kb)\|MIR4251(+7.15kb)\|PRDM16(0) | 0 |
| rs159961 | 1 | 8484228 | T | C | 0.0045 | 0.0006 | 1E-14 | RERE | RERE(0)\|SLC45A1(+80kb) | 0 |
| rs6690264 | 1 | 12613422 | A | G | -0.0047 | 0.0005 | 7.6E-19 | DHRS3 | AADACL3(-162.7kb)\|AADACL4(-91.14kb)\|C1orf158(-192.7kb)\|DHRS3(-14.52kb)\|MIR6730(-25.56kb)\|SNORA59A(+45.97kb)\|SNORA59B(+45.97kb)\|VPS13D(+41.32kb) | 0 |
| rs7536135 | 1 | 45859332 | A | G | -0.0044 | 0.0006 | 4.2E-13 | TESK2 | AKR1A1(-157.1kb)\|CCDC163P(-100.3kb)\|HPDL(+64.99kb)\|LINC01144(+88.04kb)\|MMACHC(-106.5kb)\|MUTYH(+53.19kb)\|NASP(-190.3kb)\|PRDX1(-117.4kb)\|TESK2(0)\|TOE1(+49.68kb)\|ZSWIM5(+187.1kb) | 1 |
| rs4638151 | 1 | 56955386 | T | C | 0.0034 | 0.0006 | 1.4E-09 | PPAP2B | PPAP2B(-5.032kb)\|PRKAA2(-155.6kb) | 1 |
| rs34151819 | 1 | 68773910 | T | C | -0.0172 | 0.0021 | 6.7E-16 | WLS | DEPDC1(-165.9kb)\|GNG12-AS1(+105.2kb)\|LOC101927220(-188.4kb)\|MIR1262(+124.6kb)\|RPE65(-120.6kb)\|WLS(+75.63kb) | 0 |
| rs3125918 | 1 | 68846246 | A | G | -0.0064 | 0.0005 | 6.1E-33 | RPE65 | DEPDC1(-93.59kb)\|GNG12-AS1(+177.6kb)\|LOC101927220(-116.1kb)\|MIR1262(+197kb)\|RPE65(-48.26kb)\|WLS(+148kb) | 0 |
| rs786913 | 1 | 89295330 | A | G | -0.0036 | 0.0005 | 1.7E-11 | PKN2 | CCBL2(-106.1kb)\|GBP3(-177kb)\|GTF2B(-22.99kb)\|LOC101927891(+144.4kb)\|PKN2(0)\|RBMXL1(-149.8kb) | 0 |
| rs4658101 | 1 | 92077409 | A | G | 0.0115 | 0.0007 | 6.5E-67 | TGFBR3 | CDC7(+86.09kb)\|TGFBR3(-68.49kb) | 0 |
| rs2376262 | 1 | 103404219 | T | C | -0.003 | 0.0005 | 4.6E-08 | COL11A1 | COL11A1(0) | 0 |
| rs10857812 | 1 | 110627923 | A | T | -0.0057 | 0.0006 | 9.1E-23 | ALX3 | AHCYL1(+61.56kb)\|ALX3(+14.6kb)\|CSF1(+154.3kb)\|KCNC4(-125.4kb)\|KCNC4-AS1(-123.1kb)\|SLC6A17(-65.21kb)\|STRIP1(+30.66kb)\|UBL4B(-27.14kb) | 1 |
| rs2814471 | 1 | 165739598 | T | C | -0.0065 | 0.0008 | 3.1E-15 | TMCO1 | ALDH9A1(+71.7kb)\|LOC400794(+188.3kb)\|LOC440700(+60.4kb)\|MGST3(+114.2kb)\|MIR3658(-137.6kb)\|TMCO1(+1.439kb)\|UCK2(-57.13kb) | 0 |
| rs6666046 | 1 | 169545422 | A | T | 0.0033 | 0.0006 | 1E-08 | F5 | BLZF1(+179.6kb)\|CCDC181(+148.8kb)\|F5(0)\|SELE(-146.4kb)\|SELL(-114.4kb)\|SELP(-12.66kb)\|SLC19A2(+90.21kb) | 0 |
| rs375818773 | 1 | 178065592 | A | G | 0.0071 | 0.0013 | 2.5E-08 | RASAL2 | LOC730102(+58.45kb)\|RASAL2(0)\|RASAL2-AS1(+2.464kb)\|SEC16B(+126.5kb) | 1 |
| rs6658835 | 1 | 218520995 | A | G | -0.0044 | 0.0006 | 3.1E-13 | TGFB2 | LOC728463(+1.975kb)\|RRP15(+9.67kb)\|TGFB2(0) | 0 |

| | | | | | | | | | | |
|---|---|---|---|---|---|---|---|---|---|---|
| rs13376300 | 1 | 219522662 | A | C | 0.005 | 0.0007 | 1.3E-13 | LYPLAL1 | LOC643723(+175.5kb)\|LYPLAL1(+136.5kb) | 1 |
| rs78561351 | 1 | 227587047 | A | G | 0.0078 | 0.0007 | 5.2E-32 | CDC42BPA | CDC42BPA(+81.22kb)\|ZNF678(-164.2kb) | 0 |
| rs34392899 | 2 | 12878223 | A | C | 0.0036 | 0.0005 | 9.9E-12 | TRIB2 | LOC100506457(+159.7kb)\|MIR3125(+0.653kb)\|TRIB2(0) | 1 |
| rs4380183 | 2 | 19272693 | A | G | 0.0054 | 0.0007 | 3.5E-13 | MIR4757 | . | 0 |
| rs72778330 | 2 | 19394671 | A | G | -0.0175 | 0.0022 | 2.1E-15 | MIR4757 | MIR4757(-153.5kb)\|OSR1(-156.6kb) | 0 |
| rs11886621 | 2 | 28372900 | T | C | 0.0032 | 0.0005 | 1.8E-09 | BRE | BRE(0)\|LOC100505716(-157.7kb)\|MIR4263(+153.6kb) | 1 |
| rs3770876 | 2 | 36679028 | T | G | -0.0035 | 0.0006 | 6.9E-10 | CRIM1 | CRIM1(0)\|FEZ2(-100.4kb)\|LOC100288911(+96.32kb) | 1 |
| rs13009753 | 2 | 42544096 | A | G | 0.0042 | 0.0006 | 2.3E-11 | EML4 | COX7A2L(-33.55kb)\|EML4(0)\|KCNG3(-125.1kb)\|LOC102723824(+146.7kb)\|MTA3(-177.6kb) | 1 |
| rs1946545 | 2 | 56036435 | A | G | 0.0085 | 0.0007 | 5.4E-37 | EFEMP1 | EFEMP1(-56.66kb)\|MIR216A(-179.6kb)\|MIR216B(-191.4kb)\|MIR217(-173.7kb)\|PNPT1(+115.4kb)\|SMEK2(+191.6kb) | 0 |
| rs2946373 | 2 | 66551324 | T | C | -0.0035 | 0.0006 | 4.2E-09 | MIR4778 | MEIS1(-111.2kb)\|MEIS1-AS3(-99.15kb)\|MIR4778(-34.06kb) | 1 |
| rs1533296 | 2 | 111651503 | A | G | 0.0063 | 0.0006 | 8.8E-30 | ACOXL | ACOXL(0) | 0 |
| rs6434068 | 2 | 153357541 | C | G | 0.0032 | 0.0006 | 1.1E-08 | FMNL2 | FMNL2(0)\|PRPF40A(-150.6kb) | 0 |
| rs13017232 | 2 | 190218771 | T | C | 0.0036 | 0.0006 | 3E-09 | COL5A2 | COL5A2(+174.2kb)\|WDR75(-87.39kb) | 1 |
| rs72977807 | 2 | 228084531 | A | G | -0.0059 | 0.0011 | 2.5E-08 | COL4A3 | COL4A3(0)\|COL4A4(+55.26kb)\|LOC654841(-1.236kb)\|MFF(-105.3kb)\|TM4SF20(-142.3kb) | 1 |
| rs7581970 | 2 | 241930390 | T | G | -0.004 | 0.0006 | 5.1E-13 | SNED1 | AGXT(+111.9kb)\|ANO7(-197.5kb)\|C2orf54(+94.82kb)\|KIF1A(+170.7kb)\|LOC200772(+23.52kb)\|MTERFD2(-96.12kb)\|PASK(-115.1kb)\|PPP1R7(-158.6kb)\|SNED1(-7.864kb) | 1 |
| rs62228592 | 3 | 25187362 | A | G | 0.0042 | 0.0005 | 3.4E-14 | RARB | RARB(-28.46kb) | 0 |
| rs56131903 | 3 | 32879823 | A | T | 0.0048 | 0.0006 | 1.3E-16 | TRIM71 | CCR4(-113.2kb)\|CNOT10(+64.46kb)\|GLB1(-158.3kb)\|TRIM71(0) | 0 |
| rs13064784 | 3 | 49135961 | C | G | 0.0036 | 0.0005 | 4.8E-11 | QARS | ARIH2(+113kb)\|ARIH2OS(+179.1kb)\|C3orf62(-170.1kb)\|C3orf84(-79.11kb)\|CCDC36(-99.9kb)\|CCDC71(-64.01kb)\|DALRD3(+77.46kb)\|IMPDH2(+69.09kb)\|KLHDC8B(-73.06kb)\|LAMB2(-22.58kb)\|LAMB2P1(-54.33kb)\|MIR191(+77.82kb)\|MIR425(+78.29kb)\|MIR4271(-175.6kb)\|MIR6890(-1.325kb)\|NDUFAF3(+75.04kb)\|P4HTM(+91.38kb)\|QARS(0)\|QRICH1(+4.457kb)\|SLC25A20(+199.5kb)\|USP4(-178.6kb)\|USP19(-9.517kb)\|WDR6(+82.58kb) | 1 |
| rs57161291 | 3 | 58011419 | A | G | 0.0061 | 0.0006 | 1.7E-24 | FLNB | DNASE1L3(-166.9kb)\|FLNB(0)\|SLMAP(+96.52kb) | 0 |
| rs77320716 | 3 | 64573276 | T | C | -0.0062 | 0.001 | 8.4E-10 | ADAMTS9 | ADAMTS9(0)\|ADAMTS9-AS2(-97.27kb)\|LOC101929335(0)\|MIR548A2(-132.4kb) | 1 |

| | | | | | | | | | | |
|---|---|---|---|---|---|---|---|---|---|---|
| rs2364878 | 3 | 66840166 | T | G | 0.0037 | 0.0006 | 2.2E-10 | KBTBD8 | . | 0 |
| rs1463468 | 3 | 88389403 | A | G | -0.0055 | 0.0005 | 1.1E-23 | C3orf38 | C3orf38(+182.3kb)\|CGGBP1(+190.4kb)\|ZNF654(+195.6kb) | 0 |
| rs13076500 | 3 | 98943479 | T | C | 0.0058 | 0.0005 | 7.1E-27 | DCBLD2 | . | 0 |
| rs1383847 | 3 | 99093509 | T | C | -0.0064 | 0.0006 | 1.2E-25 | MIR548G | MIR548G(-179.6kb) | 0 |
| rs68092024 | 3 | 99691522 | T | C | 0.0031 | 0.0006 | 2.5E-08 | FILIP1L | CMSS1(0)\|COL8A1(+175.9kb)\|FILIP1L(0)\|HP09053(+148.8kb)\|MIR548G(0)\|MIR3921(+8.28kb) | 0 |
| rs142287284 | 3 | 100627655 | T | C | 0.006 | 0.0007 | 8.1E-18 | ABI3BP | ABI3BP(0)\|TFG(+159.8kb) | 0 |
| rs62255018 | 3 | 106103026 | A | G | -0.0032 | 0.0005 | 1.6E-09 | CBLB | . | 1 |
| rs9842074 | 3 | 146463174 | T | C | -0.0034 | 0.0005 | 4.1E-10 | PLSCR5 | PLSCR5(+139.2kb) | 1 |
| rs4696778 | 4 | 7919308 | C | G | -0.0043 | 0.0006 | 1.6E-13 | AFAP1 | ABLIM2(-47.73kb)\|AFAP1(0)\|AFAP1-AS1(+138.7kb)\|SORCS2(+174.7kb) | 1 |
| rs6842303 | 4 | 17854055 | T | G | 0.0035 | 0.0006 | 4.3E-08 | LCORL | DCAF16(+41.67kb)\|FAM184B(+70.92kb)\|LCORL(0)\|NCAPG(+7.568kb) | 1 |
| rs4864861 | 4 | 55097685 | T | C | -0.0061 | 0.0006 | 9.1E-22 | PDGFRA | CHIC2(+166.9kb)\|GSX2(+129.6kb)\|PDGFRA(0) | 0 |
| rs7677732 | 4 | 112825940 | A | G | 0.0046 | 0.0006 | 3.2E-15 | C4orf32 | . | 1 |
| rs1395236 | 4 | 126408988 | C | G | -0.0044 | 0.0006 | 2.7E-14 | FAT4 | FAT4(0)\|MIR2054(-19.42kb) | 0 |
| rs6825775 | 4 | 184695118 | C | G | -0.0042 | 0.0006 | 2E-12 | C4orf41 | RWDD4(+114.8kb)\|STOX2(-131.4kb)\|TRAPPC11(+60.37kb) | 1 |
| rs4866595 | 5 | 3662460 | T | C | 0.0039 | 0.0006 | 8.3E-11 | IRX1 | IRX1(+60.94kb)\|LINC01019(+126.3kb)\|LOC102467075(+158.3kb) | 0 |
| rs75905830 | 5 | 14291590 | A | G | 0.0059 | 0.001 | 1.6E-08 | TRIO | TRIO(0) | 1 |
| rs72759609 | 5 | 31952051 | T | C | 0.0091 | 0.0009 | 2.6E-24 | PDZD2 | GOLPH3(-172.8kb)\|MIR4279(+15.79kb)\|PDZD2(0) | 0 |
| rs11953967 | 5 | 52045824 | T | C | -0.0041 | 0.0005 | 6E-14 | PELO | ITGA1(-38.31kb)\|PELO(-37.95kb) | 0 |
| rs158653 | 5 | 55578661 | A | G | -0.0051 | 0.0005 | 1.5E-21 | ANKRD55 | ANKRD55(+49.48kb)\|LOC102467147(-175kb) | 0 |
| rs114852261 | 5 | 55690981 | A | G | -0.004 | 0.0007 | 3.7E-08 | ANKRD55 | ANKRD55(+161.8kb)\|LOC101928448(-116.2kb)\|LOC102467147(-62.64kb) | 0 |
| rs245809 | 5 | 55739061 | A | G | 0.0057 | 0.0007 | 3.7E-15 | ANKRD55 | LOC101928448(-68.16kb)\|LOC102467147(-14.56kb) | 0 |
| rs61275591 | 5 | 55775556 | A | G | 0.0078 | 0.0011 | 7.8E-14 | ANKRD55 | LOC101928448(-31.66kb)\|LOC102467147(0) | 0 |
| rs16892523 | 5 | 60867273 | C | G | -0.0064 | 0.0009 | 1.2E-13 | ZSWIM6 | C5orf64(-66.36kb)\|LOC100506526(-161.3kb)\|LOC101928651(-91.71kb)\|ZSWIM6(+25.27kb) | 0 |
| rs11746859 | 5 | 82770558 | A | G | 0.0033 | 0.0005 | 1.6E-09 | VCAN | HAPLN1(-163.5kb)\|VCAN(0)\|XRCC4(+121kb) | 0 |
| rs114270619 | 5 | 87834483 | A | C | 0.0059 | 0.001 | 2.4E-09 | LOC645323 | LINC00461(-2.113kb)\|LOC102546226(+99.58kb)\|MEF2C(-179.6kb)\|MIR9-2(-128.2kb)\|TMEM161B-AS1(+102kb) | 0 |
| rs396321 | 5 | 112113735 | T | C | 0.003 | 0.0005 | 1.4E-08 | APC | APC(0)\|DCP2(-198.7kb)\|LOC102467214(+147.1kb)\|LOC102467216(+95.15kb)\|REEP5(-98.34kb)\|SRP19(-83.15kb) | 1 |

| | | | | | | | | | | |
|---|---|---|---|---|---|---|---|---|---|---|
| rs34133486 | 5 | 125206660 | T | C | 0.0038 | 0.0006 | 2.7E-10 | GRAMD3 | . | 1 |
| rs7448395 | 5 | 128931357 | A | G | -0.0058 | 0.0007 | 1.9E-18 | ADAMTS 19 | ADAMTS19(0)\|KIAA1024L(-152.5kb)\|MIR4460(+198.5kb) | 0 |
| rs10075459 | 5 | 131541363 | C | G | 0.0039 | 0.0006 | 1.8E-12 | P4HA2 | ACSL6(+193.6kb)\|CSF2(+129.5kb)\|IL3(+142.5kb)\|LOC553103(-105.6kb)\|MIR3936(-159.8kb)\|MIR6830(-12.18kb)\|P4HA2(0)\|P4HA2-AS1(+12.86kb)\|PDLIM4(-51.99kb)\|SLC22A4(-88.78kb)\|SLC22A5(-164kb)\|=missense | 1 |
| rs78962119 | 5 | 133408702 | A | C | -0.0089 | 0.0009 | 6.8E-22 | TCF7 | C5orf15(+104.3kb)\|MIR3661(-152.7kb)\|PPP2CA(-123.4kb)\|SKP1(-83.38kb)\|TCF7(-41.7kb)\|VDAC1(+67.88kb) | 0 |
| rs4490572 | 5 | 148612322 | A | G | -0.003 | 0.0005 | 1.3E-08 | ABLIM3 | ABLIM3(0)\|AFAP1L1(-39.08kb)\|GRPEL2(-112.7kb)\|IL17B(-141.5kb)\|MIR143(-196.2kb)\|MIR143HG(-174.1kb)\|MIR145(-197.9kb)\|PCYOX1L(-125.2kb)\|SH3TC2(+169.6kb) | 1 |
| rs2348423 | 5 | 151760320 | A | G | 0.0035 | 0.0006 | 2.3E-09 | NMUR2 | CTB-12O2.1(+110.3kb)\|NMUR2(-10.78kb) | 1 |
| rs117012610 | 5 | 172198197 | A | G | -0.0131 | 0.0012 | 5.2E-26 | DUSP1 | DUSP1(0)\|ERGIC1(-63.02kb)\|LOC100268168(-183.6kb)\|LOC101928093(+8.632kb)\|NEURL1B(+79.66kb)\|RPL26L1(-188.2kb) | 0 |
| rs9504483 | 6 | 620222 | T | C | 0.0063 | 0.0008 | 2E-15 | EXOC2 | EXOC2(0)\|HUS1B(-35.72kb) | 0 |
| rs2745572 | 6 | 1548369 | A | G | 0.0041 | 0.0006 | 3.9E-12 | FOXC1 | FOXC1(-62.31kb)\|FOXF2(+152.5kb)\|GMDS(-75.66kb)\|MIR6720(+157.7kb) | 0 |
| rs6914444 | 6 | 1983440 | T | C | 0.0066 | 0.0008 | 5.6E-16 | GMDS | GMDS(0) | 0 |
| rs4960297 | 6 | 7212084 | T | C | 0.0063 | 0.0005 | 8.3E-33 | RREB1 | CAGE1(-114.8kb)\|RIOK1(-178kb)\|RREB1(0)\|SSR1(-69.29kb)\|=missense | 0 |
| rs9263750 | 6 | 31114655 | T | C | 0.0047 | 0.0007 | 2.1E-10 | CCHCR1 | C6orf15(+34.32kb)\|CCHCR1(0)\|CDSN(+26.4kb)\|DPCR1(+192.7kb)\|HCG22(+87kb)\|HCG27(-50.88kb)\|HLA-C(-121.9kb)\|MUC21(+157kb)\|MUC22(+111.5kb)\|POU5F1(-17.46kb)\|PSORS1C1(+6.786kb)\|PSORS1C2(+7.528kb)\|PSORS1C3(-26.86kb)\|TCF19(-11.65kb) | 1 |
| rs10947614 | 6 | 36573822 | C | G | 0.0065 | 0.0007 | 3E-23 | SRSF3 | CDKN1A(-70.41kb)\|CPNE5(-134.7kb)\|KCTD20(+114.9kb)\|MIR3925(-16.39kb)\|PANDAR(-67.58kb)\|PXT1(+163.2kb)\|RAB44(-91.8kb)\|SRSF3(+1.578kb)\|STK38(+58.58kb)\|=missense | 0 |
| rs4714260 | 6 | 39536622 | T | C | -0.004 | 0.0005 | 5.5E-14 | KIF6 | KIF6(0) | 0 |
| rs2744475 | 6 | 50784880 | C | G | -0.0038 | 0.0006 | 3E-10 | TFAP2B | TFAP2B(-1.558kb)\|TFAP2D(+44.13kb) | 1 |
| rs58657694 | 6 | 56567668 | A | C | 0.0039 | 0.0007 | 3.3E-09 | DST | DST(0)\|RNU6-71P(0) | 1 |
| rs1919875 | 6 | 122171052 | A | G | 0.0062 | 0.0009 | 4.7E-12 | GJA1 | . | 1 |
| rs2684249 | 6 | 122392511 | T | C | 0.0054 | 0.0005 | 2.3E-23 | HSF2 | . | 0 |
| rs1361108 | 6 | 126767600 | T | C | 0.0033 | 0.0006 | 8.3E-09 | CENPW | CENPW(0) | 1 |

| | | | | | | | | | | |
|---|---|---|---|---|---|---|---|---|---|---|
| rs12193446 | 6 | 129820038 | A | G | 0.0055 | 0.001 | 7.9E-09 | LAMA2 | ARHGAP18(-78.2kb)\|LAMA2(0) | 1 |
| rs7739956 | 6 | 148835642 | A | G | -0.0051 | 0.0009 | 3.3E-09 | SASH1 | SASH1(0) | 1 |
| rs1125 | 6 | 149979416 | A | G | -0.0037 | 0.0006 | 1.9E-10 | LATS1 | GINM1(+67.35kb)\|KATNA1(+9.476kb)\|LATS1(0)\|LRP11(-160.5kb)\|NUP43(-66.04kb)\|PCMT1(-91.41kb)\|PPIL4(+112.2kb)\|RPS18P9(+63.7kb)\|ZC3H12D(+173.3kb)\|=missense | 1 |
| rs4242276 | 6 | 151303497 | A | G | 0.0046 | 0.0006 | 1.2E-13 | MTHFD1L | MTHFD1L(0)\|PLEKHG1(+138.7kb) | 1 |
| rs3087749 | 7 | 4780514 | T | G | 0.0033 | 0.0005 | 1.3E-09 | FOXK1 | AP5Z1(-34.75kb)\|FOXK1(0)\|MIR4656(-47.68kb)\|MMD2(-151.4kb)\|PAPOLB(-116.9kb)\|RADIL(-58.22kb) | 1 |
| rs17166655 | 7 | 13086461 | T | C | 0.0048 | 0.0007 | 1.6E-10 | ARL4A | . | 1 |
| rs10260511 | 7 | 14237240 | A | C | 0.0076 | 0.0007 | 5.8E-27 | DGKB | DGKB(0) | 0 |
| rs57584385 | 7 | 19633159 | A | G | -0.0039 | 0.0006 | 5.5E-12 | TWISTNB | MIR3146(-111.8kb)\|TMEM196(-125.8kb)\|TWISTNB(-101.9kb) | 0 |
| rs7806362 | 7 | 28386703 | A | C | -0.004 | 0.0005 | 5E-13 | CREB5 | CREB5(0)\|JAZF1(+166.3kb)\|JAZF1-AS1(+105.7kb) | 0 |
| rs2282909 | 7 | 28844815 | T | G | 0.0042 | 0.0006 | 2.3E-11 | CREB5 | CPVL(-190.4kb)\|CREB5(0)\|LOC100506497(-174.8kb)\|TRIL(-148.2kb) | 0 |
| rs2072201 | 7 | 42117040 | A | T | 0.0034 | 0.0005 | 2E-10 | GLI3 | GLI3(0) | 1 |
| rs17855988 | 7 | 73474825 | C | G | 0.0052 | 0.0009 | 2.6E-08 | ELN | EIF4H(-113.9kb)\|ELN(0)\|LAT2(-149.3kb)\|LIMK1(-23.28kb)\|MIR590(-130.7kb)\|RFC2(-171kb)\|WBSCR28(+194.6kb) | 1 |
| rs10954741 | 7 | 83679565 | C | G | -0.0042 | 0.0007 | 4.8E-10 | SEMA3A | SEMA3A(0) | 1 |
| rs201515 | 7 | 101769164 | A | G | -0.0041 | 0.0005 | 9.4E-15 | CUX1 | CUX1(0)\|MIR4285(-167.2kb)\|SH2B2(-159.2kb) | 0 |
| rs10258576 | 7 | 115218932 | A | G | 0.0049 | 0.0007 | 6E-13 | TFEC | . | 1 |
| rs4236601 | 7 | 116162729 | A | G | 0.0044 | 0.0006 | 7.4E-14 | CAV1 | CAV1(-2.109kb)\|CAV2(+14.13kb)\|MET(-149.7kb)\|=missense | 1 |
| rs7782289 | 7 | 151091515 | A | G | -0.0031 | 0.0005 | 8.8E-09 | WDR86 | ABCF2(+167.1kb)\|CHPF2(+155.6kb)\|CRYGN(-35.54kb)\|MIR671(+155.9kb)\|MIR3907(-39.06kb)\|NUB1(+15.97kb)\|PRKAG2(-161.7kb)\|RHEB(-71.58kb)\|SMARCD3(+117.3kb)\|WDR86(0)\|WDR86-AS1(-14.73kb) | 1 |
| rs2976932 | 8 | 8255319 | T | C | -0.0077 | 0.0008 | 2.4E-20 | SGK223 | FAM86B3P(+152.9kb)\|SGK223(+15.98kb) | 0 |
| rs11782619 | 8 | 17526521 | T | C | -0.0041 | 0.0006 | 1.4E-12 | MTUS1 | FGL1(-195.4kb)\|MTUS1(0)\|PDGFRL(+25.88kb)\|SLC7A2(+98.44kb) | 1 |
| rs11777732 | 8 | 30248229 | T | C | 0.0035 | 0.0006 | 3.2E-08 | RBPMS | GTF2E2(-187.8kb)\|MIR548O2(+140kb)\|RBPMS(0)\|RBPMS-AS1(+5.312kb) | 1 |
| rs35631438 | 8 | 30424440 | T | C | 0.0042 | 0.0005 | 2.4E-15 | RBPMS | GSR(-111.1kb)\|GTF2E2(-11.59kb)\|RBPMS(0)\|RBPMS-AS1(+181.5kb)\|SMIM18(-71.68kb)\|UBXN8(-177.2kb) | 1 |
| rs7014542 | 8 | 37677118 | C | G | 0.0035 | 0.0006 | 3.5E-08 | GPR124 | ADRB3(-143.4kb)\|BRF2(-24.28kb)\|ERLIN2(+61.8kb)\|GOT1L1(-114.7kb)\|GPR124(0)\|LOC728024(+71.55k | 1 |

| | | | | | | | | | | |
|---|---|---|---|---|---|---|---|---|---|---|
| | | | | | | | | | b)\|PROSC(+39.83kb)\|RAB11FIP1(-39.35kb)\|ZNF703(+120.7kb) | |
| rs77289866 | 8 | 70097282 | T | C | -0.0043 | 0.0007 | 9.1E-11 | LOC100505718 | LOC100505718(+80.86kb) | 1 |
| rs12543430 | 8 | 72278010 | T | C | -0.0046 | 0.0006 | 3.2E-16 | EYA1 | EYA1(+3.543kb) | 0 |
| rs74421737 | 8 | 72578575 | T | C | 0.0054 | 0.0007 | 2.2E-15 | MSC | LOC100132891(-176.8kb)\|MSC(-175.2kb) | 0 |
| rs10957731 | 8 | 75522500 | A | C | 0.0035 | 0.0006 | 3.9E-10 | FLJ39080 | FLJ39080(0)\|MIR2052(-95.43kb)\|MIR5681A(+61.65kb)\|MIR5681B(+61.66kb) | 1 |
| rs56309454 | 8 | 76179452 | T | C | -0.0103 | 0.0017 | 8.3E-10 | CASC9 | CASC9(0) | 1 |
| rs11985148 | 8 | 77643348 | A | G | -0.0041 | 0.0007 | 8.1E-09 | ZFHX4 | ZFHX4(0)\|ZFHX4-AS1(+47.84kb) | 1 |
| rs10955403 | 8 | 88782176 | T | C | -0.0033 | 0.0005 | 7.3E-10 | DCAF4L2 | DCAF4L2(-100.8kb) | 1 |
| rs1448146 | 8 | 109331724 | A | T | 0.0042 | 0.0006 | 1.5E-11 | EIF3E | EIF3E(+70.76kb)\|EMC2(-124.1kb) | 1 |
| rs62525274 | 8 | 122803694 | T | C | -0.0042 | 0.0007 | 3.2E-09 | HAS2-AS1 | HAS2(+150.1kb)\|HAS2-AS1(+146.1kb) | 1 |
| rs4565471 | 8 | 131636781 | T | C | -0.003 | 0.0005 | 4.3E-08 | ADCY8 | ADCY8(-155.8kb)\|ASAP1(+180.9kb) | 0 |
| rs3008699 | 9 | 15913444 | T | C | -0.0035 | 0.0006 | 3.1E-08 | CCDC171 | CCDC171(0) | 1 |
| rs4961722 | 9 | 16529174 | T | C | 0.0037 | 0.0005 | 4.8E-12 | BNC2 | BNC2(0) | 1 |
| rs113551904 | 9 | 18106426 | T | C | -0.0142 | 0.0018 | 7.4E-15 | SH3GL2 | . | 0 |
| rs74744824 | 9 | 22044904 | A | G | -0.0158 | 0.0027 | 4.2E-09 | CDKN2B-AS1 | C9orf53(+77.15kb)\|CDKN2A(+50.41kb)\|CDKN2B(+35.59kb)\|CDKN2B-AS1(0)\|MTAP(+178.9kb) | 0 |
| rs944801 | 9 | 22051670 | C | G | 0.0163 | 0.0005 | 1.8E-197 | CDKN2B-AS1 | C9orf53(+83.92kb)\|CDKN2A(+57.18kb)\|CDKN2B(+42.36kb)\|CDKN2B-AS1(0)\|MTAP(+185.7kb) | 0 |
| rs10512176 | 9 | 89252706 | T | C | -0.004 | 0.0006 | 4.3E-11 | ZCCHC6 | . | 0 |
| rs56283543 | 9 | 89798284 | A | G | 0.0038 | 0.0006 | 1.5E-09 | C9orf170 | C9orf170(+23.64kb)\|LOC440173(+141.2kb)\|LOC494127(+98.14kb)\|LOC100506834(+181.3kb) | 1 |
| rs2472494 | 9 | 107695539 | T | C | 0.0038 | 0.0005 | 9.8E-13 | ABCA1 | ABCA1(+5.012kb)\|LOC286367(+155.5kb)\|NIPSNAP3A(+173.1kb)\|NIPSNAP3B(+159.2kb) | 1 |
| rs112493797 | 9 | 111713381 | T | C | 0.0078 | 0.0012 | 8.6E-11 | CTNNAL1 | ACTL7A(+87.35kb)\|ACTL7B(+95.11kb)\|CTNNAL1(0)\|FAM206A(+10.14kb)\|FRRS1L(-186.2kb)\|IKBKAP(+16.77kb)\|MIR32(-95.13kb)\|TMEM245(-64.03kb) | 1 |
| rs78116342 | 9 | 127574786 | A | G | 0.0055 | 0.001 | 1.8E-08 | OLFML2A | ARPC5L(-56.7kb)\|GOLGA1(-65.79kb)\|MIR181A2(+120kb)\|MIR181A2HG(+113.9kb)\|MIR181B2(+118.7kb)\|NR6A1(+41.2kb)\|OLFML2A(0)\|RPL35(-45.37kb)\|SCAI(-130.1kb)\|WDR38(-40.91kb) | 1 |
| rs12377624 | 9 | 129373110 | C | G | -0.0037 | 0.0006 | 2.9E-11 | LMX1B | LMX1B(-3.611kb)\|MVB12B(+103.8kb)\|ZBTB43(-194.2kb) | 1 |
| rs7020704 | 9 | 134593583 | A | C | 0.0033 | 0.0006 | 8.2E-09 | RAPGEF1 | MED27(-141.9kb)\|POMT1(+194.4kb)\|RAPGEF1(0)\|UCK1(+186.9kb) | 1 |

| | | | | | | | | | | |
|---|---|---|---|---|---|---|---|---|---|---|
| rs7467847 | 9 | 136115235 | C | G | 0.0105 | 0.0011 | 8.2E-22 | ABO | ABO(-15.33kb)\|ADAMTS13(-164.2kb)\|C9orf96(-128kb)\|CEL(+168kb)\|CELP(+152.8kb)\|GBGT1(+75.9kb)\|GTF3C5(+181.3kb)\|MED22(-92.52kb)\|MIR6877(+187.8kb)\|OBP2B(+30.6kb)\|RALGDS(+90.63kb)\|REXO4(-155.9kb)\|RPL7A(-99.83kb)\|SNORD24(-101kb)\|SNORD36A(-102.1kb)\|SNORD36B(-101.7kb)\|SNORD36C(-102.5kb)\|SURF1(-103.4kb)\|SURF2(-108.2kb)\|SURF4(-113.1kb)\|SURF6(-82.31kb) | 0 |
| rs190927291 | 10 | 21437861 | C | G | -0.0184 | 0.0023 | 1.3E-15 | NEBL | C10orf113(+2.373kb)\|NEBL(0)\|NEBL-AS1(-25.06kb) | 0 |
| rs4141671 | 10 | 60338753 | T | C | -0.0044 | 0.0005 | 6E-16 | BICC1 | BICC1(0)\|FAM133CP(-136kb)\|TFAM(+179.8kb) | 0 |
| rs1471246 | 10 | 62074139 | A | G | -0.0034 | 0.0005 | 3.8E-11 | ANK3 | ANK3(0) | 1 |
| rs2588924 | 10 | 63641670 | A | G | -0.0031 | 0.0005 | 4.6E-09 | ARID5B | ARID5B(-19.34kb)\|C10orf107(+115.6kb)\|MIR548AV(-94.39kb) | 1 |
| rs12775581 | 10 | 65382316 | T | G | -0.0037 | 0.0006 | 7.5E-10 | REEP3 | JMJD1C(+156.6kb)\|JMJD1C-AS1(+156kb)\|REEP3(0) | 1 |
| rs149176725 | 10 | 68991138 | C | G | 0.0098 | 0.0015 | 1.4E-11 | CTNNA3 | CTNNA3(0)\|LRRTM3(+129.8kb) | 0 |
| rs7916697 | 10 | 69991853 | A | G | -0.0102 | 0.0006 | 4.3E-63 | ATOH7 | ATOH7(0)\|DNA2(-182kb)\|HERC4(+156.8kb)\|HNRNPH3(-99.91kb)\|MYPN(+20.08kb)\|PBLD(-50.56kb)\|RUFY2(-109kb) | 0 |
| rs2569359 | 10 | 78587491 | A | G | 0.0054 | 0.0009 | 8.6E-09 | KCNMA1 | KCNMA1(-41.87kb) | 1 |
| rs17108260 | 10 | 94950713 | A | G | -0.0051 | 0.0005 | 1.6E-21 | CYP26A1 | CYP26A1(+113.1kb)\|CYP26C1(+122.3kb)\|EXOC6(+131.5kb)\|MYOF(-115.5kb) | 0 |
| rs2077218 | 10 | 96071561 | A | G | -0.005 | 0.0006 | 7.2E-15 | PLCE1 | NOC3L(-21.43kb)\|PLCE1(0)\|PLCE1-AS1(+24.73kb)\|TBC1D12(-90.62kb) | 0 |
| rs61873510 | 10 | 102626510 | T | G | -0.0034 | 0.0006 | 1.3E-08 | PAX2 | C10orf2(-120.8kb)\|FAM178A(-45.82kb)\|KAZALD1(-194.5kb)\|LZTS2(-130.4kb)\|MIR608(-108.2kb)\|MRPL43(-111.1kb)\|PAX2(+36.81kb)\|PDZD7(-140.9kb)\|SEMA4G(-105.8kb)\|SFXN3(-164.5kb) | 1 |
| rs11197820 | 10 | 118546046 | A | G | 0.0046 | 0.0005 | 1.1E-17 | HSPA12A | C10orf82(+116.6kb)\|ENO4(-62.98kb)\|HSPA12A(+43.96kb)\|KIAA1598(-96.84kb)\|PNLIPRP1(+177.4kb)\|PNLIPRP2(+141.4kb) | 0 |
| rs12225529 | 11 | 19948499 | A | G | 0.0034 | 0.0006 | 1.2E-09 | NAV2 | MIR4694(+166.9kb)\|NAV2(0) | 1 |
| rs1223090 | 11 | 31499705 | A | G | 0.0039 | 0.0005 | 4.6E-13 | IMMP1L | DCDC1(+108.3kb)\|DNAJC24(+45.32kb)\|ELP4(-31.57kb)\|IMMP1L(0) | 0 |
| rs2753411 | 11 | 33405001 | A | T | -0.0039 | 0.0005 | 8.5E-14 | HIPK3 | CSTF3-AS1(+191.9kb)\|HIPK3(+26.43kb)\|KIAA1549L(-158.9kb) | 0 |
| rs686489 | 11 | 57659832 | A | G | -0.0038 | 0.0006 | 4.7E-12 | TMX2-CTNND1 | BTBD18(+140.6kb)\|C11orf31(+148.9kb)\|CTNND1(+73.18kb)\|MED19(+180.2kb)\|OR6Q1(-138.6kb)\|OR9Q1(-131.5kb)\|TMX2(+151.4kb)\|TMX2-CTNND1(+73.18kb)\|ZDHHC5(+191.2kb) | 1 |
| rs61886103 | 11 | 63606309 | T | G | -0.0071 | 0.0011 | 4E-10 | MARK2 | ATL3(+166.9kb)\|C11orf84(+11.12kb)\|C11orf95(+70.2kb)\|COX8A(-135.8kb)\|MACROD1(-159.7kb)\|MARK2(- | 0 |

| | | | | | | | | | | |
|---|---|---|---|---|---|---|---|---|---|---|
| | | | | | | | | | 0.09kb)\|NAA40(-100.1kb)\|OTUB1(-147kb)\|RCOR2(-72.38kb)\|RTN3(+78.95kb) | |
| rs12789028 | 11 | 65326154 | A | G | -0.0091 | 0.0007 | 3.9E-41 | LTBP3 | EHBP1L1(-17.35kb)\|FAM89B(-13.66kb)\|FRMD8(+145.2kb)\|KAT5(-153.3kb)\|KCNK7(-34.17kb)\|LTBP3(+0.455kb)\|MALAT1(+52.22kb)\|MAP3K11(-39.07kb)\|MIR548AR(+43.33kb)\|MIR548BA(+43.32kb)\|MIR612(+114.1kb)\|MIR4489(-90.51kb)\|MIR4690(-77.63kb)\|NEAT1(+132.2kb)\|PCNXL3(-57.63kb)\|RELA(-94.91kb)\|RNASEH2C(-159kb)\|SCYL1(+19.97kb)\|SIPA1(-79.42kb)\|SLC25A45(+175kb)\|SSSCA1(-11.79kb)\|SSSCA1-AS1(-10.54kb) | 0 |
| rs11234891 | 11 | 86670842 | A | G | 0.0033 | 0.0006 | 1.2E-08 | LOC100506368 | FZD4(+4.402kb)\|LOC100506368(0)\|OR7E2P(+101.8kb)\|PRSS23(+148.6kb)\|TMEM135(-78.04kb) | 1 |
| rs7927570 | 11 | 86736525 | T | G | -0.0059 | 0.0007 | 1E-17 | TMEM135 | FZD4(+70.08kb)\|LOC100506368(+24.54kb)\|OR7E2P(+167.5kb)\|TMEM135(-12.36kb) | 0 |
| rs11021217 | 11 | 95292922 | A | G | -0.0047 | 0.0005 | 1.2E-17 | FAM76B | . | 0 |
| rs7123718 | 11 | 100645211 | C | G | -0.0049 | 0.0009 | 2.4E-08 | ARHGAP42 | ARHGAP42(0) | 1 |
| rs4937515 | 11 | 130268147 | C | G | 0.0076 | 0.0005 | 9.4E-46 | ADAMTS8 | ADAMTS8(-6.67kb)\|ADAMTS15(-50.72kb)\|ST14(+187.9kb)\|ZBTB44(+83.54kb) | 0 |
| rs1270640 | 11 | 130328867 | T | C | 0.012 | 0.0017 | 5.5E-12 | ADAMTS15 | ADAMTS8(+30.33kb)\|ADAMTS15(0)\|ZBTB44(+144.3kb) | 1 |
| rs5442 | 12 | 6954864 | A | G | 0.0082 | 0.0011 | 1.9E-13 | GNB3 | ACRBP(+198.3kb)\|ATN1(-78.76kb)\|C12orf57(-98.34kb)\|CD4(+24.89kb)\|CDCA3(-3.107kb)\|COPS7A(+113.8kb)\|DSTNP2(-38.98kb)\|EMG1(-125.1kb)\|ENO2(-68.75kb)\|GNB3(0)\|GPR162(+18.28kb)\|ING4(+182.6kb)\|LAG3(+67.24kb)\|LEPREL2(+5.846kb)\|LPCAT3(-130.5kb)\|LRRC23(-59.03kb)\|MIR141(-118.4kb)\|MIR200C(-118kb)\|MLF2(+92.23kb)\|PHB2(-119.6kb)\|PIANP(+144.9kb)\|PTMS(+74.75kb)\|PTPN6(-100.9kb)\|RPL13P5(-38.28kb)\|SCARNA12(-121.6kb)\|SPSB2(-25.24kb)\|TPI1(-21.72kb)\|USP5(-6.42kb)\|ZNF384(+156.1kb)\|=MISSENSE | 1 |
| rs7977499 | 12 | 20047676 | T | C | -0.0041 | 0.0008 | 4.8E-08 | LOC100506393 | LOC100506393(-119.9kb) | 1 |
| rs16930371 | 12 | 26392080 | A | G | 0.006 | 0.0007 | 2.1E-18 | SSPN | BHLHE41(+114.1kb)\|ITPR2(-96.2kb)\|RASSF8(+159.3kb)\|SSPN(+4.372kb) | 0 |
| rs7307355 | 12 | 43809193 | T | G | -0.0033 | 0.0005 | 1.3E-09 | ADAMTS20 | ADAMTS20(0) | 1 |
| rs12818241 | 12 | 48157019 | T | C | -0.0057 | 0.0007 | 5.1E-15 | RAPGEF3 | ENDOU(+37.66kb)\|HDAC7(-19.49kb)\|RAPGEF3(+4.13kb)\|RPAP3(+57.18kb)\|SLC48A1(-9.947kb)\|VDR(-78.3kb)\|=missense | 0 |
| rs61927933 | 12 | 53369508 | T | C | -0.0032 | 0.0006 | 9.9E-09 | KRT18 | CSAD(-181.9kb)\|EIF4B(-30.55kb)\|IGFBP6(-121.9kb)\|KRT3(+179.6kb)\|KRT4(+161.6kb)\|KRT8(+25.86kb)\|KRT18(+22.82kb)\|KRT76(+198.4kb)\|KRT78(+126.7kb)\|KRT79(+141.4kb)\|LOC283335(-67.46kb)\|MIR6757(-81.22kb)\|SOAT2(-127.8kb)\|SPRYD3(-88.59kb)\|TENC1(-71.3kb) | 1 |

| | | | | | | | | | | |
|---|---|---|---|---|---|---|---|---|---|---|
| rs6582298 | 12 | 76114872 | A | G | -0.0036 | 0.0006 | 1.3E-10 | KRR1 | . | 0 |
| rs1397893 | 12 | 84024485 | T | C | -0.0119 | 0.0005 | 7.3E-110 | TMTC2 | . | 0 |
| rs111522534 | 12 | 84701253 | A | G | 0.0104 | 0.0018 | 4.3E-09 | SLC6A15 | . | 0 |
| rs9651957 | 12 | 107170128 | T | C | 0.0046 | 0.0005 | 8.6E-17 | RIC8B | LOC100287944(+1.519kb)\|LOC100505978(+91.65kb)\|RFX4(+13.55kb)\|RIC8B(0)\|TMEM263(-179.4kb) | 0 |
| rs11057488 | 12 | 124665773 | A | G | -0.0052 | 0.0005 | 3E-22 | ZNF664-FAM101A | FAM101A(-107.9kb)\|MIR6880(-156kb)\|NCOR2(-143.2kb)\|ZNF664(+165.8kb)\|ZNF664-FAM101A(0) | 0 |
| rs17081882 | 13 | 25755286 | T | C | -0.0066 | 0.0009 | 2.7E-14 | FAM123A | AMER2(+8.865kb)\|ATP8A2(-190.9kb)\|LOC101928922(+1.147kb)\|LOC102723318(-0.183kb)\|MTMR6(-65.05kb)\|NUPL1(-120.4kb)\|PABPC3(+82.58kb) | 1 |
| rs9527463 | 13 | 33935146 | T | C | -0.0034 | 0.0006 | 1.6E-09 | STARD13 | STARD13(0)\|STARD13-AS(+79.68kb) | 1 |
| rs9546383 | 13 | 36683268 | T | C | -0.0078 | 0.0006 | 2.3E-37 | DCLK1 | CCDC169(-117.9kb)\|CCDC169-SOHLH2(-59.08kb)\|DCLK1(0)\|MIR548F5(+167.9kb)\|SOHLH2(-59.08kb)\|SPG20(-192.5kb) | 0 |
| rs4942561 | 13 | 47209347 | T | G | 0.0036 | 0.0006 | 1.8E-08 | LRCH1 | ESD(-136kb)\|HTR2A(-196.3kb)\|LOC101929344(+168.4kb)\|LRCH1(0) | 1 |
| rs1028727 | 13 | 51900442 | T | C | -0.005 | 0.0008 | 2.8E-11 | SERPINE3 | FAM124A(+42.06kb)\|INTS6(-35.26kb)\|INTS6-AS1(-127kb)\|LINC00371(+153.9kb)\|MIR5693(-22.26kb)\|SERPINE3(-14.72kb) | 1 |
| rs9559368 | 13 | 109223792 | T | C | 0.006 | 0.0006 | 2.8E-23 | MYO16 | MYO16(-24.71kb) | 0 |
| rs4238271 | 13 | 110769947 | A | C | 0.0061 | 0.0005 | 9.3E-30 | COL4A1 | COL4A1(-31.36kb)\|COL4A2(-189.7kb) | 0 |
| rs3811183 | 14 | 23452128 | C | G | -0.0055 | 0.0005 | 4.2E-25 | JUB | ACIN1(-75.64kb)\|AJUBA(+0.277kb)\|C14orf93(-3.981kb)\|C14orf119(-112.6kb)\|CDH24(-64.14kb)\|CEBPE(-134.4kb)\|HAUS4(+25.78kb)\|LOC101926933(+28.23kb)\|LRP10(+104.8kb)\|MIR4707(+25.89kb)\|MMP14(+135.3kb)\|MRPL52(+147.9kb)\|PRMT5(+53.33kb)\|PSMB5(-42.93kb)\|PSMB11(-59.25kb)\|RBM23(+63.73kb)\|REM2(+95.24kb)\|SLC7A7(+163.1kb)\|SLC7A8(-142.4kb) | 0 |
| rs2251171 | 14 | 53989447 | A | G | -0.0053 | 0.0005 | 6.2E-22 | DDHD1 | . | 0 |
| rs4898820 | 14 | 54427057 | T | G | -0.0046 | 0.0005 | 5.5E-17 | BMP4 | BMP4(+3.503kb)\|MIR5580(+11.86kb) | 0 |
| rs34935520 | 14 | 61091401 | A | G | -0.0106 | 0.0005 | 2.6E-86 | SIX1 | C14orf39(+138.6kb)\|MNAT1(-110.1kb)\|SIX1(-20.02kb)\|SIX4(-84.85kb)\|SIX6(+112.9kb) | 0 |
| rs28761289 | 14 | 65113768 | A | G | -0.0059 | 0.0007 | 7.8E-18 | PLEKHG3 | AKAP5(+172.5kb)\|HSPA2(+103.8kb)\|LOC102723809(+106.7kb)\|MIR548AZ(+186.3kb)\|MIR7855(-138.6kb)\|MTHFD1(+187kb)\|PLEKHG3(-57.42kb)\|PPP1R36(+57.67kb)\|SPTB(-99.23kb)\|ZBTB1(+113.4kb)\|ZBTB25(+143.2kb) | 0 |
| rs11628885 | 14 | 68724790 | A | G | 0.0041 | 0.0007 | 4.9E-08 | RAD51L1 | RAD51B(0) | 1 |

| | | | | | | | | | | |
|---|---|---|---|---|---|---|---|---|---|---|
| rs1957330 | 14 | 85864225 | T | C | 0.0037 | 0.0006 | 1.7E-09 | FLRT2 | FLRT2(-132.3kb)\|LINC00911(0) | 0 |
| rs7160530 | 14 | 86010805 | A | G | -0.0047 | 0.0006 | 1.9E-13 | FLRT2 | FLRT2(0)\|LINC00911(+124.4kb) | 0 |
| rs1815237 | 14 | 91881908 | T | G | -0.0031 | 0.0005 | 7.4E-09 | CCDC88C | C14orf159(+190.2kb)\|CATSPERB(-165.2kb)\|CCDC88C(0)\|GPR68(+161.7kb)\|SMEK1(-41.92kb) | 1 |
| rs2249954 | 14 | 92383999 | T | C | -0.0065 | 0.0011 | 1E-08 | FBLN5 | ATXN3(-140.9kb)\|CATSPERB(+185.6kb)\|FBLN5(0)\|NDUFB1(-198.5kb)\|TC2N(+50.12kb)\|TRIP11(-50.24kb) | 1 |
| rs11160251 | 14 | 95957694 | T | G | 0.0033 | 0.0006 | 2.3E-08 | C14orf49 | CLMN(+171.4kb)\|GLRX5(-43.63kb)\|LINC00341(+81.27kb)\|LOC101929080(+155.9kb)\|SCARNA13(-42kb)\|SNHG10(-41.55kb)\|SYNE3(+15.52kb)\|TCL1B(-195.1kb)\|TCL6(-159.8kb) | 1 |
| rs589135 | 15 | 35001442 | A | G | -0.0039 | 0.0006 | 4.6E-12 | GJD2 | ACTC1(-78.85kb)\|AQR(-147.1kb)\|GJD2(-43.2kb)\|GOLGA8B(+125.7kb)\|LOC101928174(-45.84kb)\|MIR1233-1(+180.9kb)\|MIR1233-2(+180.9kb) | 1 |
| rs79404983 | 15 | 71841635 | T | C | -0.0043 | 0.0007 | 2.6E-09 | THSD4 | LOC101929173(-139.4kb)\|THSD4(0) | 0 |
| rs893819 | 15 | 74229524 | A | G | -0.005 | 0.0006 | 2.1E-19 | LOXL1 | C15orf59(+185.7kb)\|GOLGA6A(-132.7kb)\|ISLR2(-192.2kb)\|LOC283731(-189.2kb)\|LOC101929221(+167.4kb)\|LOXL1(0)\|LOXL1-AS1(+8.935kb)\|PML(-57.49kb)\|STOML1(-46.03kb)\|TBC1D21(+47.97kb) | 0 |
| rs13380109 | 15 | 79378775 | A | G | 0.0029 | 0.0005 | 2.9E-08 | RASGRF1 | ANKRD34C(-196.4kb)\|CTSH(+141.4kb)\|LOC729911(-105.3kb)\|MIR184(-123.4kb)\|MORF4L1(+188.7kb)\|RASGRF1(0) | 1 |
| rs59199978 | 15 | 84484384 | A | G | -0.0039 | 0.0007 | 1.9E-08 | ADAMTSL3 | ADAMTSL3(0)\|SH3GL3(+196.9kb) | 1 |
| rs11638313 | 15 | 96723448 | T | C | -0.0036 | 0.0006 | 6.4E-09 | NR2F2 | MIR1469(-153kb)\|NR2F2(-145.7kb)\|NR2F2-AS1(-86.17kb) | 1 |
| rs28612945 | 15 | 99458902 | T | C | -0.0047 | 0.0006 | 7.3E-13 | IGF1R | IGF1R(0)\|MIR4714(+131.2kb)\|PGPEP1L(-52.56kb)\|SYNM(-186.4kb) | 0 |
| rs34222435 | 15 | 101200873 | T | C | 0.0094 | 0.0008 | 2.8E-32 | ASB7 | ASB7(+8.969kb)\|CERS3(+115.9kb)\|LINS(+58.43kb)\|PRKXP1(+101.4kb) | 0 |
| rs6598413 | 15 | 101551869 | A | T | 0.0037 | 0.0006 | 3.7E-09 | LRRK1 | ALDH1A3(+95.04kb)\|CHSY1(-164.1kb)\|LRRK1(0) | 1 |
| rs28731492 | 15 | 101749434 | A | G | -0.0039 | 0.0006 | 2.2E-10 | CHSY1 | CHSY1(0)\|LOC100507472(-98.02kb)\|LRRK1(+139.1kb)\|PCSK6(-94.7kb)\|SNRPA1(-72.28kb)\|VIMP(-61.68kb) | 1 |
| rs2516740 | 16 | 2097110 | A | C | -0.0034 | 0.0006 | 3.4E-08 | NTHL1 | BRICD5(-162.1kb)\|CASKIN1(-130.1kb)\|DNASE1L2(-189.4kb)\|E4F1(-176.4kb)\|ECI1(-192.8kb)\|GFER(+59.36kb)\|HS3ST6(+128.9kb)\|LINC00254(+162.9kb)\|MEIOB(+174.9kb)\|MIR1225(-43.08kb)\|MIR3180-5(-88.87kb)\|MIR4516(-86.01kb)\|MIR6511B1(-59.56kb)\|MLST8(-158.1kb)\|MSRB1(+103.8kb)\|NDUFB10(+85.13kb)\|NOXO1(+65.56kb)\|NPW(+26.35kb)\|NTHL1(0)\|PGP(-164.5kb)\|PKD1(-41.6kb)\|RAB26(-101.5kb)\|RNF151(+78.13kb)\|RPL3L(+92.43kb)\|RPS2(+82.28kb)\|SLC9A3R2(+8.083k | 1 |

| | | | | | | | | | | |
|---|---|---|---|---|---|---|---|---|---|---|
| | | | | | | | | | b)\|SNHG9(+81.6kb)\|SNORA10(+84.64kb)\|SNORA64(+84kb)\|SNORA78(+81.8kb)\|SNORD60(-107.9kb)\|SYNGR3(+52.83kb)\|TBL3(+68.36kb)\|TRAF7(-108.7kb)\|TSC2(-0.879kb)\|ZNF598(+37.29kb) | |
| rs14347 | 16 | 15133889 | T | G | -0.0033 | 0.0006 | 1.5E-08 | NTAN1 | LOC100288162(+112.7kb)\|MIR3179-1(+138.4kb)\|MIR3179-2(+138.4kb)\|MIR3179-3(+138.4kb)\|MIR3180-1(+128.7kb)\|MIR3180-2(+128.7kb)\|MIR3180-3(+128.7kb)\|MIR3180-4(-114.8kb)\|MIR6511A2(+114kb)\|MIR6511B1(-94.03kb)\|MIR6770-2(+109.2kb)\|NOMO1(+143.9kb)\|NPIPA1(+87.96kb)\|NTAN1(0)\|PDXDC1(0)\|RRN3(-19.99kb) | 1 |
| rs11645288 | 16 | 51172677 | A | G | -0.0041 | 0.0007 | 5.1E-10 | SALL1 | LOC101927334(+103kb)\|SALL1(0) | 0 |
| rs8059770 | 16 | 51394394 | A | G | 0.0061 | 0.0006 | 3.9E-24 | SALL1 | . | 0 |
| rs71386552 | 16 | 51412230 | C | G | -0.0045 | 0.0007 | 1.3E-09 | SALL1 | . | 0 |
| rs1420993 | 16 | 51464898 | C | G | -0.0094 | 0.0006 | 1.4E-58 | SALL1 | . | 0 |
| rs28514893 | 16 | 51573119 | T | C | 0.007 | 0.0007 | 7.6E-26 | SALL1 | . | 0 |
| rs11865100 | 16 | 74404812 | T | C | -0.0063 | 0.0009 | 1.5E-13 | LOC283922 | CLEC18B(-37.72kb)\|GLG1(-76.51kb)\|LOC283922(+2.659kb)\|LOC101928035(+155.4kb)\|PSMD7(+64.63kb) | 1 |
| rs75673420 | 16 | 77553790 | A | G | -0.0056 | 0.001 | 1.6E-08 | ADAMTS18 | ADAMTS18(+84.78kb) | 1 |
| rs36008565 | 16 | 84751046 | C | G | 0.0034 | 0.0006 | 4.3E-08 | USP10 | COTL1(+99.34kb)\|CRISPLD2(-102.5kb)\|KLHL36(+55.13kb)\|USP10(0) | 1 |
| rs12447168 | 16 | 85502220 | A | C | -0.0032 | 0.0006 | 4.2E-08 | KIAA0182 | GSE1(-142.8kb)\|LINC00311(+180.5kb)\|MIR5093(+162.3kb)\|MIR7851(-157.6kb) | 1 |
| rs1687626 | 16 | 86382120 | A | G | 0.0036 | 0.0006 | 8.9E-11 | LOC732275 | FENDRR(-126kb)\|FOXF1(-162kb)\|LINC00917(+2.835kb)\|LINC01081(+62.27kb)\|LINC01082(+148.8kb)\|LOC146513(+55.12kb)\|MTHFSD(-181.7kb) | 0 |
| rs17273989 | 16 | 86466387 | T | C | -0.0154 | 0.0013 | 8.8E-31 | LOC400550 | FENDRR(-41.74kb)\|FLJ30679(-122.5kb)\|FOXC2(-134.5kb)\|FOXF1(-77.74kb)\|FOXL1(-145.7kb)\|LINC00917(+87.1kb)\|LINC01081(+146.5kb)\|LOC146513(+139.4kb)\|MTHFSD(-97.39kb) | 0 |
| rs4843969 | 16 | 86531602 | T | G | -0.0032 | 0.0005 | 5.7E-10 | LOC400550 | FENDRR(0)\|FLJ30679(-57.32kb)\|FOXC2(-69.25kb)\|FOXF1(-12.53kb)\|FOXL1(-80.51kb)\|LINC00917(+152.3kb)\|MTHFSD(-32.18kb) | 0 |
| rs116875733 | 17 | 10033214 | A | G | -0.0054 | 0.0008 | 5.9E-12 | GAS7 | GAS7(0)\|MYH13(-171kb) | 1 |
| rs8078686 | 17 | 45735706 | T | C | 0.0036 | 0.0005 | 5.8E-11 | KPNB1 | KPNB1(0)\|LRRC46(-173.3kb)\|MRPL10(-164.9kb)\|MRPL45P2(+165.7kb)\|NPEPPS(+35.06kb)\|OSBPL7(-149kb)\|SCRN2(-179.3kb)\|SP6(-186.6kb)\|TBKBP1(-36.92kb)\|TBX21(-74.9kb) | 1 |
| rs6504608 | 17 | 47424681 | A | C | -0.0038 | 0.0006 | 1E-11 | ZNF652 | ABI3(+124.1kb)\|B4GALNT2(+177.3kb)\|FLJ40194(+88.65kb)\|GNGT2(+136.7kb)\|LOC100288866(-158.3kb)\|LOC101927207(-110.5kb)\|LOC102724596(- | 1 |

| | | | | | | | | | | |
|---|---|---|---|---|---|---|---|---|---|---|
| | | | | | | | | | 13.84kb)\|MIR6129(+58.86kb)\|MIR6165(-163.5kb)\|NGFR(-148kb)\|PHB(-56.73kb)\|PHOSPHO1(+116.6kb)\|ZNF652(0) | |
| rs4794104 | 17 | 48225686 | C | G | -0.0065 | 0.0007 | 2.3E-19 | PPP1R9B | COL1A1(-35.77kb)\|DLX3(+153.1kb)\|DLX4(+173.4kb)\|HILS1(-23.1kb)\|ITGA3(+57.84kb)\|LOC284080(+92.58kb)\|LOC101927230(-66.37kb)\|PDK2(+36.95kb)\|PPP1R9B(0)\|SAMD14(+18.44kb)\|SGCA(-17.68kb)\|TMEM92(-123.1kb)\|XYLT2(-197.7kb) | 0 |
| rs9915128 | 17 | 55533005 | T | C | 0.0035 | 0.0006 | 9.9E-10 | MSI2 | LOC101927539(-145.2kb)\|LOC101927557(-66.6kb)\|MSI2(0) | 1 |
| rs1558259 | 17 | 59265918 | A | G | -0.0077 | 0.0007 | 9.1E-31 | BCAS3 | BCAS3(0) | 0 |
| rs1420791 | 17 | 63914750 | A | G | -0.0088 | 0.0011 | 5.5E-17 | CEP112 | CEP112(0) | 1 |
| rs4789729 | 17 | 80146089 | A | G | -0.0032 | 0.0006 | 3.4E-08 | CCDC57 | ASPSCR1(+170.8kb)\|CCDC57(0)\|CD7(-126.7kb)\|CSNK1D(-54.45kb)\|DCXR(+150.5kb)\|DUS1L(+122.4kb)\|FASN(+89.98kb)\|GPS1(+130.7kb)\|LRRC45(+157.1kb)\|MIR6787(-48.45kb)\|RAC3(+154kb)\|RFNG(+136.4kb)\|SECTM1(-132.8kb)\|SLC16A3(-40.19kb)\|STRA13(+165.3kb)\|TEX19(-171kb)\|UTS2R(-186.1kb) | 1 |
| rs11664106 | 18 | 2846812 | A | T | -0.0035 | 0.0006 | 3.2E-09 | EMILIN2 | CBX3P2(+191.4kb)\|EMILIN2(-0.215kb)\|LOC727896(-96.4kb)\|LPIN2(-70.18kb)\|SMCHD1(+41.8kb) | 1 |
| rs568267 | 18 | 8799828 | T | C | -0.0037 | 0.0006 | 5.9E-09 | CCDC165 | MTCL1(0)\|RAB12(+160.4kb) | 1 |
| rs11080508 | 18 | 11283857 | T | C | 0.0043 | 0.0007 | 1.4E-09 | FAM38B | PIEZO2(+135.1kb) | 1 |
| rs55696318 | 18 | 60226704 | A | T | 0.0037 | 0.0006 | 6.1E-09 | ZCCHC2 | PHLPP1(-156kb)\|TNFRSF11A(+171.8kb)\|ZCCHC2(0) | 1 |
| rs123698 | 19 | 807442 | C | G | -0.005 | 0.0006 | 1.4E-19 | PTBP1 | ARID3A(-118.6kb)\|AZU1(-20.39kb)\|CFD(-52.22kb)\|ELANE(-44.85kb)\|FGF22(+163.8kb)\|FSTL3(+124kb)\|GRIN3B(-193kb)\|HCN2(+190.3kb)\|KISS1R(-109.9kb)\|LPPR3(-5.045kb)\|MED16(-60.52kb)\|MIR3187(-6.141kb)\|MIR4745(+2.441kb)\|MISP(+43.12kb)\|PALM(+59.11kb)\|POLRMT(+173.9kb)\|PRSS57(+112kb)\|PRTN3(-33.54kb)\|PTBP1(0)\|R3HDM4(-89.06kb)\|RNF126(+144.2kb)\|WDR18(-176.9kb) | 0 |
| rs8102936 | 19 | 32027330 | A | G | -0.0046 | 0.0006 | 2.5E-14 | THEG5 | THEG5(-51.76kb)\|TSHZ3(+187.1kb) | 0 |
| rs311384 | 19 | 47455315 | A | G | 0.0039 | 0.0006 | 3.6E-11 | ARHGAP35 | AP2S1(+101.1kb)\|ARHGAP35(0)\|FKRP(+193.5kb)\|NPAS1(-68.83kb)\|SAE1(-178.8kb)\|SLC1A5(+163.5kb)\|SNAR-E(+121.4kb)\|TMEM160(-93.85kb)\|ZC3H4(-112.1kb) | 1 |
| rs6140010 | 20 | 6473123 | A | G | 0.0117 | 0.0006 | 4.8E-97 | BMP2 | CASC20(0) | 0 |
| rs6038531 | 20 | 6514692 | A | G | 0.0182 | 0.0021 | 7.1E-18 | BMP2 | CASC20(+5.586kb) | 0 |
| rs173106 | 20 | 6743306 | A | C | -0.0045 | 0.0006 | 1.3E-14 | BMP2 | BMP2(-5.438kb) | 0 |

| | | | | | | | | | | |
|---|---|---|---|---|---|---|---|---|---|---|
| rs6141836 | 20 | 31465932 | A | G | 0.0057 | 0.0007 | 3.3E-15 | MAPRE1 | BPIFB2(-129.5kb)\|BPIFB3(-<br>177.3kb)\|BPIFB6(-<br>153.5kb)\|COMMD7(+134.1kb)\|DNMT3B(+<br>68.77kb)\|MAPRE1(+27.72kb)\|SUN5(-<br>105.6kb) | 0 |
| rs2073325 | 20 | 31669990 | T | C | -0.0035 | 0.0006 | 2.3E-10 | C20orf18<br>6 | BPIFA1(-153.8kb)\|BPIFA2(-<br>85.97kb)\|BPIFA3(-135.1kb)\|BPIFA4P(-<br>111.4kb)\|BPIFB2(+58.48kb)\|BPIFB3(+8.55<br>6kb)\|BPIFB4(0)\|BPIFB6(+38.14kb)\|SUN5(<br>+77.75kb)\|=missense | 1 |
| rs1105402 | 20 | 45781924 | T | C | 0.0033 | 0.0005 | 7.8E-10 | EYA2 | EYA2(0)\|LOC100131496(-<br>165.3kb)\|MIR3616(-13.68kb)\|ZMYND8(-<br>55.93kb) | 1 |
| rs4822983 | 22 | 29115066 | T | C | -0.0115 | 0.0006 | 1.3E-88 | CHEK2 | CCDC117(-53.6kb)\|CHEK2(0)\|HSCB(-<br>22.98kb)\|TTC28(+39.21kb)\|XBP1(-<br>75.48kb)\|ZNRF3(-164.7kb) | 0 |
| rs5762919 | 22 | 29363344 | T | C | 0.0105 | 0.0014 | 3.1E-14 | ZNRF3 | C22orf31(-<br>91.32kb)\|CCDC117(+178.1kb)\|KREMEN1(<br>-<br>105.7kb)\|XBP1(+166.8kb)\|ZNRF3(0)\|ZNRF<br>3-AS1(-57.64kb) | 0 |
| rs713875 | 22 | 30592487 | C | G | 0.0059 | 0.0005 | 9.2E-29 | HORMAD<br>2 | CCDC157(-160.1kb)\|GATSL3(-<br>88.62kb)\|HORMAD2(+19.42kb)\|KIAA1656(<br>-172.3kb)\|LIF(-<br>43.95kb)\|LOC101929664(+116kb)\|MIR681<br>8(+189.4kb)\|MTMR3(+165.6kb)\|OSM(-<br>66.33kb)\|RNF215(-182.3kb)\|SF3A1(-<br>135.5kb)\|TBC1D10A(-95.49kb) | 0 |
| rs73166584 | 22 | 30649229 | T | C | 0.0049 | 0.0008 | 8.3E-11 | LIF | CCDC157(-103.4kb)\|GATSL3(-<br>31.88kb)\|HORMAD2(+76.17kb)\|KIAA1656(<br>-<br>115.6kb)\|LIF(+6.389kb)\|LOC101929664(+<br>172.8kb)\|MTFP1(-172.4kb)\|OSM(-<br>9.589kb)\|RNF215(-125.6kb)\|SEC14L2(-<br>143.7kb)\|SF3A1(-78.75kb)\|TBC1D10A(-<br>38.75kb) | 1 |
| rs8140671 | 22 | 37910591 | A | G | 0.0066 | 0.0007 | 1.8E-23 | CARD10 | CARD10(0)\|CDC42EP1(-<br>45.88kb)\|CYTH4(+199.2kb)\|ELFN2(+87.09<br>kb)\|GGA1(-93.89kb)\|LGALS1(-<br>161kb)\|LGALS2(-<br>55.66kb)\|LOC100506271(+159.7kb)\|LOC1<br>01927051(-<br>127.2kb)\|MFNG(+28.11kb)\|NOL12(-<br>171.8kb)\|PDXP(-144.1kb)\|SH3BP1(-<br>125.1kb)\|TRIOBP(-182.4kb) | 0 |
| rs4396807 | 22 | 38138379 | C | G | 0.0065 | 0.0006 | 5.7E-31 | TRIOBP | ANKRD54(-<br>88.48kb)\|CDC42EP1(+173kb)\|EIF3L(-<br>107kb)\|GALR3(-81.01kb)\|GCAT(-<br>65.53kb)\|GGA1(+108.8kb)\|H1F0(-<br>62.73kb)\|LGALS1(+62.57kb)\|LGALS2(+16<br>2.4kb)\|LOC101927051(+84kb)\|MICALL1(-<br>163.8kb)\|MIR658(-101.9kb)\|MIR659(-<br>105.3kb)\|NOL12(+48.89kb)\|PDXP(+75.44k<br>b)\|SH3BP1(+86.33kb)\|TRIOBP(0) | 0 |
| rs12558081 | 23 | 43939978 | A | G | -0.0035 | 0.0006 | 9.7E-09 | EFHC2 | EFHC2(-<br>67.15kb)\|MAOB(+198.3kb)\|NDP(+107.1kb) | 0 |

#### Table S3. 231 lead genome-wide significant independent SNPs for vertical disc diameter from the meta-analysis of UKB, CLSA, and IGGC (European ancestry).

Abbreviations: SNP, single nucleotide polymorphism. CHR, Chromosome; BP, position, A1, effect allele; A2, non-effect allele; SE, Standard error; P, p value; novel, “1” novel SNPs for vertical disc diameter or disc area.

| SNP | CHR | BP | A1 | A2 | BETA | SE | P | Nearest gene name | Genes within 200kb | novel |
|---|---|---|---|---|---|---|---|---|---|---|
| rs12741594 | 1 | 3053957 | A | T | -0.0421 | 0.0034 | 5.7E-36 | PRDM16 | ACTRT2(+114.5kb)\|LINC00982(+69.67kb)\|MIR4251(+9.358kb)\|PRDM16(0) | 0 |
| rs75978447 | 1 | 23510830 | A | G | -0.0146 | 0.0021 | 2.3E-12 | HTR1D | C1orf213(-184.6kb)\|C1orf234(+168.5kb)\|HNRNPR(-125.4kb)\|HTR1D(-7.557kb)\|KDM1A(+100.6kb)\|LOC100996511(-96.97kb)\|LUZP1(+15.48kb)\|MIR3115(+140kb)\|TCEA3(-196.7kb)\|ZNF436(-175.1kb) | 1 |
| rs3768321 | 1 | 40035928 | T | G | -0.0128 | 0.0021 | 4.9E-10 | PABPC4 | BMP8A(+40.39kb)\|BMP8B(-188kb)\|HEYL(-53.17kb)\|HPCAL4(-108.4kb)\|KIAA0754(+153.8kb)\|MACF1(+83.12kb)\|NT5C1A(-88.86kb)\|OXCT2(-199.3kb)\|PABPC4(0)\|PPIE(-168.6kb)\|PPIEL(+10.56kb)\|SNORA55(+2.746kb) | 1 |
| rs787540 | 1 | 68051137 | A | G | -0.0142 | 0.0017 | 3.6E-16 | GADD45A | GADD45A(-99.72kb)\|GNG12(-116kb)\|IL12RB2(+188.6kb)\|SERBP1(+155kb) | 0 |
| rs6673575 | 1 | 81349395 | A | G | 0.0138 | 0.0017 | 2.7E-15 | LPHN2 | . | 0 |
| rs963912 | 1 | 92013998 | A | G | -0.0206 | 0.0032 | 8.2E-11 | CDC7 | CDC7(+22.68kb)\|HFM1(+143.6kb)\|TGFBR3(-131.9kb) | 0 |
| rs2391018 | 1 | 92034142 | A | G | -0.0119 | 0.002 | 4.5E-09 | CDC7 | CDC7(+42.82kb)\|HFM1(+163.7kb)\|TGFBR3(-111.8kb) | 0 |
| rs1192415 | 1 | 92077097 | A | G | -0.0682 | 0.002 | 2.8E-243 | HSP90B3P | CDC7(+85.78kb)\|TGFBR3(-68.8kb) | 0 |
| rs2765879 | 1 | 92135804 | A | G | -0.0097 | 0.0017 | 2.2E-08 | TGFBR3 | CDC7(+144.5kb)\|TGFBR3(-10.1kb) | 0 |
| rs2810886 | 1 | 92163330 | A | G | 0.0129 | 0.0019 | 5.6E-12 | TGFBR3 | CDC7(+172kb)\|TGFBR3(0) | 0 |
| rs61785513 | 1 | 110608419 | A | G | 0.0106 | 0.0019 | 1.1E-08 | ALX3 | AHCYL1(+42.06kb)\|ALX3(0)\|CSF1(+134.8kb)\|KCNC4(-144.9kb)\|KCNC4-AS1(-142.7kb)\|SLC6A17(-84.71kb)\|STRIP1(+11.16kb)\|UBL4B(-46.64kb) | 0 |
| rs12136690 | 1 | 116208944 | T | C | 0.0259 | 0.0019 | 3.8E-41 | VANGL1 | CASQ2(-33.68kb)\|NHLH2(-170.1kb)\|VANGL1(0) | 0 |
| rs11264319 | 1 | 155083942 | T | C | -0.0116 | 0.0016 | 1.1E-13 | EFNA1 | ADAM15(+48.69kb)\|CKS1B(+132.2kb)\|CLK2(-148.7kb)\|DCST1(+60.54kb)\|DCST2(+77.68kb)\|DPM3(-28.42kb)\|EFNA1(-16.41kb)\|EFNA3(+23.93kb)\|EFNA4(+41.91kb)\|FAM189B(-133.1kb)\|FDPS(-194.6kb)\|FLAD1(+118.4kb)\|GBA(-120.3kb)\|GBAP1(-99.67kb)\|HCN3(-163.3kb)\|KRTCAP2(-57.94kb)\|LENEP(+117.2kb)\|LOC100505666(+47.48kb)\|MIR92B(-81.02kb)\|MIR4258(+135.7kb)\|MTX1(- | 0 |

| | | | | | | | | | | |
|---|---|---|---|---|---|---|---|---|---|---|
| | | | | | | | | | 94.55kb)\|MUC1(-74.36kb)\|PBXIP1(+155.4kb)\|PKLR(-175.1kb)\|PMVK(+174.5kb)\|PYGO2(+149.7kb)\|SCAMP3(-141.8kb)\|SHC1(+137kb)\|SLC50A1(-23.88kb)\|THBS3(-81.44kb)\|TRIM46(-62.32kb)\|ZBTB7B(+92.94kb)\|=missense | |
| rs970741 | 1 | 169480045 | T | C | 0.0114 | 0.0017 | 3.8E-11 | F5 | BLZF1(+114.3kb)\|CCDC181(+83.38kb)\|F5(-1.146kb)\|NME7(+142.8kb)\|SELL(-179.8kb)\|SELP(-78.04kb)\|SLC19A2(+24.84kb)\|=missense | 0 |
| rs2239854 | 1 | 169525808 | A | G | 0.0186 | 0.0018 | 2.8E-26 | F5 | BLZF1(+160kb)\|CCDC181(+129.1kb)\|F5(0)\|NME7(+188.6kb)\|SELE(-166kb)\|SELL(-134kb)\|SELP(-32.28kb)\|SLC19A2(+70.6kb) | 0 |
| rs11584075 | 1 | 201598915 | A | C | 0.0288 | 0.0031 | 1.1E-20 | NAV1 | CSRP1(+122.5kb)\|IPO9(-199.4kb)\|IPO9-AS1(-58.47kb)\|MIR1231(-178.8kb)\|MIR5191(-89.72kb)\|NAV1(-18.53kb)\|PHLDA3(+160.6kb)\|RNU6-79P(-138kb)\|RPS10P7(+109.2kb) | 0 |
| rs114772174 | 1 | 227564515 | T | G | 0.0531 | 0.003 | 4.6E-71 | CDC42BPA | CDC42BPA(+58.69kb)\|ZNF678(-186.7kb) | 0 |
| rs7595075 | 2 | 264019 | A | C | -0.0109 | 0.0018 | 1.7E-09 | SH3YL1 | ACP1(-0.849kb)\|FAM150B(-15.54kb)\|SH3YL1(0) | 1 |
| rs13409347 | 2 | 5680264 | T | G | -0.0112 | 0.0016 | 5.3E-12 | SOX11 | LINC01248(-94.01kb)\|SOX11(-152.5kb) | 1 |
| rs629526 | 2 | 16475090 | T | C | 0.0112 | 0.0017 | 3.3E-11 | FAM49A | . | 0 |
| rs72778322 | 2 | 19378101 | A | G | -0.072 | 0.0069 | 2.8E-25 | MIR4757 | MIR4757(-170.1kb)\|OSR1(-173.1kb) | 0 |
| rs851357 | 2 | 19473075 | T | C | -0.0107 | 0.0016 | 8.6E-11 | MIR4757 | MIR4757(-75.11kb)\|OSR1(-78.17kb) | 0 |
| rs6726716 | 2 | 28381833 | A | G | -0.0117 | 0.0019 | 8.1E-10 | BRE | BRE(0)\|LOC100505716(-148.7kb)\|MIR4263(+162.5kb) | 1 |
| rs13417287 | 2 | 33219147 | T | C | -0.0163 | 0.002 | 7.4E-17 | LTBP1 | LINC00486(+56.88kb)\|LOC100271832(+47.94kb)\|LTBP1(0)\|TTC27(+173kb) | 0 |
| rs115781177 | 2 | 33348494 | A | G | 0.0203 | 0.0032 | 2.7E-10 | LTBP1 | LINC00486(+186.2kb)\|LOC100271832(+177.3kb)\|LTBP1(0) | 0 |
| rs163524 | 2 | 45157553 | A | C | -0.0158 | 0.0021 | 3.7E-14 | SIX3 | CAMKMT(+157.8kb)\|SIX2(-74.77kb)\|SIX3(-11.48kb)\|SIX3-AS1(-9.739kb) | 0 |
| rs11686318 | 2 | 56205824 | T | C | 0.0169 | 0.0019 | 3.1E-18 | MIR217 | EFEMP1(+54.53kb)\|MIR216A(-10.26kb)\|MIR216B(-22.02kb)\|MIR217(-4.277kb) | 0 |
| rs13009725 | 2 | 85998858 | T | C | -0.0126 | 0.0017 | 4.5E-14 | ATOH8 | ATOH8(0)\|C2orf68(+159.7kb)\|GNLY(+72.98kb)\|LOC284950(-43.39kb)\|LOC101928113(-117.5kb)\|MIR6071(-11.86kb)\|RNF181(+174kb)\|SFTPB(+103kb)\|ST3GAL5(-67.41kb)\|TMEM150A(+169kb)\|USP39(+122.5kb)\|VAMP5(+178.3kb)\|VAMP8(+189.7kb) | 0 |
| rs11684168 | 2 | 88694167 | T | C | 0.0114 | 0.002 | 2.1E-08 | FOXI3 | EIF2AK3(-162.1kb)\|FOXI3(-53.56kb)\|LOC101928371(-144.1kb)\|TEX37(-130kb) | 0 |
| rs2241843 | 2 | 111879381 | A | C | 0.0099 | 0.0016 | 5.2E-10 | BCL2L11 | ACOXL(+3.582kb)\|BCL2L11(0)\|MIR4435-1(-199.2kb)\|MIR4435-2(-199.2kb) | 1 |
| rs3748916 | 2 | 113984033 | A | G | -0.012 | 0.0015 | 6.2E-15 | PAX8 | IL1F10(+150.6kb)\|IL1RN(+92.44kb)\|IL36B(+173.6kb)\|IL36RN(+161.7kb)\|PAX8(0)\|PAX8-AS1(-9.07kb)\|PSD4(+23.36kb) | 0 |
| rs10439201 | 2 | 134437102 | T | C | 0.0119 | 0.0018 | 1.1E-11 | NCKAP5 | NCKAP5(+111.1kb) | 0 |

| | | | | | | | | | | |
|---|---|---|---|---|---|---|---|---|---|---|
| rs980772 | 2 | 145442190 | T | G | 0.0149 | 0.0016 | 5.8E-20 | DKFZp686O1327 | LOC101928455(+105.2kb)\|TEX41(0)\|ZEB2(+164.2kb)\|ZEB2-AS1(+163.7kb) | 0 |
| rs35152708 | 2 | 147152024 | A | G | 0.0094 | 0.0015 | 1.3E-09 | PABPC1P2 | PABPC1P2(-192.6kb) | 1 |
| rs2544373 | 2 | 170155249 | A | T | 0.011 | 0.0016 | 1.1E-11 | LRP2 | BBS5(-180.8kb)\|LRP2(0) | 0 |
| rs17428076 | 2 | 172851936 | C | G | 0.0103 | 0.0019 | 4.8E-08 | HAT1 | DLX1(-98.27kb)\|DLX2(-112.2kb)\|HAT1(+3.336kb)\|METAP1D(-12.87kb)\|SLC25A12(+101.1kb) | 1 |
| rs2675089 | 2 | 183914247 | A | G | -0.0171 | 0.0026 | 8.9E-11 | NCKAP1 | DUSP19(-29.04kb)\|FRZB(+182.7kb)\|NCKAP1(+10.66kb)\|NUP35(-67.97kb) | 1 |
| rs73049729 | 2 | 190949599 | A | C | 0.0171 | 0.0023 | 8.2E-14 | MSTN | C2orf88(-52.89kb)\|HIBCH(-119.8kb)\|MSTN(+22.14kb) | 1 |
| rs76952435 | 2 | 208647635 | C | G | 0.0215 | 0.0039 | 2.4E-08 | FZD5 | CCNYL1(+26.74kb)\|CREB1(+177.4kb)\|FZD5(+13.49kb)\|METTL21A(+157.7kb)\|MIR4775(+28.03kb)\|PLEKHM3(-38.38kb) | 1 |
| rs7602943 | 2 | 217632085 | A | G | 0.0156 | 0.0023 | 1.1E-11 | IGFBP5 | IGFBP2(+102.9kb)\|IGFBP5(+71.81kb)\|LINC01280(+160.4kb)\|TNP1(-92.1kb) | 1 |
| rs718686 | 2 | 218385952 | A | G | 0.0133 | 0.002 | 2.6E-11 | DIRC3 | DIRC3(0) | 0 |
| rs2565684 | 2 | 218477690 | C | G | -0.019 | 0.002 | 3.5E-21 | DIRC3 | DIRC3(0)\|TNS1(-186.8kb) | 0 |
| rs413028 | 2 | 225904282 | C | G | 0.0087 | 0.0015 | 1.6E-08 | DOCK10 | DOCK10(0)\|MIR4439(+29.02kb) | 1 |
| rs1550094 | 2 | 233385396 | A | G | 0.0166 | 0.0018 | 3.7E-21 | PRSS56 | ALPI(+60.65kb)\|ALPP(+137.8kb)\|ALPPL2(+110kb)\|CHRND(-5.473kb)\|CHRNG(-19.04kb)\|DIS3L2(+176.7kb)\|ECEL1(+32.83kb)\|ECEL1P2(+133.6kb)\|EFHD1(-85.37kb)\|EIF4E2(-29.9kb)\|GIGYF2(-176.6kb)\|MIR5001(-29.79kb)\|PRSS56(0)\|TIGD1(-27.38kb) | 0 |
| rs2948090 | 3 | 20070747 | A | C | 0.0106 | 0.0016 | 8.4E-11 | KAT2B | EFHB(+95.04kb)\|KAT2B(-10.78kb)\|PP2D1(+16.98kb)\|RAB5A(+44.08kb)\|SGOL1(-131.3kb)\|SGOL1-AS1(-145kb) | 0 |
| rs138526983 | 3 | 20152113 | T | C | 0.0411 | 0.0067 | 7.8E-10 | KAT2B | EFHB(+176.4kb)\|KAT2B(0)\|PP2D1(+98.35kb)\|RAB5A(+125.4kb)\|SGOL1(-49.97kb)\|SGOL1-AS1(-63.62kb) | 0 |
| rs9863531 | 3 | 21722110 | T | C | 0.0219 | 0.0033 | 2.6E-11 | ZNF385D | ZNF385D(0) | 0 |
| rs12487626 | 3 | 25043829 | C | G | -0.0172 | 0.0018 | 7E-22 | RARB | RARB(-172kb) | 0 |
| rs321537 | 3 | 25190819 | T | C | 0.0122 | 0.0021 | 1E-08 | RARB | RARB(-25kb) | 0 |
| rs6787363 | 3 | 25388569 | A | G | 0.0232 | 0.0016 | 1.6E-45 | RARB | RARB(0) | 0 |
| rs13070407 | 3 | 25493178 | T | C | 0.0107 | 0.0018 | 6.9E-09 | RARB | RARB(0)\|TOP2B(-146.2kb) | 0 |
| rs73048068 | 3 | 25595631 | A | G | -0.012 | 0.002 | 3.7E-09 | RARB | MIR4442(-110.7kb)\|NGLY1(-164.8kb)\|RARB(0)\|TOP2B(-43.76kb) | 0 |
| rs114103097 | 3 | 62360232 | A | G | -0.0159 | 0.0028 | 2E-08 | FEZF2 | C3orf14(+38.34kb)\|CADPS(-23.79kb)\|FEZF2(+1.042kb)\|PTPRG(+79.66kb)\|PTPRG-AS1(+55.61kb) | 0 |
| rs35667547 | 3 | 64547477 | C | G | -0.0165 | 0.0025 | 2.8E-11 | ADAMTS9 | ADAMTS9(0)\|ADAMTS9-AS2(-123.1kb)\|LOC101929335(0)\|MIR548A2(-158.2kb) | 1 |
| rs77877421 | 3 | 71182447 | A | T | -0.0362 | 0.0037 | 9.1E-23 | FOXP1 | FOXP1(0) | 0 |
| rs9284814 | 3 | 88273071 | A | G | 0.015 | 0.0026 | 4.9E-09 | C3orf38 | C3orf38(+65.96kb)\|CGGBP1(+74.06kb)\|ZNF654(+79.26kb)\|=missense | 1 |

| | | | | | | | | | | |
|---|---|---|---|---|---|---|---|---|---|---|
| rs2623343 | 3 | 99136009 | T | C | -0.0128 | 0.0017 | 6.6E-14 | MIR548G | MIR548G(-137.1kb) | 1 |
| rs2063638 | 3 | 99248498 | T | C | -0.0206 | 0.003 | 5.1E-12 | MIR548G | COL8A1(-108.9kb)\|MIR548G(-24.65kb) | 1 |
| rs1021103 | 3 | 99603230 | A | G | -0.0148 | 0.0019 | 1.2E-14 | FILIP1L | CMSS1(0)\|COL8A1(+87.65kb)\|FILIP1L(0)\|HP09053(+60.52kb)\|MIR548G(0)\|MIR3921(-79.93kb)\|=missense | 0 |
| rs17398137 | 3 | 100625703 | A | G | -0.0265 | 0.0021 | 4.2E-36 | ABI3BP | ABI3BP(0)\|TFG(+157.9kb) | 0 |
| rs2687728 | 3 | 127885008 | A | C | 0.0124 | 0.0021 | 3.2E-09 | EEFSEC | EEFSEC(0)\|KBTBD12(+178.5kb)\|RUVBL1(+42.34kb)\|SEC61A1(+94.48kb) | 1 |
| rs6774654 | 3 | 156770833 | A | G | 0.0106 | 0.0017 | 1.1E-09 | LEKR1 | CCNL1(-94.75kb)\|LEKR1(+6.915kb)\|LINC00880(-28.62kb)\|LINC00881(-36.84kb) | 0 |
| rs79932746 | 3 | 157460522 | C | G | 0.0129 | 0.0022 | 6.9E-09 | C3orf55 | C3orf55(+141.5kb) | 1 |
| rs12633493 | 3 | 171778354 | T | C | 0.0095 | 0.0017 | 1.4E-08 | FNDC3B | FNDC3B(0) | 1 |
| rs977265 | 3 | 189914180 | T | C | -0.011 | 0.0018 | 4.2E-10 | LEPREL1 | CLDN1(-109.3kb)\|CLDN16(-191.5kb)\|LEPREL1(+73.95kb) | 1 |
| rs4861118 | 4 | 41528563 | A | G | -0.0093 | 0.0016 | 1.1E-08 | LIMCH1 | LIMCH1(0) | 0 |
| rs62336823 | 4 | 53757661 | T | C | -0.0126 | 0.0019 | 1E-10 | SCFD2 | DANCR(+177.4kb)\|ERVMER34-1(+139.9kb)\|LOC152578(+76.03kb)\|MIR4449(+178.7kb)\|RASL11B(+24.66kb)\|SCFD2(0)\|SNORA26(+178.1kb) | 1 |
| rs11133309 | 4 | 55037174 | T | C | 0.0128 | 0.0019 | 3.1E-11 | PDGFRA | CHIC2(+106.4kb)\|GSX2(+69.05kb)\|PDGFRA(-58.09kb)\|RPL21P44(+183.7kb) | 0 |
| rs35471029 | 4 | 77690812 | C | G | -0.0179 | 0.0031 | 8.9E-09 | SHROOM3 | MIR4450(+196kb)\|SEPT11(-180.1kb)\|SHROOM3(0)\|SOWAHB(-125.3kb) | 1 |
| rs74764079 | 4 | 81952637 | A | T | -0.0441 | 0.0051 | 3.4E-18 | BMP3 | BMP3(0)\|C4orf22(+67.73kb)\|PRKG2(-55.89kb) | 0 |
| rs346504 | 4 | 86857864 | T | G | 0.011 | 0.0017 | 7.2E-11 | ARHGAP24 | ARHGAP24(0)\|LOC101929064(-183.1kb)\|MAPK10(-78.41kb) | 1 |
| rs7679158 | 4 | 111812327 | A | G | -0.0093 | 0.0017 | 2.2E-08 | PITX2 | . | 1 |
| rs1566795 | 4 | 112375172 | T | C | 0.0136 | 0.0017 | 3.1E-16 | C4orf32 | . | 0 |
| rs67399037 | 4 | 126397056 | T | C | -0.0094 | 0.0016 | 9.2E-09 | FAT4 | FAT4(0)\|MIR2054(-31.36kb) | 1 |
| rs13167406 | 5 | 169301 | A | G | 0.0102 | 0.0016 | 1.3E-10 | PLEKHG4B | AHRR(-135kb)\|CCDC127(-35.57kb)\|LOC102467073(-100.4kb)\|LRRC14B(-22.32kb)\|PDCD6(-102.4kb)\|PLEKHG4B(0)\|SDHA(-49.05kb) | 1 |
| rs72759609 | 5 | 31952051 | T | C | 0.0332 | 0.0028 | 4.8E-33 | PDZD2 | GOLPH3(-172.8kb)\|MIR4279(+15.79kb)\|PDZD2(0) | 0 |
| rs251526 | 5 | 52586585 | A | G | -0.0174 | 0.003 | 5.5E-09 | LOC257396 | FST(-189.7kb)\|ITGA2(+196kb)\|LOC257396(+175.6kb)\|MOCS2(+181kb) | 0 |
| rs30371 | 5 | 55742832 | T | C | -0.0124 | 0.0017 | 9.9E-13 | ANKRD55 | LOC101928448(-64.39kb)\|LOC102467147(-10.79kb) | 0 |
| rs186818 | 5 | 78205075 | A | G | 0.0106 | 0.0018 | 3.3E-09 | ARSB | ARSB(0)\|BHMT2(-160.5kb)\|DMGDH(-88.31kb) | 1 |
| rs79051849 | 5 | 81848970 | A | G | -0.0123 | 0.0021 | 3E-09 | ATP6AP1L | . | 1 |
| rs10474101 | 5 | 82739803 | T | C | 0.0136 | 0.0016 | 3.2E-17 | VCAN | HAPLN1(-194.2kb)\|VCAN(-27.69kb)\|XRCC4(+90.22kb) | 0 |

| | | | | | | | | | | |
|---|---|---|---|---|---|---|---|---|---|---|
| rs115803211 | 5 | 87907437 | T | C | -0.0349 | 0.0031 | 3.3E-30 | LOC645323 | LINC00461(0)\|LOC102546226(+172.5kb)\|MEF2C(-106.6kb)\|MIR9-2(-55.23kb)\|TMEM161B-AS1(+174.9kb) | 0 |
| rs10463643 | 5 | 112032675 | T | C | 0.0107 | 0.0017 | 6E-10 | APC | APC(-10.53kb)\|LOC102467214(+66.06kb)\|LOC102467216(+14.09kb)\|REEP5(-179.4kb)\|SRP19(-164.2kb) | 1 |
| rs6892256 | 5 | 126546211 | T | G | 0.0129 | 0.0016 | 1.8E-15 | MEGF10 | C5orf63(+137kb)\|MARCH3(+179.7kb)\|MEGF10(-80.24kb) | 1 |
| rs6888519 | 5 | 172171643 | T | C | 0.0195 | 0.0035 | 3.4E-08 | DUSP1 | DUSP1(-23.45kb)\|ERGIC1(-89.58kb)\|LOC101928093(-10.86kb)\|NEURL1B(+53.11kb) | 1 |
| rs1962432 | 6 | 1680806 | T | C | -0.0099 | 0.0017 | 9.1E-09 | GMDS | FOXC1(+66.68kb)\|GMDS(0) | 0 |
| rs4512269 | 6 | 1987897 | A | T | 0.0177 | 0.0024 | 1.9E-13 | GMDS | GMDS(0) | 0 |
| rs78271362 | 6 | 11425571 | A | C | 0.0161 | 0.0026 | 5.6E-10 | NEDD9 | NEDD9(+42.99kb)\|TMEM170B(-112.9kb) | 1 |
| rs2092524 | 6 | 39529692 | A | G | 0.0157 | 0.0017 | 5.5E-20 | KIF6 | KIF6(0) | 0 |
| rs943080 | 6 | 43826627 | T | C | -0.0128 | 0.0016 | 7.6E-15 | LOC100132354 | C6orf223(-141.7kb)\|LOC100132354(-32.14kb)\|MRPS18A(+171.1kb)\|RSPH9(+187.9kb)\|VEGFA(+72.4kb) | 1 |
| rs41402445 | 6 | 50086239 | A | G | -0.0112 | 0.002 | 1.7E-08 | DEFB112 | DEFB110(+96.54kb)\|DEFB112(+69.88kb)\|DEFB113(+148.9kb)\|DEFB114(+154.4kb)\|DEFB133(+169.1kb) | 1 |
| rs7744813 | 6 | 73643289 | A | C | -0.0143 | 0.0017 | 1.2E-17 | KCNQ5 | KCNQ5(0)\|MIR4282(-34.12kb) | 0 |
| rs313160 | 6 | 86042898 | A | G | -0.013 | 0.0023 | 7.1E-09 | NT5E | NT5E(-116.4kb)\|SNX14(-172.3kb) | 1 |
| rs59472001 | 6 | 122668771 | T | C | -0.0152 | 0.0018 | 3E-17 | HSF2 | HSF2(-51.92kb)\|PKIB(-124.3kb)\|SERINC1(-95.72kb) | 0 |
| rs9388489 | 6 | 126698719 | A | G | -0.0176 | 0.0016 | 4.5E-27 | CENPW | CENPW(0) | 0 |
| rs17069768 | 6 | 140400161 | T | G | 0.0109 | 0.0019 | 1.8E-08 | MIR3668 | MIR3668(-126.2kb) | 1 |
| rs1335307 | 6 | 141646340 | A | T | -0.0092 | 0.0017 | 3.7E-08 | MIR4465 | . | 1 |
| rs117117522 | 7 | 47353446 | A | G | 0.0351 | 0.0062 | 1.2E-08 | TNS3 | TNS3(0) | 1 |
| rs78808004 | 7 | 73492035 | A | G | -0.0152 | 0.0027 | 3.6E-08 | LIMK1 | EIF4H(-96.67kb)\|ELN(+7.799kb)\|LAT2(-132.1kb)\|LIMK1(-6.071kb)\|MIR590(-113.5kb)\|RFC2(-153.8kb) | 1 |
| rs10281028 | 7 | 90232723 | A | G | -0.0116 | 0.0021 | 4.3E-08 | CDK14 | CDK14(0)\|CLDN12(+187.5kb) | 1 |
| rs34298819 | 7 | 115580893 | A | G | -0.0493 | 0.0072 | 5.7E-12 | TFEC | TFEC(0) | 1 |
| rs2402019 | 7 | 115598764 | T | G | -0.0141 | 0.0025 | 1.6E-08 | TFEC | TFEC(0) | 1 |
| rs1905014 | 8 | 13407297 | T | C | -0.0134 | 0.0016 | 1.6E-16 | C8orf48 | C8orf48(-17.05kb)\|DLC1(+34.87kb) | 0 |
| rs868444 | 8 | 49502481 | A | C | -0.0132 | 0.0023 | 4.6E-09 | EFCAB1 | EFCAB1(-125kb)\|LOC101929217(-30.48kb)\|LOC101929268(0) | 1 |
| rs72652582 | 8 | 61860746 | T | C | 0.0135 | 0.0021 | 7.9E-11 | LOC100130298 | CHD7(+80.16kb)\|LOC100130298(-17.93kb) | 0 |
| rs3857971 | 8 | 61969579 | A | G | -0.016 | 0.0017 | 4.8E-20 | LOC100130298 | CHD7(+189kb)\|LOC100130298(+89.27kb) | 0 |
| rs13258368 | 8 | 71548751 | A | G | 0.0118 | 0.0016 | 7.6E-13 | LOC286190 | LACTB2(-0.749kb)\|LOC286190(0)\|LOC101926892(+150.8kb)\|TRAM1(+28.06kb)\|XKR9(-32.85kb) | 1 |

| | | | | | | | | | | |
|---|---|---|---|---|---|---|---|---|---|---|
| rs12543430 | 8 | 72278010 | T | C | -0.0102 | 0.0016 | 1E-10 | EYA1 | EYA1(+3.543kb) | 1 |
| rs17734073 | 8 | 72598450 | A | G | -0.0246 | 0.0025 | 3.5E-23 | MSC | LOC100132891(-156.9kb)\|MSC(-155.3kb) | 0 |
| rs4276683 | 8 | 78939763 | T | G | 0.0152 | 0.0017 | 1.7E-18 | PKIA | . | 0 |
| rs12547416 | 8 | 88761223 | T | C | -0.0149 | 0.0016 | 1.3E-21 | DCAF4L2 | DCAF4L2(-121.7kb) | 0 |
| rs10956000 | 8 | 122146899 | A | G | 0.0129 | 0.0021 | 2E-09 | SNTB1 | . | 1 |
| rs6982521 | 8 | 122738226 | T | C | -0.011 | 0.0018 | 2.3E-09 | HAS2-AS1 | HAS2(+84.6kb)\|HAS2-AS1(+80.66kb) | 1 |
| rs2717600 | 8 | 143813988 | A | G | 0.0125 | 0.0016 | 3.8E-14 | C8orf55 | ARC(+118.2kb)\|BAI1(+187.6kb)\|CYP11B1(-139.8kb)\|CYP11B2(-178kb)\|GML(-102.2kb)\|JRK(+62.58kb)\|LOC100288181(+5.597kb)\|LY6D(-52.31kb)\|LY6K(+28.4kb)\|LYNX1(-31.77kb)\|LYPD2(-17.64kb)\|PSCA(+49.84kb)\|SLURP1(-8.373kb)\|THEM6(0) | 1 |
| rs10961437 | 9 | 14231361 | A | G | -0.0181 | 0.0023 | 4.4E-15 | NFIB | NFIB(0) | 1 |
| rs275528 | 9 | 76175652 | T | C | -0.0131 | 0.002 | 7.8E-11 | ANXA1 | MIR6130(-192.4kb) | 1 |
| rs11143754 | 9 | 76622068 | A | C | -0.0101 | 0.0016 | 4.5E-10 | RORB | MIR6130(0) | 1 |
| rs10512176 | 9 | 89252706 | T | C | -0.0178 | 0.0018 | 3.1E-23 | ZCCHC6 | . | 0 |
| rs35071563 | 9 | 89256298 | A | G | -0.0389 | 0.007 | 2.3E-08 | ZCCHC6 | . | 0 |
| rs1111066 | 9 | 89380805 | C | G | 0.0095 | 0.0016 | 2.8E-09 | GAS1 | GAS1(-178.5kb)\|LOC100506834(-182.8kb) | 1 |
| rs28457693 | 9 | 98217348 | A | G | -0.0211 | 0.0026 | 1.6E-15 | PTCH1 | FANCC(+137.4kb)\|LOC100507346(-8.542kb)\|PTCH1(0) | 0 |
| rs700981 | 9 | 98496244 | T | C | -0.0107 | 0.0017 | 2E-10 | C9orf130 | ERCC6L2(-141.7kb)\|LINC00476(-72.12kb) | 1 |
| rs4978765 | 9 | 111735885 | T | G | 0.02 | 0.003 | 3.4E-11 | CTNNAL1 | ACTL7A(+109.8kb)\|ACTL7B(+117.6kb)\|CTNNAL1(0)\|EPB41L4B(-198.4kb)\|FAM206A(+32.65kb)\|FRRS1L(-163.7kb)\|IKBKAP(+39.28kb)\|MIR32(-72.62kb)\|TMEM245(-41.53kb)\|=missense | 1 |
| rs35704524 | 9 | 134560240 | A | G | 0.0159 | 0.0019 | 4.2E-17 | RAPGEF1 | MED27(-175.3kb)\|POMT1(+161kb)\|PRRC2B(+184.7kb)\|RAPGEF1(0)\|SNORD62A(+194.3kb)\|SNORD62B(+194.3kb)\|UCK1(+153.6kb) | 0 |
| rs4570483 | 10 | 25054210 | A | G | 0.0135 | 0.0017 | 4.6E-15 | ARHGAP21 | ARHGAP21(+41.61kb)\|PRTFDC1(-83.32kb) | 0 |
| rs149176725 | 10 | 68991138 | C | G | 0.0358 | 0.0043 | 1.6E-16 | CTNNA3 | CTNNA3(0)\|LRRTM3(+129.8kb) | 0 |
| rs35898435 | 10 | 69962881 | A | C | 0.0272 | 0.0048 | 1.4E-08 | MYPN | ATOH7(-27.47kb)\|HERC4(+127.8kb)\|HNRNPH3(-128.9kb)\|MYPN(0)\|PBLD(-79.54kb)\|RUFY2(-138kb) | 0 |
| rs7916697 | 10 | 69991853 | A | G | -0.0797 | 0.0019 | 0 | ATOH7 | ATOH7(0)\|DNA2(-182kb)\|HERC4(+156.8kb)\|HNRNPH3(-99.91kb)\|MYPN(+20.08kb)\|PBLD(-50.56kb)\|RUFY2(-109kb) | 0 |
| rs116979034 | 10 | 70288480 | A | G | 0.0245 | 0.0044 | 3.3E-08 | SLC25A16 | CCAR1(-192.4kb)\|DNA2(+56.6kb)\|HNRNPH3(+185.5kb)\|PBLD(+195.8kb)\|RUFY2(+121.4kb)\|SLC25A16(+1.2kb)\|TET1(-31.64kb) | 0 |

| | | | | | | | | | | |
|---|---|---|---|---|---|---|---|---|---|---|
| rs113492938 | 10 | 70410975 | A | C | 0.0357 | 0.006 | 3E-09 | TET1 | CCAR1(-69.92kb)\|DNA2(+179.1kb)\|SLC25A16(+123.7kb)\|SNORD98(-104kb)\|STOX1(-176.3kb)\|TET1(0) | 0 |
| rs76741022 | 10 | 70540624 | A | G | -0.0255 | 0.0038 | 1.4E-11 | CCAR1 | CCAR1(0)\|DDX21(-175.3kb)\|DDX50(-120.4kb)\|SNORD98(+25.63kb)\|STOX1(-46.67kb)\|TET1(+86.38kb) | 0 |
| rs12357925 | 10 | 71609313 | C | G | 0.0143 | 0.0021 | 3.7E-12 | COL13A1 | COL13A1(0) | 0 |
| rs117927780 | 10 | 72432306 | T | G | -0.0105 | 0.0017 | 7.3E-10 | ADAMTS14 | ADAMTS14(-0.252kb)\|PALD1(+104.1kb)\|PRF1(+69.78kb)\|SGPL1(-143.4kb)\|TBATA(-98.69kb) | 0 |
| rs3740440 | 10 | 72521309 | C | G | -0.0097 | 0.0017 | 4.8E-09 | ADAMTS14 | ADAMTS14(0)\|PALD1(+193.1kb)\|PCBD1(-122kb)\|PRF1(+158.8kb)\|SGPL1(-54.39kb)\|TBATA(-9.685kb) | 0 |
| rs12260286 | 10 | 77844316 | T | C | -0.0113 | 0.0017 | 1.1E-11 | C10orf11 | C10orf11(0) | 1 |
| rs11594050 | 10 | 90093298 | T | C | 0.0128 | 0.0018 | 3.6E-13 | RNLS | RNLS(0) | 1 |
| rs2149108 | 10 | 94866290 | T | C | -0.0118 | 0.0018 | 5.5E-11 | CYP26A1 | CYP26A1(+28.65kb)\|CYP26C1(+37.84kb)\|EXOC6(+47.04kb)\|MYOF(-199.9kb) | 0 |
| rs12411442 | 10 | 104349531 | A | G | 0.0167 | 0.0016 | 3.3E-24 | SUFU | ACTR1A(+87.02kb)\|ARL3(-83.95kb)\|C10orf95(+138.2kb)\|CUEDC2(+157.1kb)\|FBXL15(+166.6kb)\|MIR146B(+153.2kb)\|NFKB2(+187.2kb)\|PSD(+169.8kb)\|RPARP-AS1(+133.5kb)\|SFXN2(-124.8kb)\|SUFU(0)\|TMEM180(+112.7kb)\|TRIM8(-54.72kb)\|WBP1L(-154.2kb) | 0 |
| rs593864 | 11 | 30047720 | T | C | -0.0114 | 0.0019 | 2E-09 | KCNA4 | KCNA4(+9.143kb) | 1 |
| rs2146569 | 11 | 31609108 | A | C | -0.023 | 0.0019 | 5.4E-35 | ELP4 | DNAJC24(+154.7kb)\|ELP4(0)\|IMMP1L(+77.94kb)\|PAX6(-197.2kb) | 0 |
| rs147470123 | 11 | 31872347 | A | G | -0.0321 | 0.0043 | 8.8E-14 | DKFZp686K1684 | DKFZp686K1684(0)\|ELP4(+66.27kb)\|PAUPAR(+21.5kb)\|PAX6(+32.84kb) | 0 |
| rs1126809 | 11 | 89017961 | A | G | -0.0109 | 0.0017 | 4.3E-10 | TYR | NOX4(-39.56kb)\|TYR(0)\|=MISSENSE | 0 |
| rs7101609 | 11 | 92623493 | C | G | -0.0106 | 0.0016 | 8.9E-11 | FAT3 | FAT3(0)\|MTNR1B(-79.3kb) | 1 |
| rs7949779 | 11 | 95282879 | C | G | -0.0106 | 0.0019 | 1.5E-08 | FAM76B | . | 1 |
| rs73048927 | 12 | 3375780 | A | G | 0.0273 | 0.0032 | 3.8E-17 | TSPAN9 | PRMT8(-114.7kb)\|TSPAN9(0) | 0 |
| rs55637942 | 12 | 31038208 | T | C | 0.0208 | 0.0022 | 3.7E-21 | TSPAN11 | CAPRIN2(+130.8kb)\|DDX11(-188.6kb)\|DDX11-AS1(-135.5kb)\|IPO8(+189.3kb)\|LINC00941(+82.56kb)\|TSPAN11(-41.63kb) | 0 |
| rs11182092 | 12 | 43864034 | T | C | -0.0181 | 0.0033 | 3.8E-08 | ADAMTS20 | ADAMTS20(0) | 1 |
| rs1994670 | 12 | 77997178 | A | G | -0.0127 | 0.0017 | 2.4E-13 | NAV3 | . | 0 |
| rs11609693 | 12 | 78257054 | T | C | -0.0105 | 0.0018 | 7.6E-09 | NAV3 | NAV3(0) | 0 |
| rs1511581 | 12 | 84007857 | T | C | 0.0215 | 0.0016 | 3.9E-40 | TMTC2 | . | 0 |
| rs2642536 | 12 | 85473240 | A | G | 0.01 | 0.0016 | 6.4E-10 | LRRIQ1 | LRRIQ1(0)\|SLC6A15(+166.6kb)\|TSPAN19(+43.18kb) | 0 |
| rs4964618 | 12 | 108155108 | A | G | 0.0096 | 0.0017 | 2.6E-08 | PRDM4 | ASCL4(-13.05kb)\|BTBD11(+101.7kb)\|LOC728739(-141.8kb)\|LOC101929162(+1.363kb)\|PRDM4(+0.194kb)\|PWP1(+48.85kb) | 1 |

| | | | | | | | | | | |
|---|---|---|---|---|---|---|---|---|---|---|
| rs73191214 | 12 | 108975623 | A | G | 0.0155 | 0.0021 | 5.5E-14 | TMEM119 | CORO1C(-63.26kb)\|FICD(+62.24kb)\|ISCU(+12.46kb)\|LOC102723562(+108.2kb)\|SART3(+20.46kb)\|SELPLG(-40.06kb)\|TMEM119(-7.998kb) | 0 |
| rs12865567 | 13 | 24869978 | A | C | -0.0145 | 0.0026 | 3.5E-08 | SPATA13 | C1QTNF9(-13.74kb)\|MIR2276(+133.3kb)\|PARP4(-125.1kb)\|SPATA13(0)\|SPATA13-AS1(+41.4kb) | 1 |
| rs111516934 | 13 | 26624935 | T | C | 0.0117 | 0.002 | 1.1E-08 | SHISA2 | ATP8A2(+29.52kb)\|RNF6(-162kb)\|SHISA2(0) | 1 |
| rs7330329 | 13 | 36644724 | T | G | -0.0119 | 0.002 | 1.5E-09 | DCLK1 | CCDC169(-156.5kb)\|CCDC169-SOHLH2(-97.62kb)\|DCLK1(0)\|MIR548F5(+129.3kb)\|SOHLH2(-97.62kb) | 1 |
| rs9531993 | 13 | 37475197 | C | G | -0.012 | 0.0019 | 6.1E-10 | SMAD9 | ALG5(-48.71kb)\|EXOSC8(-99.48kb)\|RFXAP(+71.46kb)\|SMAD9(0)\|SUPT20H(-108.3kb) | 1 |
| rs9534439 | 13 | 47192049 | T | C | 0.0196 | 0.002 | 5.7E-23 | LRCH1 | ESD(-153.3kb)\|LOC101929344(+151.1kb)\|LRCH1(0) | 0 |
| rs2806933 | 13 | 53643370 | A | C | -0.0135 | 0.0017 | 1.9E-15 | OLFM4 | OLFM4(+17.17kb) | 1 |
| rs3742244 | 13 | 101256563 | A | T | 0.011 | 0.0019 | 1.2E-08 | TMTC4 | GGACT(+15.52kb)\|NALCN-AS1(-104kb)\|PCCA(+73.87kb)\|PCCA-AS1(+123.3kb)\|TMTC4(0) | 1 |
| rs56346803 | 13 | 111011400 | A | T | -0.0127 | 0.0021 | 1.1E-09 | COL4A2 | COL4A1(+51.9kb)\|COL4A2(0)\|COL4A2-AS1(-143.5kb)\|MIR8073(+18.02kb)\|RAB20(-164kb) | 1 |
| rs10149874 | 14 | 52004575 | A | G | -0.0102 | 0.0017 | 4.3E-09 | FRMD6 | FRMD6(0)\|FRMD6-AS1(-111.7kb)\|FRMD6-AS2(0)\|LINC00640(+172.3kb) | 1 |
| rs34321730 | 14 | 54667777 | A | T | 0.0148 | 0.0021 | 1.5E-12 | CDKN3 | CDKN3(-195.9kb) | 0 |
| rs113374826 | 14 | 59156412 | T | C | -0.0395 | 0.0072 | 4.7E-08 | DACT1 | DACT1(+41.37kb)\|KIAA0586(+140.9kb)\|LOC102723742(-138.6kb) | 1 |
| rs2103941 | 14 | 59480893 | A | G | -0.0179 | 0.0023 | 3.4E-15 | DAAM1 | DAAM1(-174.5kb)\|LOC102723742(0) | 0 |
| rs61985972 | 14 | 59550263 | A | G | 0.03 | 0.0033 | 2.8E-19 | DAAM1 | DAAM1(-105.1kb)\|LOC102723742(+66.22kb) | 0 |
| rs77587000 | 14 | 60624161 | T | C | -0.1056 | 0.0074 | 1.6E-46 | DHRS7 | DHRS7(0)\|LRRC9(+93.88kb)\|PCNXL4(+22.63kb)\|PPM1A(-88.31kb) | 0 |
| rs33912345 | 14 | 60976537 | A | C | 0.0203 | 0.0017 | 5E-34 | SIX6 | C14orf39(+23.77kb)\|SIX1(-134.9kb)\|SIX4(-199.7kb)\|SIX6(0) | 0 |
| rs10483740 | 14 | 61988231 | T | C | 0.0436 | 0.0075 | 5.6E-09 | PRKCH | FLJ22447(-49.03kb)\|HIF1A(-173.9kb)\|HIF1A-AS1(-159.5kb)\|LOC101927780(-34.76kb)\|PRKCH(0) | 1 |
| rs76320564 | 14 | 65115953 | T | C | -0.0225 | 0.0021 | 2.1E-26 | PLEKHG3 | AKAP5(+174.7kb)\|HSPA2(+106kb)\|LOC102723809(+108.9kb)\|MIR548AZ(+188.5kb)\|MIR7855(-136.4kb)\|MTHFD1(+189.2kb)\|PLEKHG3(-55.24kb)\|PPP1R36(+59.86kb)\|SPTB(-97.05kb)\|ZBTB1(+115.5kb)\|ZBTB25(+145.4kb) | 0 |
| rs34235559 | 14 | 85729874 | A | G | -0.0188 | 0.0017 | 7.5E-29 | FLRT2 | LINC00911(-130.3kb) | 0 |
| rs1289426 | 14 | 85922578 | A | G | -0.0144 | 0.002 | 2.5E-13 | FLRT2 | FLRT2(-73.91kb)\|LINC00911(+36.16kb) | 0 |
| rs2430356 | 14 | 92356289 | A | T | -0.0122 | 0.0019 | 3.1E-10 | FBLN5 | ATXN3(-168.6kb)\|CATSPERB(+157.9kb)\|FBLN5(0)\|TC2N(+22.41kb)\|TRIP11(-77.95kb) | 0 |

| | | | | | | | | | | |
|---|---|---|---|---|---|---|---|---|---|---|
| rs62007715 | 14 | 103917392 | T | C | 0.0134 | 0.002 | 1.9E-11 | MARK3 | APOPT1(-111.9kb)\|BAG5(-105.5kb)\|CKB(-68.6kb)\|EIF5(+106kb)\|KLC1(-178.1kb)\|MARK3(0)\|SNORA28(+113.1kb)\|TRMT61A(-78.12kb) | 0 |
| rs11635984 | 15 | 33012232 | T | C | -0.0089 | 0.0016 | 8.9E-09 | GREM1 | ARHGAP11A(+80.08kb)\|FMN1(-45.51kb)\|GOLGA8N(+112.7kb)\|GREM1(0)\|LOC100131315(+1.166kb)\|LOC100996255(+139.4kb)\|SCG5(+22.93kb)\|WHAMMP1(+186.3kb) | 1 |
| rs11073064 | 15 | 35028126 | A | T | -0.0107 | 0.0018 | 7.7E-09 | GJD2 | ACTC1(-52.17kb)\|AQR(-120.4kb)\|GJD2(-16.52kb)\|GOLGA8B(+152.4kb)\|LOC101928174(-19.16kb) | 1 |
| rs1077918 | 15 | 41203736 | A | G | -0.0128 | 0.002 | 5.7E-11 | VPS18 | C15orf62(+139.1kb)\|CHAC1(-41.9kb)\|DLL4(-17.79kb)\|DNAJC17(+104.1kb)\|GCHFR(+143.8kb)\|INO80(-67.34kb)\|PPP1R14D(+82.83kb)\|RAD51(+179.4kb)\|RHOV(+37.25kb)\|RMDN3(+156.3kb)\|SPINT1(+53.88kb)\|VPS18(+7.563kb)\|ZFYVE19(+96.97kb) | 1 |
| rs4776133 | 15 | 53610699 | A | T | -0.0097 | 0.0016 | 3.7E-09 | WDR72 | WDR72(-195.2kb) | 1 |
| rs7183049 | 15 | 54001509 | T | C | -0.0144 | 0.0023 | 6.1E-10 | WDR72 | WDR72(0) | 1 |
| rs1002007 | 15 | 71582050 | A | G | 0.012 | 0.0019 | 3.1E-10 | THSD4 | CT62(+174.2kb)\|LOC101929196(+100.7kb)\|THSD4(0) | 1 |
| rs1052622 | 15 | 74467856 | A | G | 0.012 | 0.0017 | 5E-12 | ISLR | CCDC33(-60.77kb)\|CYP11A1(-162.2kb)\|GOLGA6A(+92.96kb)\|ISLR(0)\|ISLR2(+38.71kb)\|LOC283731(+46.24kb)\|LOC729739(-186kb)\|PML(+127.7kb)\|STOML1(+180.9kb)\|STRA6(-3.951kb) | 0 |
| rs8029039 | 15 | 79100871 | C | G | -0.0109 | 0.0019 | 9.1E-09 | ADAMTS7 | ADAMTS7(0)\|CHRNA3(+187.2kb)\|CHRNB4(+167.3kb)\|CTSH(-113.2kb)\|LOC646938(+55.14kb)\|MORF4L1(-64.25kb)\|RASGRF1(-151.4kb) | 1 |
| rs7169595 | 15 | 84405574 | A | T | -0.0113 | 0.0016 | 5.7E-12 | ADAMTSL3 | ADAMTSL3(0)\|SH3GL3(+118.1kb) | 0 |
| rs980169 | 15 | 96720568 | A | G | -0.0155 | 0.0018 | 3.7E-18 | NR2F2 | MIR1469(-155.9kb)\|NR2F2(-148.6kb)\|NR2F2-AS1(-89.05kb) | 0 |
| rs4965806 | 15 | 101752435 | A | G | -0.0103 | 0.0016 | 2.4E-10 | CHSY1 | CHSY1(0)\|LOC100507472(-95.02kb)\|LRRK1(+142.1kb)\|PCSK6(-91.7kb)\|SNRPA1(-69.28kb)\|VIMP(-58.68kb) | 1 |
| rs2541593 | 16 | 103423 | A | C | -0.0113 | 0.002 | 1.5E-08 | POLR3K | DDX11L10(+39.33kb)\|HBA1(-123.3kb)\|HBA2(-119.4kb)\|HBM(-112.5kb)\|HBQ1(-126.9kb)\|HBZ(-99.43kb)\|ITFG3(-181.1kb)\|LUC7L(-135.5kb)\|MIR6859-1(+36.3kb)\|MIR6859-2(+36.3kb)\|MPG(-23.59kb)\|NPRL3(-32.38kb)\|POLR3K(0)\|RHBDF1(-4.634kb)\|SNRNP25(-0.405kb) | 1 |
| rs7184522 | 16 | 7460699 | A | G | 0.0142 | 0.0018 | 4.6E-16 | RBFOX1 | RBFOX1(0) | 0 |
| rs8059770 | 16 | 51394394 | A | G | 0.0104 | 0.0018 | 3.3E-09 | SALL1 | . | 0 |
| rs8053277 | 16 | 51469726 | T | C | 0.024 | 0.0018 | 5.2E-41 | SALL1 | . | 0 |
| rs28514893 | 16 | 51573119 | T | C | 0.0211 | 0.002 | 3.2E-26 | SALL1 | . | 0 |
| rs11076224 | 16 | 58239644 | A | C | 0.0185 | 0.0032 | 8E-09 | CSNK2A2 | C16orf80(+76.35kb)\|CCDC113(-44.2kb)\|CSNK2A2(+7.862kb)\|GINS3(-186.7kb)\|MMP15(+158.8kb)\|PRSS54(-74.26kb)\|USB1(+184.1kb) | 1 |

| | | | | | | | | | | |
|---|---|---|---|---|---|---|---|---|---|---|
| rs251798 | 16 | 69328442 | T | C | 0.01 | 0.0017 | 5.1E-09 | SNTB2 | CHTF8(+161.9kb)\|CIRH1A(+125.5kb)\|COG8(-34.08kb)\|CYB5B(-130.1kb)\|HAS3(+175.8kb)\|NIP7(-44.97kb)\|PDF(-34.08kb)\|SNTB2(0)\|TERF2(-61.02kb)\|TMED6(-48.71kb)\|VPS4A(-16.84kb) | 1 |
| rs10500557 | 16 | 71509779 | T | C | -0.0268 | 0.0047 | 1.3E-08 | ZNF19 | CALB2(+85.44kb)\|CHST4(-50.24kb)\|CMTR2(+186.3kb)\|LOC100132529(-89.14kb)\|MARVELD3(-150.3kb)\|PHLPP2(-169kb)\|TAT(-90.97kb)\|ZNF19(0)\|ZNF23(+13.66kb)\|=MISSENSE | 1 |
| rs8066620 | 17 | 738853 | T | C | -0.0098 | 0.0018 | 2.9E-08 | NXN | ABR(-167.9kb)\|DBIL5P(+80.28kb)\|FAM57A(+92.78kb)\|GEMIN4(+83.35kb)\|GLOD4(+53.28kb)\|MIR3183(-186.9kb)\|NXN(0)\|RNMTL1(+43.11kb)\|TIMM22(-161.5kb)\|VPS53(+120.8kb) | 1 |
| rs11655511 | 17 | 59287269 | T | C | -0.0297 | 0.0019 | 1E-52 | BCAS3 | BCAS3(0)\|TBX2(-190kb) | 0 |
| rs72843145 | 17 | 61052949 | T | C | 0.0153 | 0.0021 | 8.2E-13 | MIR548W | MARCH10(+167.2kb)\|MIR548W(0)\|TANC2(-33.95kb) | 0 |
| rs12453895 | 17 | 67578709 | T | C | 0.0095 | 0.0016 | 1.5E-09 | MAP2K6 | LOC101928122(-11.42kb)\|MAP2K6(+40.24kb) | 0 |
| rs2529681 | 17 | 68264797 | T | G | -0.011 | 0.0017 | 2.8E-10 | KCNJ2 | KCNJ2(+88.61kb)\|KCNJ2-AS1(+99.25kb)\|KCNJ16(+133.1kb) | 1 |
| rs717419 | 17 | 68718188 | T | C | 0.01 | 0.0016 | 3.3E-10 | KCNJ2 | . | 1 |
| rs12603193 | 17 | 70387448 | A | G | 0.0109 | 0.0016 | 1.2E-11 | LOC100499467 | LINC00673(-12.01kb) | 1 |
| rs62075722 | 17 | 79611271 | A | G | -0.0173 | 0.0017 | 9.2E-25 | TSPAN10 | ACTG1(+131.4kb)\|ARL16(-36.95kb)\|BAHCC1(+177.9kb)\|C17orf70(+91.84kb)\|CCDC137(-22.49kb)\|FAM195B(-169kb)\|FSCN2(+107.1kb)\|GCGR(-150.7kb)\|HGS(-39.69kb)\|MIR3186(+193.1kb)\|MIR6786(-49.52kb)\|MRPL12(-59.13kb)\|NPLOC4(+7.133kb)\|OXLD1(-20.79kb)\|P4HB(-189.8kb)\|PDE6G(-6.217kb)\|PPP1R27(-180.1kb)\|SLC25A10(-67.99kb)\|TSPAN10(0) | 0 |
| rs8269 | 17 | 81052423 | A | G | -0.0118 | 0.0018 | 4.6E-11 | METRNL | B3GNTL1(+42.74kb)\|FLJ43681(-122.2kb)\|METRNL(0)\|TBCD(+151.4kb) | 1 |
| rs17452020 | 18 | 359884 | A | G | 0.0111 | 0.002 | 1.2E-08 | COLEC12 | COLEC12(0)\|THOC1(+91.82kb)\|USP14(+146.1kb) | 1 |
| rs2621182 | 18 | 437286 | A | G | -0.0103 | 0.0016 | 4.1E-10 | COLEC12 | CETN1(-143.1kb)\|CLUL1(-159.7kb)\|COLEC12(0)\|THOC1(+169.2kb) | 1 |
| rs11564401 | 18 | 25666690 | A | G | -0.0175 | 0.0018 | 2.3E-21 | CDH2 | CDH2(0) | 0 |
| rs61735998 | 18 | 34289285 | T | G | -0.0433 | 0.0052 | 4.8E-17 | FHOD3 | FHOD3(0)\|KIAA1328(-119.8kb)\|TPGS2(-70.7kb) | 1 |
| rs7239759 | 18 | 57044710 | T | C | -0.0098 | 0.0016 | 1.9E-09 | LMAN1 | CCBE1(-53.46kb)\|CPLX4(+58.83kb)\|GRP(+146.7kb)\|LMAN1(+18.2kb)\|RAX(+104.1kb) | 1 |
| rs630621 | 18 | 57156096 | T | C | 0.0095 | 0.0017 | 1.9E-08 | CCBE1 | CCBE1(0)\|CPLX4(+170.2kb)\|LMAN1(+129.6kb) | 1 |
| rs35429067 | 19 | 2545186 | A | C | -0.0177 | 0.0029 | 2E-09 | GNG7 | DIRAS1(-169.4kb)\|GADD45B(+66.93kb)\|GNG7(0)\|LMNB2(+88.22kb)\|MIR7108(+110.2kb)\|SLC39A3(- | 1 |

| | | | | | | | | | | |
|---|---|---|---|---|---|---|---|---|---|---|
| | | | | | | | | | 187.3kb)\|SPPL2B(+190.1kb)\|TIMM13(+117.3kb)\|TMPRSS9(+119.1kb) | |
| rs10164397 | 19 | 32036003 | T | C | -0.0145 | 0.0017 | 5E-17 | THEG5 | THEG5(-43.09kb)\|TSHZ3(+195.8kb) | 0 |
| rs73037060 | 19 | 38175632 | T | C | 0.0102 | 0.0017 | 5.8E-09 | ZNF781 | LOC644554(-132.4kb)\|LOC100631378(-138.7kb)\|LOC101927720(+177.6kb)\|WDR87(-199.8kb)\|ZFP30(+29.32kb)\|ZNF540(+70.55kb)\|ZNF570(+199.4kb)\|ZNF571(+89.94kb)\|ZNF571-AS1(+97.38kb)\|ZNF573(-53.57kb)\|ZNF607(-11.63kb)\|ZNF781(0)\|ZNF793(+141.4kb) | 1 |
| rs235768 | 20 | 6759115 | A | T | -0.0104 | 0.0016 | 1.7E-10 | BMP2 | BMP2(0)\|=MISSENSE | 1 |
| rs6057598 | 20 | 31168281 | C | G | 0.0122 | 0.0018 | 8.8E-12 | LOC149950 | ASXL1(+141.2kb)\|C20orf203(-51.14kb)\|COMMD7(-122.2kb)\|DNMT3B(-181.9kb)\|LOC149950(-6.999kb)\|LOC101929698(+60.19kb)\|NOL4L(0) | 0 |
| rs6063031 | 20 | 45522102 | A | G | 0.0099 | 0.0016 | 9.8E-10 | EYA2 | EYA2(-1.16kb)\|SLC2A10(+157.1kb) | 1 |
| rs3091459 | 20 | 45789666 | T | C | -0.0091 | 0.0016 | 2.2E-08 | EYA2 | EYA2(0)\|LOC100131496(-157.6kb)\|MIR3616(-5.942kb)\|ZMYND8(-48.19kb) | 1 |
| rs58926663 | 20 | 61714040 | T | C | 0.0195 | 0.0028 | 5.6E-12 | LOC63930 | ARFGAP1(-190.1kb)\|BHLHE23(+75.65kb)\|BIRC7(-153.2kb)\|DIDO1(+144.7kb)\|FLJ16779(-171.3kb)\|GID8(+134.2kb)\|HAR1A(-18.6kb)\|HAR1B(-12.8kb)\|LINC00029(+45.66kb)\|LINC01056(+28.82kb)\|LOC63930(0)\|MIR124-3(-95.81kb)\|MIR3196(-156.1kb)\|NKAIN4(-158.1kb)\|SLC17A9(+114.1kb)\|YTHDF1(-112.7kb) | 1 |
| rs2223744 | 21 | 29503716 | T | C | -0.012 | 0.002 | 2.5E-09 | NCRNA00314 | LINC00314(+108.2kb) | 1 |
| rs74848904 | 21 | 36612241 | A | C | 0.0174 | 0.0029 | 2.8E-09 | RUNX1 | LOC100506403(-132.6kb)\|RUNX1(+190.6kb) | 1 |
| rs9984958 | 21 | 39908935 | A | T | 0.01 | 0.0016 | 1.3E-09 | ERG | ERG(0)\|LOC102724678(+190.9kb) | 1 |
| rs5754100 | 22 | 21916166 | T | C | -0.0138 | 0.002 | 4.5E-12 | UBE2L3 | CCDC116(-70.92kb)\|HIC2(+110.4kb)\|MAPK1(-197.8kb)\|MIR130B(-91.43kb)\|MIR301B(-91.1kb)\|PI4KAP2(+44.39kb)\|PPIL2(-104.1kb)\|RIMBP3B(+10.79kb)\|RIMBP3C(+10.42kb)\|SDF2L1(-80.38kb)\|TMEM191C(+91.94kb)\|UBE2L3(0)\|YDJC(-66.21kb)\|YPEL1(-135.7kb)\|=missense | 0 |
| rs6005752 | 22 | 28676252 | C | G | -0.0097 | 0.0018 | 4.8E-08 | TTC28 | MIR5739(-179.6kb)\|TTC28(0) | 0 |
| rs134545 | 22 | 28799080 | T | C | 0.0271 | 0.0017 | 2.1E-58 | TTC28 | MIR5739(-56.78kb)\|TTC28(0) | 0 |
| rs59032073 | 22 | 37910715 | A | G | 0.0353 | 0.0027 | 2.9E-39 | CARD10 | CARD10(0)\|CDC42EP1(-45.76kb)\|CYTH4(+199.3kb)\|ELFN2(+87.21kb)\|GGA1(-93.76kb)\|LGALS1(-160.9kb)\|LGALS2(-55.54kb)\|LOC100506271(+159.8kb)\|LOC101927051(-127.1kb)\|MFNG(+28.24kb)\|NOL12(-171.6kb)\|PDXP(-144kb)\|SH3BP1(-125kb)\|TRIOBP(-182.3kb) | 0 |
| rs5750495 | 22 | 38179120 | A | T | -0.0138 | 0.0017 | 3.1E-16 | TRIOBP | ANKRD54(-47.74kb)\|C22orf23(-159.9kb)\|EIF3L(-66.26kb)\|GALR3(- | 1 |

| | | | | | | | | | | |
|---|---|---|---|---|---|---|---|---|---|---|
| | | | | | | | | | 40.27kb)\|GCAT(-24.79kb)\|GGA1(+149.5kb)\|H1F0(-21.99kb)\|LGALS1(+103.3kb)\|LOC101927051(+124.7kb)\|MICALL1(-123kb)\|MIR658(-61.16kb)\|MIR659(-64.56kb)\|MIR6820(-184.4kb)\|NOL12(+89.64kb)\|PDXP(+116.2kb)\|POLR2F(-170.6kb)\|SH3BP1(+127.1kb)\|SOX10(-189.2kb)\|TRIOBP(+6.557kb)\|=frameshift\|=missense | |
| rs80557 | 22 | 46157281 | T | C | 0.0109 | 0.0018 | 7.6E-10 | ATXN10 | ATXN10(0)\|FBLN1(+160.3kb)\|MIR4762(+0.803kb)\|WNT7B(-159kb) | 1 |
| rs28541591 | 22 | 46300220 | A | C | -0.0203 | 0.0017 | 3.4E-33 | WNT7B | ATXN10(+59.03kb)\|LINC00899(-135.6kb)\|LOC150381(-149.5kb)\|LOC730668(-102.3kb)\|MIR3619(-186.7kb)\|MIR4762(+143.7kb)\|MIRLET7BHG(-181.7kb)\|PRR34(-146.1kb)\|WNT7B(-16.03kb) | 0 |
| rs9723267 | 22 | 46365557 | T | G | -0.018 | 0.0019 | 3.9E-22 | WNT7B | ATXN10(+124.4kb)\|LINC00899(-70.23kb)\|LOC150381(-84.17kb)\|LOC730668(-36.94kb)\|MIR3619(-121.4kb)\|MIR4763(-143.9kb)\|MIRLET7A3(-143.1kb)\|MIRLET7B(-144kb)\|MIRLET7BHG(-116.3kb)\|PPARA(-180.9kb)\|PRR34(-80.78kb)\|WNT7B(0) | 0 |
| rs6609798 | 23 | 48691511 | C | G | -0.0098 | 0.0018 | 3.2E-08 | PCSK1N | ERAS(+3.232kb)\|GATA1(+38.79kb)\|GLOD5(+59.45kb)\|GRIPAP1(-138.6kb)\|HDAC6(+8.131kb)\|KCND1(-127.1kb)\|OTUD5(-87.79kb)\|PCSK1N(0)\|PIM2(-78.95kb)\|PQBP1(-63.68kb)\|SLC35A2(-68.94kb)\|SUV39H1(+124.1kb)\|TFE3(-194.7kb)\|TIMM17B(-59.22kb)\|WAS(+141.7kb) | 1 |
| rs17250985 | 23 | 55831715 | T | C | -0.0339 | 0.0062 | 3.8E-08 | RRAGB | FOXR2(+179.1kb)\|RRAGB(+46.51kb) | 1 |
| rs626840 | 23 | 68057932 | A | G | -0.0231 | 0.0017 | 1.7E-41 | EFNB1 | EFNB1(0)\|STARD8(+112.2kb) | 0 |
| rs113485657 | 23 | 136508373 | A | C | 0.0244 | 0.0034 | 7.6E-13 | ZIC3 | ZIC3(-140kb) | 0 |

**Table S4. Cross population genetic effect correlation for optic nerve head parameters.**

| Traits | European - Asian[1] | European - African |
|---|---|---|
| VCDR (adjusted for VDD) | pge = 0.80<br>SE = 0.17<br>P = 0.22 | pge = 0.46<br>SE = 0.27<br>P = 0.047 |
| VDD | pge = 1.27<br>SE = 0.32<br>P = 0.40 | pge = 1.20<br>SE = 2.29<br>P = 0.93 |

[1]The genetic effect correlation was calculated using the “Popcorn” package.[3]

Pge, genetic effect correlation; SE, standard error; P value for the test that the genetic correlation is less than 1.

The meta-analysis of UKB and CLSA summary statistics was used as “European” VCDR/VDD. The VCDR/VDD GWAS summary statistics for Asian ancestry were from IGGC.[4] The VCDR/VDD GWAS summary statistics for African ancestry were from UKB (N = 2245). The GWAS sample size in Afrian population is very small, therefore, the results here should be interpreted with caution.

**Table S5. Pathway analysis of VDD-adjusted VCDR (list of 65 significant pathways after Bonferroni correction)**

| FULL_NAME | NGENES | BETA | BETA_STD | SE | P |
|---|---|---|---|---|---|
| Curated_gene_sets:reactome_elastic_fibre_formation | 46 | 1.0184 | 0.049136 | 0.15655 | 3.9718E-11 |
| Curated_gene_sets:reactome_molecules_associated_with_elastic_fibres | 39 | 1.0793 | 0.047956 | 0.16868 | 8.0324E-11 |
| GO_bp:go_negative_regulation_of_chondrocyte_differentiation | 20 | 1.7443 | 0.055527 | 0.27374 | 9.5529E-11 |
| GO_bp:go_telencephalon_regionalization | 13 | 2.0828 | 0.053465 | 0.34676 | 9.6612E-10 |
| GO_bp:go_skeletal_system_development | 512 | 0.27552 | 0.043819 | 0.04844 | 6.5304E-09 |
| GO_mf:go_extracellular_matrix_structural_constituent | 161 | 0.49614 | 0.044651 | 0.088377 | 1.0038E-08 |
| GO_bp:go_metanephros_development | 90 | 0.67734 | 0.045659 | 0.12104 | 1.1122E-08 |
| GO_bp:go_cell_differentiation_involved_in_metanephros_development | 26 | 1.2783 | 0.046388 | 0.22955 | 1.3029E-08 |
| GO_bp:go_tube_morphogenesis | 805 | 0.21088 | 0.041733 | 0.038687 | 2.5366E-08 |
| GO_bp:go_mesonephros_development | 101 | 0.58827 | 0.041997 | 0.10902 | 3.4545E-08 |
| GO_bp:go_renal_tubule_development | 94 | 0.61101 | 0.042089 | 0.11332 | 3.5302E-08 |
| GO_bp:go_mesenchymal_to_epithelial_transition_involved_in_metanephros_morphogenesis | 13 | 1.8113 | 0.046496 | 0.33633 | 3.6555E-08 |
| GO_bp:go_kidney_morphogenesis | 94 | 0.60273 | 0.041518 | 0.11302 | 4.89E-08 |
| Curated_gene_sets:reactome_extracellular_matrix_organization | 296 | 0.34406 | 0.04184 | 0.064705 | 5.3241E-08 |
| GO_cc:go_extracellular_matrix | 513 | 0.25233 | 0.040169 | 0.047806 | 6.6E-08 |
| GO_bp:go_metanephric_nephron_development | 41 | 0.94185 | 0.042906 | 0.17884 | 7.0286E-08 |
| GO_bp:go_embryo_development | 975 | 0.18666 | 0.040469 | 0.035743 | 8.9388E-08 |
| GO_bp:go_bone_development | 210 | 0.39147 | 0.040186 | 0.075071 | 9.3062E-08 |
| GO_bp:go_negative_regulation_of_cartilage_development | 25 | 1.2347 | 0.043939 | 0.23729 | 9.8904E-08 |
| GO_bp:go_regulation_of_metanephros_development | 23 | 1.3142 | 0.044862 | 0.25447 | 1.2174E-07 |
| Curated_gene_sets:charafe_breast_cancer_luminal_vs_mesenchymal_dn | 460 | 0.26454 | 0.039934 | 0.051228 | 1.2209E-07 |
| GO_bp:go_nephron_morphogenesis | 78 | 0.63704 | 0.039989 | 0.12364 | 1.3E-07 |
| Curated_gene_sets:reactome_collagen_formation | 88 | 0.61273 | 0.040844 | 0.12008 | 1.6916E-07 |
| GO_bp:go_kidney_epithelium_development | 137 | 0.48057 | 0.03992 | 0.094313 | 1.7566E-07 |
| GO_bp:go_chondrocyte_differentiation | 117 | 0.51134 | 0.039274 | 0.10062 | 1.8864E-07 |
| GO_bp:go_morphogenesis_of_a_branching_structure | 196 | 0.39843 | 0.039527 | 0.079034 | 2.3352E-07 |

| | | | | | |
|---|---|---|---|---|---|
| GO_bp:go_mesenchymal_to_epithelial_transition | 20 | 1.3429 | 0.042749 | 0.26666 | 2.4013E-07 |
| GO_bp:go_tube_development | 990 | 0.17492 | 0.0382 | 0.03484 | 2.5987E-07 |
| Curated_gene_sets:reactome_o_glycosylation_of_tsr_domain_containing_proteins | 37 | 0.94577 | 0.040933 | 0.18861 | 2.6859E-07 |
| GO_cc:go_collagen_containing_extracellular_matrix | 397 | 0.27259 | 0.03829 | 0.054467 | 2.8233E-07 |
| GO_bp:go_metanephric_nephron_tubule_epithelial_cell_differentiation | 7 | 2.2529 | 0.042442 | 0.45495 | 3.7074E-07 |
| GO_bp:go_animal_organ_morphogenesis | 1026 | 0.17085 | 0.037948 | 0.034529 | 3.7807E-07 |
| GO_bp:go_regulation_of_chondrocyte_differentiation | 47 | 0.84146 | 0.041035 | 0.17042 | 3.9899E-07 |
| GO_bp:go_s_shaped_body_morphogenesis | 8 | 2.1352 | 0.043001 | 0.43434 | 4.4584E-07 |
| Curated_gene_sets:biocarta_alk_pathway | 35 | 0.89915 | 0.03785 | 0.18464 | 5.6357E-07 |
| GO_cc:go_extracellular_matrix_component | 49 | 0.78923 | 0.039296 | 0.16238 | 5.9114E-07 |
| Curated_gene_sets:boquest_stem_cell_up | 260 | 0.34041 | 0.038833 | 0.070699 | 7.4211E-07 |
| GO_bp:go_mesonephric_tubule_morphogenesis | 66 | 0.64774 | 0.037414 | 0.13499 | 8.0551E-07 |
| GO_bp:go_sensory_organ_development | 531 | 0.22652 | 0.03667 | 0.04729 | 8.4042E-07 |
| Curated_gene_sets:naba_core_matrisome | 270 | 0.32375 | 0.037626 | 0.06775 | 8.8955E-07 |
| GO_bp:go_metanephric_s_shaped_body_morphogenesis | 6 | 2.3813 | 0.041536 | 0.49926 | 9.2938E-07 |
| GO_bp:go_cranial_skeletal_system_development | 67 | 0.64425 | 0.037492 | 0.13615 | 1.1208E-06 |
| Curated_gene_sets:kegg_focal_adhesion | 200 | 0.35466 | 0.035539 | 0.075135 | 1.1862E-06 |
| GO_bp:go_branch_elongation_involved_in_ureteric_bud_branching | 5 | 2.4776 | 0.03945 | 0.52698 | 1.3011E-06 |
| GO_bp:go_digestive_tract_morphogenesis | 49 | 0.7422 | 0.036955 | 0.15814 | 1.3534E-06 |
| GO_bp:go_embryonic_organ_development | 420 | 0.25475 | 0.036783 | 0.054286 | 1.3583E-06 |
| Curated_gene_sets:davicioni_molecular_arms_vs_erms_dn | 175 | 0.37723 | 0.035382 | 0.080402 | 1.3638E-06 |
| GO_bp:go_gland_development | 434 | 0.23942 | 0.035128 | 0.051263 | 1.5141E-06 |
| GO_bp:go_urogenital_system_development | 328 | 0.28251 | 0.036134 | 0.060613 | 1.5859E-06 |
| GO_bp:go_metanephric_renal_vesicle_morphogenesis | 16 | 1.4197 | 0.040428 | 0.30464 | 1.5895E-06 |
| Curated_gene_sets:reactome_assembly_of_collagen_fibrils_and_other_multimeric_structures | 61 | 0.68289 | 0.037925 | 0.14668 | 1.6273E-06 |
| Curated_gene_sets:kegg_basal_cell_carcinoma | 55 | 0.64126 | 0.033822 | 0.13833 | 1.7901E-06 |
| GO_bp:go_epithelial_tube_morphogenesis | 313 | 0.28295 | 0.035367 | 0.061334 | 1.9956E-06 |
| GO_bp:go_skeletal_system_morphogenesis | 236 | 0.33175 | 0.036078 | 0.072159 | 2.1531E-06 |
| GO_bp:go_metanephros_morphogenesis | 33 | 0.90988 | 0.037193 | 0.19834 | 2.2593E-06 |
| GO_mf:go_transmembrane_receptor_protein_serine_threonine | 13 | 1.5128 | 0.038832 | 0.33078 | 2.4158E-06 |

| | | | | | |
|---|---|---|---|---|---|
| _kinase_binding | | | | | |
| Curated_gene_sets:lindgren_bladder_cancer_high_recurrence | 49 | 0.73395 | 0.036544 | 0.16052 | 2.428E-06 |
| GO_bp:go_embryonic_cranial_skeleton_morphogenesis | 46 | 0.73304 | 0.035367 | 0.16062 | 2.5273E-06 |
| Curated_gene_sets:galie_tumor_stemness_genes | 6 | 1.983 | 0.034587 | 0.43557 | 2.6667E-06 |
| Curated_gene_sets:dacosta_uv_response_via_ercc3_dn | 854 | 0.17461 | 0.035544 | 0.038367 | 2.6867E-06 |
| GO_bp:go_kidney_mesenchyme_development | 19 | 1.2174 | 0.037774 | 0.26774 | 2.7391E-06 |
| GO_bp:go_morphogenesis_of_an_epithelium | 525 | 0.21301 | 0.034294 | 0.046904 | 2.8114E-06 |
| GO_bp:go_glial_cell_differentiation | 204 | 0.34017 | 0.034422 | 0.075126 | 2.9968E-06 |
| GO_bp:go_regulation_of_cartilage_development | 64 | 0.64825 | 0.036874 | 0.14353 | 3.1643E-06 |
| GO_bp:go_negative_regulation_of_cell_population_proliferation | 664 | 0.18484 | 0.033345 | 0.040954 | 3.2103E-06 |

**Table S6. Pathway analysis of VDD (list of 82 significant pathways after Bonferroni correction).**

| FULL_NAME | NGENES | BETA | BETA_STD | SE | P |
|---|---|---|---|---|---|
| GO_bp:go_kidney_epithelium_development | 137 | 0.59355 | 0.049495 | 0.087871 | 7.374E-12 |
| GO_bp:go_tube_morphogenesis | 806 | 0.24186 | 0.04807 | 0.036085 | 1.0547E-11 |
| GO_bp:go_tube_development | 991 | 0.21628 | 0.047428 | 0.032551 | 1.5668E-11 |
| GO_bp:go_nephron_epithelium_development | 106 | 0.63291 | 0.046461 | 0.099097 | 8.6797E-11 |
| GO_bp:go_renal_tubule_development | 94 | 0.67056 | 0.046369 | 0.10667 | 1.6619E-10 |
| GO_bp:go_animal_organ_morphogenesis | 1026 | 0.20304 | 0.045261 | 0.032385 | 1.8544E-10 |
| GO_bp:go_telencephalon_regionalization | 13 | 1.9534 | 0.050336 | 0.33041 | 1.7212E-09 |
| GO_bp:go_cell_fate_commitment | 249 | 0.39026 | 0.043747 | 0.06705 | 2.9857E-09 |
| GO_bp:go_artery_development | 83 | 0.65014 | 0.042257 | 0.11194 | 3.221E-09 |
| GO_bp:go_mesonephros_development | 101 | 0.58775 | 0.042121 | 0.10159 | 3.6734E-09 |
| GO_bp:go_tube_formation | 144 | 0.5023 | 0.042935 | 0.08735 | 4.5252E-09 |
| GO_bp:go_epithelial_tube_morphogenesis | 313 | 0.33069 | 0.041492 | 0.058093 | 6.361E-09 |
| GO_bp:go_epithelium_development | 1241 | 0.17282 | 0.042123 | 0.030659 | 8.7977E-09 |
| GO_bp:go_ossification | 368 | 0.29617 | 0.040236 | 0.052556 | 8.8689E-09 |
| GO_bp:go_nephron_development | 138 | 0.49571 | 0.041486 | 0.088275 | 9.9462E-09 |
| GO_bp:go_forebrain_regionalization | 24 | 1.2834 | 0.044925 | 0.22921 | 1.0919E-08 |

| | | | | | |
|---|---|---|---|---|---|
| GO_bp:go_aorta_development | 52 | 0.79416 | 0.040888 | 0.14289 | 1.3847E-08 |
| GO_bp:go_morphogenesis_of_an_epithelium | 524 | 0.24491 | 0.039541 | 0.044294 | 1.6313E-08 |
| GO_bp:go_regionalization | 339 | 0.31869 | 0.041586 | 0.057648 | 1.641E-08 |
| GO_bp:go_urogenital_system_development | 328 | 0.31432 | 0.040356 | 0.057085 | 1.8598E-08 |
| GO_bp:go_kidney_morphogenesis | 94 | 0.58193 | 0.04024 | 0.1057 | 1.8671E-08 |
| GO_bp:go_outflow_tract_morphogenesis | 75 | 0.6681 | 0.041287 | 0.1224 | 2.4345E-08 |
| GO_bp:go_nephron_morphogenesis | 78 | 0.62891 | 0.039631 | 0.11586 | 2.8812E-08 |
| GO_bp:go_skeletal_system_development | 512 | 0.24449 | 0.039031 | 0.045328 | 3.4948E-08 |
| GO_bp:go_mesonephric_tubule_morphogenesis | 66 | 0.67782 | 0.039303 | 0.12675 | 4.5101E-08 |
| GO_bp:go_circulatory_system_development | 1023 | 0.17148 | 0.038174 | 0.032085 | 4.5857E-08 |
| GO_bp:go_anatomical_structure_formation_involved_in_morphogenesis | 1060 | 0.16674 | 0.037746 | 0.031407 | 5.5783E-08 |
| GO_bp:go_cardiac_septum_development | 105 | 0.53891 | 0.039375 | 0.10162 | 5.752E-08 |
| GO_bp:go_heart_morphogenesis | 247 | 0.34719 | 0.038763 | 0.065955 | 7.1311E-08 |
| GO_bp:go_pattern_specification_process | 433 | 0.26691 | 0.039266 | 0.050738 | 7.2682E-08 |
| GO_bp:go_osteoblast_differentiation | 200 | 0.36088 | 0.036301 | 0.068925 | 8.3087E-08 |
| GO_bp:go_cell_differentiation_involved_in_kidney_development | 53 | 0.76568 | 0.039798 | 0.14625 | 8.3262E-08 |
| GO_bp:go_tissue_morphogenesis | 651 | 0.20867 | 0.037426 | 0.039863 | 8.3593E-08 |
| GO_bp:go_renal_system_development | 290 | 0.31541 | 0.038116 | 0.060725 | 1.0403E-07 |
| Curated_gene_sets:kinsey_targets_of_ewsr1_flii_fusion_dn | 320 | 0.27854 | 0.03533 | 0.053709 | 1.0862E-07 |
| GO_bp:go_positive_regulation_of_animal_organ_morphogenesis | 81 | 0.6029 | 0.038713 | 0.11696 | 1.2843E-07 |
| GO_bp:go_mesenchyme_development | 258 | 0.33025 | 0.037674 | 0.064153 | 1.3315E-07 |
| GO_bp:go_regulation_of_ossification | 179 | 0.3962 | 0.037724 | 0.076965 | 1.3319E-07 |
| GO_bp:go_embryonic_morphogenesis | 567 | 0.22404 | 0.037584 | 0.043747 | 1.534E-07 |
| GO_bp:go_regulation_of_heart_morphogenesis | 37 | 0.86145 | 0.037428 | 0.16866 | 1.649E-07 |
| GO_bp:go_embryo_development | 973 | 0.17015 | 0.036989 | 0.033411 | 1.7854E-07 |
| GO_bp:go_cardiovascular_system_development | 688 | 0.1968 | 0.036251 | 0.038841 | 2.0426E-07 |
| GO_bp:go_heart_development | 533 | 0.22042 | 0.035883 | 0.043934 | 2.649E-07 |
| Curated_gene_sets:kegg_basal_cell_carcinoma | 55 | 0.65895 | 0.03489 | 0.13157 | 2.7725E-07 |
| GO_bp:go_appendage_morphogenesis | 146 | 0.42903 | 0.036924 | 0.085753 | 2.8491E-07 |
| GO_bp:go_cardiac_chamber_development | 163 | 0.4008 | 0.036431 | 0.080126 | 2.8635E-07 |

| | | | | | |
|---|---|---|---|---|---|
| GO_bp:go_mesenchymal_cell_proliferation | 45 | 0.82325 | 0.039438 | 0.16615 | 3.6505E-07 |
| GO_bp:go_cell_cell_signaling_by_wnt | 504 | 0.21884 | 0.034669 | 0.0442 | 3.7252E-07 |
| GO_bp:go_cell_surface_receptor_signaling_pathway_involved_in_cell_cell_signaling | 604 | 0.20091 | 0.034753 | 0.040614 | 3.8064E-07 |
| GO_bp:go_metanephros_development | 90 | 0.53229 | 0.03602 | 0.10886 | 5.0992E-07 |
| GO_bp:go_regulation_of_muscle_organ_development | 133 | 0.40852 | 0.033569 | 0.084165 | 6.1062E-07 |
| GO_bp:go_mesenchymal_cell_differentiation | 206 | 0.34268 | 0.034978 | 0.070843 | 6.6421E-07 |
| Curated_gene_sets:gryder_pax3foxo1_enhancers_in_tads | 1009 | 0.15954 | 0.035284 | 0.03305 | 6.9823E-07 |
| GO_mf:go_bmp_receptor_binding | 8 | 1.8621 | 0.037647 | 0.38583 | 7.0173E-07 |
| GO_bp:go_striated_muscle_cell_proliferation | 62 | 0.6353 | 0.035707 | 0.13206 | 7.5827E-07 |
| GO_bp:go_regulation_of_epithelial_cell_migration | 213 | 0.33158 | 0.034409 | 0.069807 | 1.0253E-06 |
| GO_bp:go_metanephric_nephron_development | 41 | 0.79629 | 0.036415 | 0.16799 | 1.0769E-06 |
| GO_bp:go_regulation_of_osteoblast_differentiation | 109 | 0.45851 | 0.034128 | 0.09678 | 1.0898E-06 |
| GO_bp:go_negative_regulation_of_cartilage_development | 25 | 0.95042 | 0.033953 | 0.20081 | 1.1146E-06 |
| GO_bp:go_heart_field_specification | 15 | 1.3062 | 0.036155 | 0.27628 | 1.1427E-06 |
| GO_bp:go_negative_regulation_of_developmental_process | 905 | 0.15724 | 0.033027 | 0.033283 | 1.1633E-06 |
| GO_bp:go_tongue_development | 20 | 1.0353 | 0.033084 | 0.21953 | 1.2135E-06 |
| GO_bp:go_epithelial_cell_differentiation_involved_in_kidney_development | 43 | 0.77001 | 0.03606 | 0.16476 | 1.4915E-06 |
| Curated_gene_sets:kegg_hedgehog_signaling_pathway | 56 | 0.61972 | 0.033108 | 0.13275 | 1.5288E-06 |
| GO_bp:go_metanephros_morphogenesis | 33 | 0.85774 | 0.035198 | 0.18387 | 1.5546E-06 |
| GO_bp:go_animal_organ_formation | 63 | 0.58579 | 0.033188 | 0.1259 | 1.6479E-06 |
| GO_bp:go_cardiac_ventricle_development | 122 | 0.43101 | 0.03393 | 0.092908 | 1.7621E-06 |
| GO_bp:go_outflow_tract_septum_morphogenesis | 27 | 0.99332 | 0.036876 | 0.2142 | 1.7764E-06 |
| GO_bp:go_cardiac_septum_morphogenesis | 72 | 0.58578 | 0.035471 | 0.12641 | 1.8072E-06 |
| GO_bp:go_cell_differentiation_involved_in_metanephros_development | 26 | 0.99946 | 0.036411 | 0.21584 | 1.8358E-06 |
| GO_bp:go_positive_regulation_of_rna_biosynthetic_process | 1568 | 0.11801 | 0.032042 | 0.025502 | 1.8643E-06 |
| GO_bp:go_ventricular_septum_development | 68 | 0.58756 | 0.03458 | 0.12754 | 2.0585E-06 |
| GO_bp:go_aorta_morphogenesis | 30 | 0.87871 | 0.034383 | 0.19107 | 2.1398E-06 |
| GO_bp:go_specification_of_animal_organ_identity | 34 | 0.81227 | 0.033833 | 0.17715 | 2.2824E-06 |
| GO_bp:go_positive_regulation_of_biosynthetic_process | 1928 | 0.10568 | 0.031497 | 0.023075 | 2.3447E-06 |
| Curated_gene_sets:reactome_elastic_fibre_formation | 46 | 0.65429 | 0.031689 | 0.14293 | 2.366E-06 |

| | | | | | |
|---|---|---|---|---|---|
| GO_bp:go_renal_vesicle_development | 20 | 1.158 | 0.037006 | 0.2533 | 2.4356E-06 |
| GO_bp:go_sensory_organ_development | 531 | 0.20272 | 0.032942 | 0.044422 | 2.5313E-06 |
| GO_mf:go_transmembrane_receptor_protein_serine_threonine_kinase_activity | 17 | 1.1058 | 0.032582 | 0.2425 | 2.5748E-06 |
| GO_bp:go_canonical_wnt_signaling_pathway | 320 | 0.2536 | 0.032168 | 0.055716 | 2.6784E-06 |
| Curated_gene_sets:reactome_extracellular_matrix_organization | 296 | 0.2674 | 0.032641 | 0.058757 | 2.6891E-06 |
| GO_bp:go_commitment_of_neuronal_cell_to_specific_neuron_type_in_forebrain | 7 | 1.9593 | 0.037055 | 0.43063 | 2.7014E-06 |

**Table S7. Training specifics of Convolutional Neural Network algorithms.**

| Model | Architecture | LR | Batch Size | Data Augmentations |
|---|---|---|---|---|
| VCDR | ResNet-34 | Cyclic (10e-03, 10e-04) | 32 | Rotation (max 10 deg)<br>Symmetric warp (max 0.2)<br>Zoom (max scale= 1.1)<br>Brightness (max 15%)<br>Contrast (max 15%)<br>Horizontal flip |
| VDD | ResNet-34 | Cyclic (10e-03, 10e-04) | 32 | Rotation (max 10 deg)<br>Brightness (max 15%)<br>Contrast (max 15%<br>Horizontal flip |
| Gradeability | ResNet-34 | Cyclic (10e-03, 10e-04) | 32 | Rotation (max 10 deg)<br>Symmetric warp (max 0.2)<br>Zoom (max scale= 1.1)<br>Brightness (max 15%)<br>Contrast (max 15%)<br>Horizontal flip |

**Table S8. Summary of study datasets.**

| Study | Description | Refs |
|---|---|---|
| UKB VCDR/VDD GWAS | The sample size is 68,240 for VCDR/VDD GWAS in UKB participants of white ancestry. Using a K-means clustering method to confirm genetic ancestry, we also identified participants in other ancestry groups for African (N = 2245), South Asian (N = 2100) and East Asian (N = ~500) ancestry. We performed GWAS in African and South Asian participants ancestry using PLINK. | - |
| CLSA VCDR/VDD GWAS | The sample size is 18,304 for VCDR/VDD GWAS in CLSA participants of European ancestry. We didn't perform GWAS in other ethinic groups in CLSA due to the very small sample size (N <300). | - |
| IGGC VCDR/disc area | The GWAS summary statistics were downloaded for individuals of European descent ($N_{VCDR}$= 25,180, $N_{disc}$= 24,509, from the latest HRC imputation), as well as Asian descent ($N_{VCDR}$= 8,373, $N_{disc}$= 7,307). | [4,5] |
| Glaucoma GWAS datasets | The glaucoma datasets were used to look up AI-based GWAS new findings, including 7,947 glaucoma cases and 119,318 controls from UK Biobank and 3,071 POAG cases and 6,750 historic controls from the Australian & New Zealand Registry of Advanced Glaucoma (ANZRAG) study. | [6] |

CLSA, the Canadian Longitudinal Study on Aging; GWAS, genome-wide association study; IGGC, the International Glaucoma Genetic Consortium; UKB, UK Biobank; VCDR, vertical cup-to-disc ratio; VDD, vertical disc diameter.
